# Syntaxin11 Deficiency Inhibits CRAC Channel Priming to Suppress Cytotoxicity and Gene Expression in T Lymphocytes Independent of Membrane Trafficking

**DOI:** 10.1101/2024.10.25.620144

**Authors:** Sritama Datta, Abhikarsh Gupta, Kunal Mukesh Jagetiya, Resmi Bera, Vikas Tiwari, Megumi Yamashita, Atharva Rahul Yande, Abdul Rishad, Vishal Malik, Sreejith Raran-Kurussi, Sandra Ammann, Mohammad Shahrooei, Kalyaneswar Mandal, Ramanathan Sowdhamini, Murali Prakriya, Adish Dani, Monika Vig

**Author notes:** These authors contributed equally (Sritama Datta, Abhikarsh Gupta, Kunal Mukesh Jagetiya, Resmi Bera, Vikas Tiwari, Megumi Yamashita).

## Abstract

Mutations in Syntaxin11, a Q-SNARE, result in a fatal immune disorder known as familial hemophagocytic lymphohistiocytosis 4 (FHLH4) in human patients. A key diagnostic feature of FHLH4 is defective T and natural killer (NK) cell cytotoxicity. Here we show that Syntaxin11 directly binds and regulates Orai1, the pore forming subunit of calcium release activated calcium (CRAC) channels. CRAC channels enable store-operated calcium entry (SOCE) from the extracellular space and are crucial for granule exocytosis and nuclear factor of activated T cell (NFAT) dependent gene expression in activated lymphocytes. Syntaxin11 depletion strongly inhibited SOCE, CRAC currents, NFAT activation, interleukin-2 gene expression and degranulation in FHLH4 patient T lymphocytes and cell lines without affecting membrane trafficking. Remarkably, defects of cytolytic granule exocytosis as well as interleukin-2 expression could be reversed by ionomycin in patient T lymphocytes and a constitutively active, H134S, mutant of Orai1 rescued calcium entry in Syntaxin11 depleted cells. Further analyses showed that Syntaxin11 ‘primes’ Orai1 for optimal on-site multimeric assembly which was Stim independent but required for gating. Priming of ion channel pore subunits is, therefore, a primary function of specific SNAREs which may have preceded their role in membrane trafficking and vesicle fusion.

## Introduction

Familial hemophagocytic lymphohistiocytosis 4 (FHLH4) is a life-threatening immune disease caused due to mutations of STX11 [1]. The underlying cause of FHLH is reduced cytolytic activity of T and NK cells [2], which renders patients susceptible to recurrent infections. Patients suffer from high fever, severe lymphopenia in early infancy and succumb to disease by adolescence, unless given bone marrow transplants. STX11 is highly expressed in almost all cells of the immune system and, being a Q-SNARE, is associated with exocytosis of vesicles, by default. Intriguingly, FHLH4 patients also manifest cytokine storm, which cannot be reconciled with a presumed general defect in membrane trafficking and vesicle fusion.

Most eukaryotic cells have a limited amount of calcium sequestered inside intracellular stores, the largest of which is the endoplasmic reticulum (ER). Signaling from cell surface receptors induces the release of stored calcium which, in turn, activates store-operated calcium entry (SOCE) to replenish stores, sustain signaling and drive other calcium dependent cellular processes [3]. Calcium release activated calcium (CRAC) channels are the major conductors of SOCE [4]. In lymphocytes and mast cells, CRAC currents have been shown to be crucial for cytotoxicity, degranulation as well as gene expression [5, 6]. Working in concert with a number of accessory proteins, Orai (CRACM) multimers conduct SOCE upon functional clustering with Stim proteins in ER-PM junctions [7–9]. Yet, the prevailing thought attributes all structural transitions that result in the activation of CRAC currents to Stim proteins [10].

Genome-wide RNAi screens, which initially identified Orai (CRACM), have also yielded a wealth of information regarding additional crucial players in SOCE [8, 11, 12]. For instance, we have previously shown that alpha-soluble N-ethylmaleimide sensitive factor (NSF) attachment protein (α-SNAP), a well-known synaptic family adaptor protein that forms a part of the 20S SNARE super-complex [13], is a crucial component of the CRAC channel supramolecular complex [8, 9]. α-SNAP independently binds Stim as well as Orai and is required for the correct assembly and calcium selectivity of CRAC channels [14–16]. α-SNAP depletion revealed that Stim:Orai coupling is necessary but not sufficient for SOCE, a process that requires distinct time-resolved molecular steps *in vivo*.

In addition to α-SNAP, SNAREs also came up as candidates in two genome-wide RNAi screens for regulators of SOCE but were not characterized [8, 12]. Furthermore, specific SNAREs have been previously found to form inhibitory associations with specific ion channels, but their exact role was not defined [17]. Therefore, we hypothesized a more fundamental function for the association of SNAREs with ion channel pore subunits and set out to test this in the context of SOCE. Using a targeted RNAi screen, biochemical and biophysical analyses, we have found a direct role of STX11 in facilitating the assembly of CRAC channel pore subunit, Orai1, into optimal multimers. Assembly of Orai1 was independent of, and preceded, trapping and gating by Stim1. Analyses of T cells derived from symptomatic STX11 deficient, FHLH4 patient showed that the previously reported [2], as well as newly identified, defects in degranulation, interleukin-2 gene expression predominantly stem from suppressed SOCE and not due to a general defect in vesicle trafficking or reduced levels of Orai1 in PM. Based on the findings presented here, we demonstrate a novel SNARE dependent priming step for CRAC channel pore in the molecular sequence of SOCE and suggest preventive pharmacological interventions for FHLH4 patients.

## Results

RNAi dependent ablation of a Q-SNARE, Syntaxin 5 (STX5), has earlier shown inhibition of SOCE in a genome-wide screen performed in *Drosophila* S2 cells [12] and STX1A has been previously shown to bind a variety of ion channels [17]. Therefore, we first knocked down STX5 (Figure 1-figure supplement 1A) and STX1A (Figure 1-figure supplement 1B) using different sequences of shRNA targeting each gene in HEK293 cells and measured SOCE in response to Thapsigargin (TG). In both cases, we did not see a significant defect in SOCE. Since several proteins involved in synaptic function are not ubiquitously expressed and many show redundancy in function, we searched online databases to target SNAREs that are expressed in HEK293 and/or T cells (Figure 1-figure supplement 1C-H). We found that Syntaxin11 knockdown showed strong and consistent inhibition of SOCE by day 3 to 4 in HEK293 (Figure 1-figure supplement 2A-B) and Jurkat T cells (Figure 1A-B). Quantitative PCR (qPCR) analysis of total RNA extracted from scramble (scr) and STX11 shRNA treated cells showed ∼60-70% depletion of STX11 mRNA in both the cell lines (Figure 1-figure supplement 2C-D) and ectopic expression of STX11 in STX11 depleted Jurkat cells rescued SOCE (Figure 1C, D). The efficiency of STX11 depletion was assessed by qPCR for each experiment reported in this study. We restricted our analysis to day 3 to 4, post RNAi treatment, as longer incubations with RNAi or CRISPR knockout of STX11 affected cell viability.

**Figure 1:**
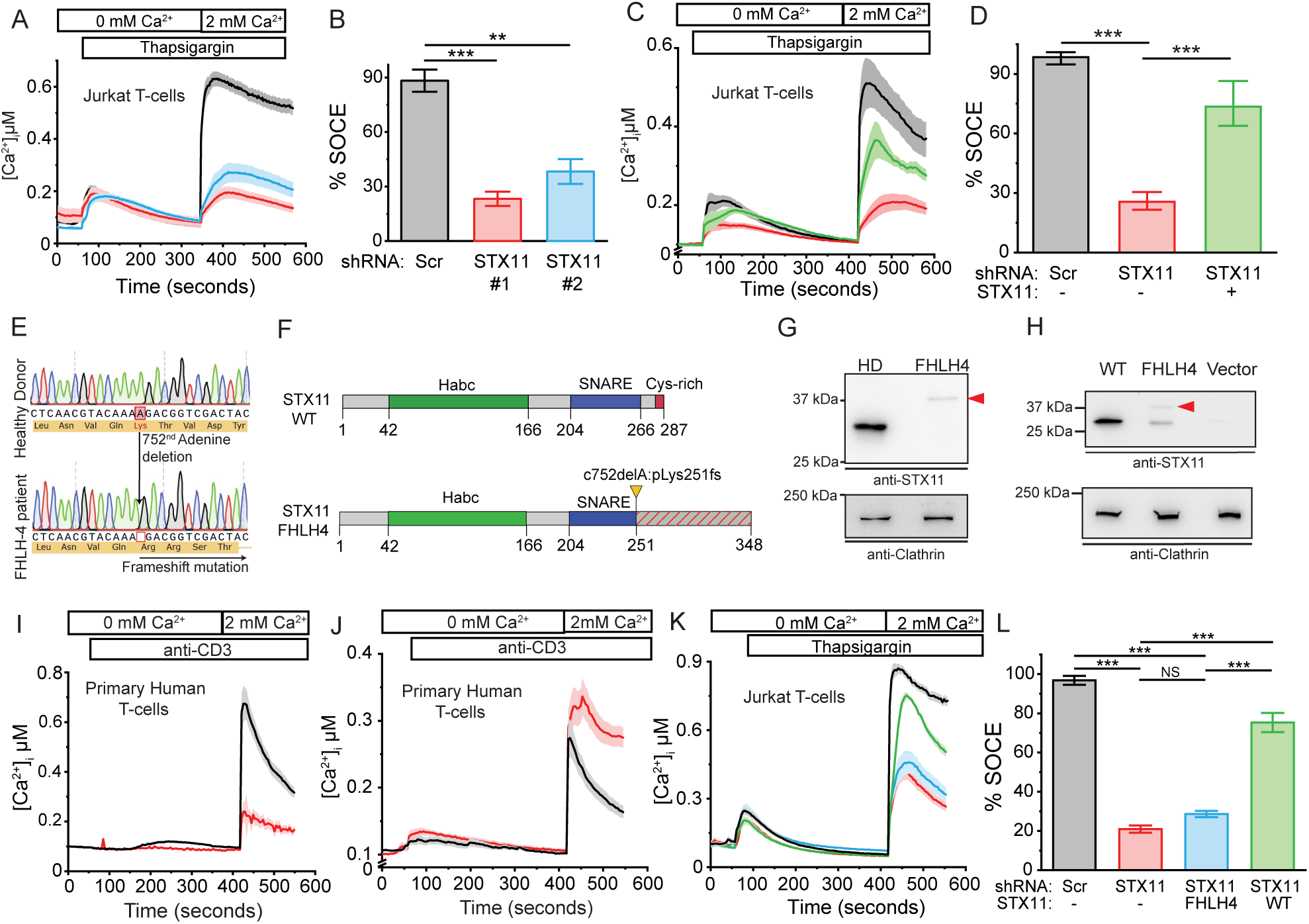
STX11 is required for SOCE. Measurement of SOCE in STX11 depleted Jurkat and FHLH4 patient T cells. (**A-B**) Representative Fura-2 calcium imaging assay (**A**) and quantification of independent repeats (**B**) measuring thapsigargin (TG) induced SOCE in Jurkat T cells treated with Scr (black), STX11#1 (red) or STX11#2 (blue) shRNAs. (n=30-40 cells/group, N=3). (**C-D**) Representative Fura-2 calcium imaging assay (**C**) and quantification of repeats (**D**) showing reconstitution of SOCE in STX11 depleted Jurkat T cells by ectopic expression of STX11. Scr shRNA with empty vector (EV) expression (black), STX11 shRNA with EV expression (red), STX11 shRNA with STX11 expression (green). (n=30-40 cells/group, N=3). (**E**) Sanger sequencing of FHLH4 patient DNA showing deletion of a single Adenine at the 752^nd^ position and the resulting frameshift. (**F**) Schematic showing the predicted domain distribution of the wildtype *versus* FHLH4 mutant STX11 protein. (**G**) Western blot on WCLs of healthy donor (HD) and FHLH4 patient PBMCs showing the relative molecular weight and abundance of the wildtype and mutant STX11 bands. (**H**) Western blot on WCLs of HEK293 cells over-expressing wildtype or FHLH4 mutant STX11 and empty vector. (**I**) Fura-2 calcium imaging assay measuring anti-CD3 induced SOCE in HD (black) and FHLH4 (red) T cells. (n=30-40 cells/group). (**J**) Fura-2 calcium imaging assay measuring SOCE in HD expressing empty vector (EV) (black) and FHLH4 T cells expressing human STX11 (red), respectively. (n=30-40 cells/group). (**K-L**) Representative Fura-2 calcium imaging assay (**K**) and quantification of repeats (**L**) measuring SOCE in Jurkat T cells treated either with Scr shRNA expressing empty vector (EV) (black), STX11 shRNA, expressing EV (red), STX11 shRNA, expressing wildtype STX11 (green), STX11 shRNA, expressing FHLH4 mutant STX11 (blue).

Given that STX11 is highly expressed in most immune cells and is the only known SNARE that, when mutated, causes defects in T and NK cell cytotoxicity in humans and mice [2], we hypothesized that reduced SOCE contributes to the cytotoxicity defects of FHLH4 patient T cells. To test this hypothesis, we isolated PBMCs from an FHLH4 patient with a homozygous deletion frameshift mutation in the STX11 coding sequence, c.752delA:p.Lys251fs. The patient presented with all symptoms of FHLH4 disease by 4 years of age. Given the limited sample amount, we expanded PBMC numbers by periodically stimulating patient and healthy donor cells with PHA and IL-2, prior to analysis. Isolation and Sanger sequencing of DNA from patient PBMCs showed deletion of a single Adenine at position 752 in the STX11 gene which lead to frameshift following Lysine 251 (Figure 1E). The absence of any contaminating wildtype peaks showed that the mutation was present in all blood cells. Figure 1F shows schematic of the predicted mutant STX11 protein with an estimated molecular weight of ∼39.5 KDa compared to ∼33 KDa of wildtype STX11. The frameshift mutation resulted in disruption of the SNARE domain, ablation of the terminal cysteines and elongation of the resulting transcript. We subjected whole cell lysates prepared from the FHLH4 patient and healthy donor (HD) PBMCs to SDS-PAGE and Western blot and found that while the wildtype STX11 band was absent, a faint but distinct band running at ∼37 KDa was present in the FHLH4 patient sample lysate (Figure 1G). Therefore, the predicted and observed molecular weight of the FHLH4 mutant protein was higher but its levels appeared significantly reduced compared to STX11 from HD. This is likely due to protein instability and degradation because ectopic expression of the FHLH4 STX11 mutant in HEK293 cells showed at least two clear bands (Figure 1H), although reduced reactivity to anti-STX11 antibody and premature termination of translation are also possible. Interestingly, all prior FHLH4 STX11 patient mutations that have been biochemically characterized, till date, also showed severe depletion of STX11 protein levels in patient T and NK cells [18–21]. Therefore, the frame shift mutation that has been characterized in our study is representative of most other STX11 human mutations, characterized earlier, in terms of the mechanisms underlying the observed phenotypes. In accordance with our findings in the STX11 depleted cell lines (Figure 1A-B), SOCE was found to be significantly defective in FHLH4 T cells when compared to HD (Figure 1I). Further, expression of wildtype (Figure 1J), but not FHLH4 mutant STX11 (Figure 1K-L), reversed defective SOCE in STX11 deficient patient (Figure 1J) and Jurkat T cells (Figure 1K-L), respectively. These data suggest that, in addition to being unstable, FHLH4 mutant STX11 was functionally compromised and no additional unknown mutations resulted in reduced SOCE.

To determine whether the inhibition of SOCE in Figure 1 resulted from changes in the activity of CRAC channels or their PM expression levels, we quantified the total as well as plasma membrane levels of Orai1 in different cell lines (Figure 2). We generated a stable HEK293 cell line by expressing a previously characterized Orai1 fusion construct containing bungarotoxin binding site (BBS) in the extracellular loop and YFP in the C-terminus (Orai-BBS-YFP) (Figure 2-figure supplement 1A) [9]. Resting and store-depleted Orai-BBS-YFP expressing HEK293 cells were labelled with Alexa-647 conjugated alpha-bungarotoxin (BTX-A647), washed, fixed and analyzed using flow-cytometry (Figure 2A-B) as well as microscopy (Figure 2C) to estimate total expression levels and PM localization of Orai1. We found no difference in the total (Figure 2A) or surface Orai1 (Figure 2B-C) expression in any group. Verification of knockdown efficiency in these HEK293 cells is shown in (Figure 2-figure supplement 1(B-C)). Similar observations were made in Jurkat T cells transiently transduced with pMSCV-Orai1-BBS-YFP (Figure 2D) (Figure 2-figure supplement 1D) as well as U2OS cells (Figure 2-figure supplement 2A-E) stably expressing the same plasmid construct as described above for HEK293. Neither total Orai1 expression levels, nor it’s PM localization was found to be altered upon store-depletion or STX11 ablation in these cell types.

**Figure 2.**
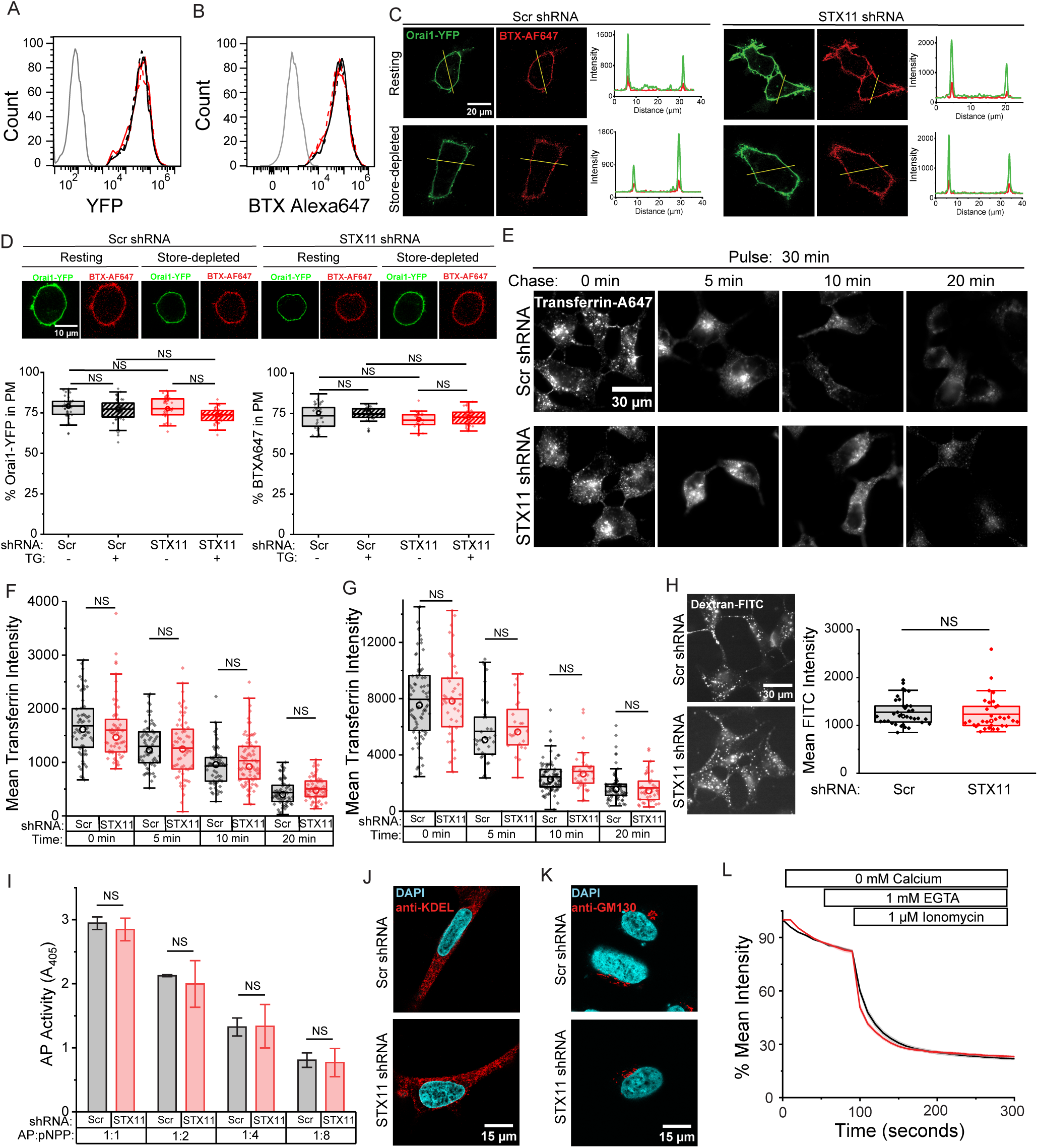
STX11 depletion does not affect PM Orai1 levels and global membrane trafficking or secretion. (**A-C**) Measurement of Bungarotoxin binding site (BBS) tagged Orai1-YFP in the PM of scramble and STX11 shRNA treated, resting and store-depleted, HEK293 cells. Measurement of Orai1-YFP **(A)** and Alexa Fluor 647-conjugated Bungarotoxin binding **(B)** in population of cells using flow-cytometry. Quantification of Orai1-YFP and BTX-A647 in individual cells using confocal imaging **(C)**. **(D)** Confocal images and quantification of Orai1-YFP and BTX-A647 in scramble and STX11 shRNA treated, resting and store-depleted, Jurkat T cells. (n=30-40, N=3) **(E-G)** Scr or STX11 shRNA treated HEK293 and Jurkat T cells were loaded with Alexa-647-Transferrin (30 minute pulse), washed, chased for 0, 5, 10 and 20 min and imaged to quantify total Transferrin levels inside each cell. Representative images from HEK293 cells **(E)** and quantification from 50-60 HEK293 **(F)** and Jurkat T cells **(G)** is shown. N=3 (**H**) Quantification of non-receptor mediated endocytosis in STX11-depleted HEK293 cells. Representative images of FITC-conjugated Dextran uptake in Scr and STX11-shRNA treated HEK293 cells (left panel) and quantification of the mean intensity per cell (right panel). (**I**) Scr or STX11 shRNA treated HEK293 cells were transfected with secretory Alkaline phosphatase (AP) expressing plasmid and levels of AP secreted into the culture supernatants were measured using p-Nitrophenyl phosphate substrate at 48h post-transfection. **(J-K)** Immunolabelling of HEK293 cells to assess ER **(J)** and Golgi **(K)** health. Scr and STX11 shRNA treated HEK293 cells were stained with anti-KDEL **(J)** and anti-GM130 antibodies **(K)** respectively and imaged. **(L)** Measurement of ER calcium content in Scr and STX11 shRNA treated HEK293 cells using transiently transfected ER-localized CEPIA. Cells were incubated in Ringer’s buffer containing 0 mM calcium followed by 1 mM EGTA and stimulated with 1 μM ionomycin. (N=3).

To determine whether STX11 deficient cells harbored a general defect in vesicle trafficking and fusion, we used the Transferrin receptor (TfR) endocytosis and recycling assay, which internalizes iron bound holo-transferrin via endocytosis of cell surface TfR, followed by exocytosis and extracellular release of iron free apo-Transferrin [22]. RNAi treated HEK293 (Figure 2E-F) and Jurkat T cells (Figure 2G) were incubated with Alexa-647 conjugated Transferrin (A647-TfR) for a 30 minute endocytic pulse. Excess A647-Transferrin was washed off and the emptying of internalized A647-Transferrin via recycling of TfRs over time was quantified at various time points (chase). Levels of A647-Transferrin after endocytosis (30-minute pulse-0-minute chase) as well as it’s exocytosis during 5, 10, 20-minute chase times were found to be identical between STX11 or Scr shRNA treated cells. Non-receptor mediated fluid-phase endocytosis of FITC-conjugated Dextran was also found to be identical between Scr and STX11 shRNA treated HEK293 cells (Figure 2H). In addition, we transfected HEK293 cells with a plasmid containing alkaline phosphatase (AP) enzyme fused to Ig-κ chain secretion signal peptide at N-terminal, which allowed us to measure AP secreted into the culture supernatant via the biosynthetic-exocytosis pathway. Scramble and STX11 depleted cells showed comparable levels of AP secretion (Figure 2I). Overall, these experiments did not indicate any generalized defects in the membrane recycling or secretory pathways upon depletion of STX11 in multiple cell types.

Using a dominant negative mutant of N-ethylmaleimide sensitive factor (NSF), which blocked TfR recycling but not SOCE, we have previously shown that CRAC channels are not dependent on vesicle trafficking for their activation [9]. Yet, Q-SNAREs are often found in complex with other SNAREs such as SNAP23/25 in target membranes and some previous studies have implicated the complex of SNAP23/25 and STX1A in the regulation of specific ion channels [23, 24]. Each of the three SNAPs, SNAP23/SNAP25/SNAP29 are capable of contributing two SNARE domains to the *trans-* SNARE complex. RNAi mediated depletion of SNAP23/SNAP25/SNAP29 did not show a significant reduction in SOCE of HEK 293 cells (Figure 2, figure supplement 3A-C). To determine whether SNAP23/SNAP25/SNAP29 might still form a part of the STX11:Orai complex to regulate SOCE, we co-expressed myc-tagged Orai1 with untagged SNAPs and performed respective co-IPs. We did not find any interaction between either of the three SNAPs and Orai1 (Figure 2-figure supplement 3D). Similar studies were performed by co-expressing YFP-Stim1 and the three myc-tagged SNAPs. Again, no interaction was found between any of the SNAPs and Stim1 (Figure 2-figure supplement 3E). Collectively, these data established that Orai1 is constitutively expressed in the PM, and not vesicles, of diverse cell types and STX11 depletion does not lead to a general defect in vesicle fusion or PM targeting of Orai1.

Labelling of STX11 depleted cells with ER, Golgi organelle specific markers also showed no abnormalities (Figure 2J-K) suggesting that the overall health and function of ER and Golgi were not adversely affected by STX11 depletion. To assess ER calcium content, we ectopically expressed ER-targeted CEPIA in control and STX11 depleted cells and measured the mean intensity. Both groups showed comparable ER calcium content (Figure 2L).

Previous studies have shown that SOCE in T cells is mediated by CRAC channels formed by Orai proteins. To test whether CRAC current (I_CRAC_) was affected by knockdown of STX11 in T cells, we performed whole cell patch clamp recordings on scramble and STX11 shRNA treated Jurkat T cells (Figure 3A-B). Leak-subtracted I_CRAC_ was recorded in 20 mM Ca^2+^ and a Na^+^-based divalent cation free (DVF) solution. These recordings showed that both the Ca^2+^ and DVF CRAC currents were significantly decreased in the STX11 shRNA treated cells with the normalized currents shown in Figure 3C. The electrophysiological properties of the residual current in STX11 shRNA treated cells was indistinguishable from I_CRAC_ in scramble shRNA treated cells in terms of blockade of the Ca^2+^ current by La^3+^ and depotentiation of the DVF current over tens of seconds. Moreover, the reversal potential of the Ca^2+^ and DVF currents were similar between scramble and STX11 shRNA treated Jurkat T cells, suggesting that STX11 depletion does not affect the calcium selectivity of CRAC channels (Figure 3D). Together, these results demonstrate that STX11 works by specifically modulating I_CRAC_ in T cells.

**Figure 3.**
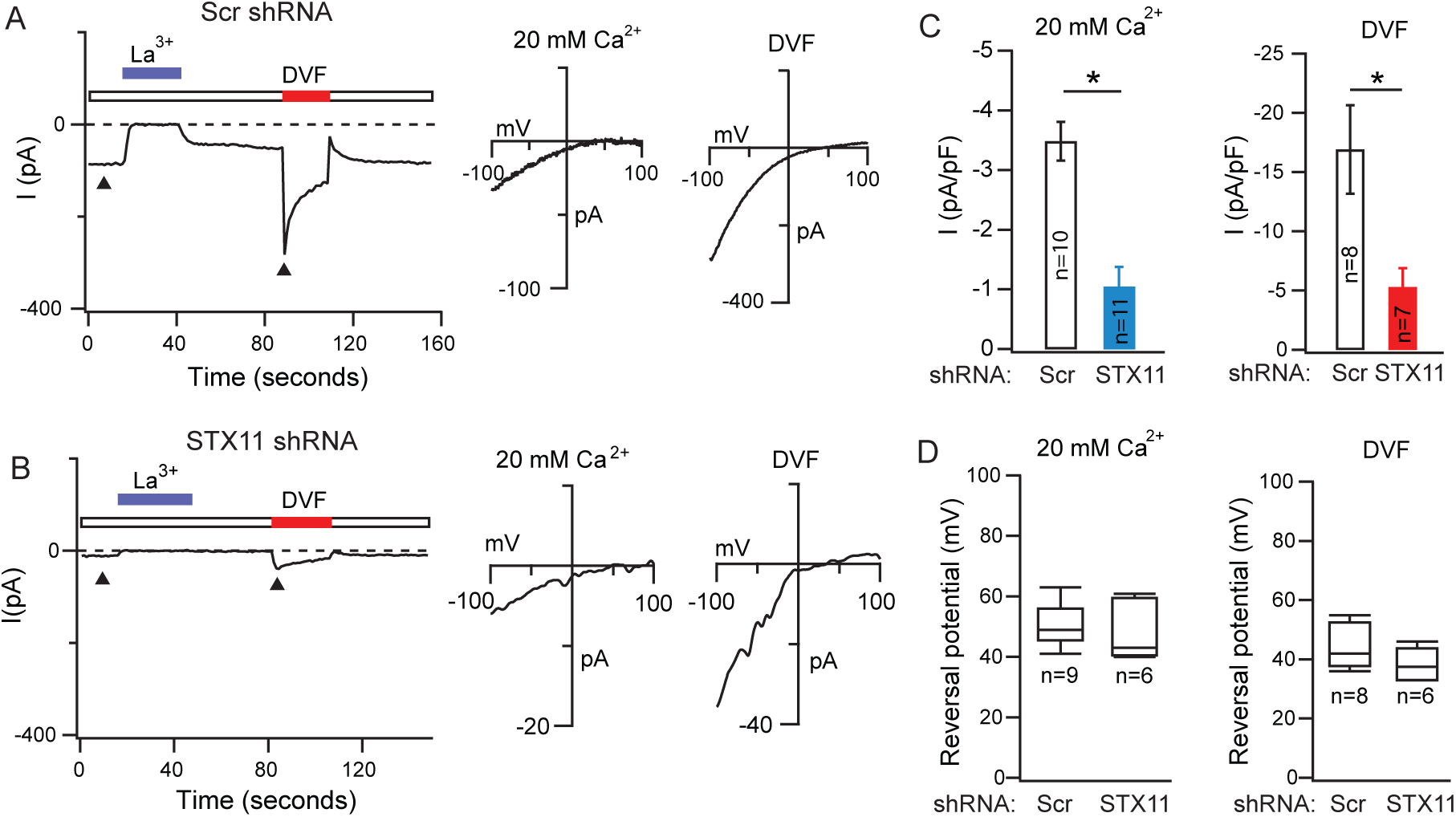
STX11 regulates I_CRAC_. (**A-D**) Measurement of I_CRAC_ in STX11 depleted Jurkat T-cells in the whole-cell recording configuration in 20 mM extracellular Ca^2+^ Ringer’s solution. I_CRAC_ was induced by passive depletion of intracellular Ca^2+^ stores by dialyzing 8 mM BAPTA into the cell via the patch-pipette. (**A**) Representative current at −100 mV in Jurkat T cells treated with Scr shRNA construct. The current is blocked by extracellular La^3+^ (10 µM) and replacing the 20 mM Ca^2+^ Ringer’s solution with a divalent free solution (DVF) evokes a large Na^+^ current which depotentiates over tens of seconds. The current-voltage (I-V) relationship of the Ca^2+^ and DVF currents are shown on the right. (**B**) I_CRAC_ from a Jurkat T cell treated with STX11 shRNA. Both Ca^2+^ and Na^+^ current amplitudes are reduced relative to control cells. The I-V relationships (right plots) show no change in ion selectivity. (**C-D**) Summary of the current amplitudes of Ca^2+^ and Na^+^ currents and current reversal potentials in Scr and STX11 knockdown cells.

CRAC channel mediated sustained calcium influx is crucial for the activation of a key calcium dependent transcription factor known as nuclear factor of activated T cells (NFAT) [4, 25]. We stimulated control and STX11 depleted Jurkat T cells with Thapsigargin (TG)+PMA, prepared nuclear, cytosolic extracts and subjected them to western blot using anti-NFAT1 antibody (Figure 4A). Nuclear translocation of NFAT was severely compromised in STX11 depleted T cells. In addition, immunofluorescence imaging and quantification of total *versus* nuclear NFAT from anti-CD3 stimulated Jurkat cells showed a defect in nuclear translocation of NFAT1 in STX11 depleted cells (Figure 4B-C). To determine whether NFAT induced gene expression was affected [4, 25], we quantified IL-2 mRNA by qPCR and secreted IL-2 levels from culture supernatants of anti-CD3 stimulated Jurkat T cells. STX11 depleted cells showed a significant decrease in IL-2 gene expression as well as secretion (Figure 4D, 4E). We stimulated HD and FHLH4 patient T cells either through T cell receptor using anti-CD3 or with phorbol myristate acetate (PMA) + Ionomycin and assessed IL-2 gene expression. Importantly, induction of IL-2 expression was significantly defective in FHLH4 patient T cells but could be largely restored with Ionomycin (Figure 4F). We next performed granule release assay on the *in vitro* cultured HD and FHLH4 patient CD8 T cells in response to anti-CD3 mediated stimulation and observed a significant defect in the FHLH4 patient CD8 T cell degranulation (Figure 4G, 4H). To determine whether the reduced granule release results from a direct defect in vesicle fusion or SOCE, we stimulated cells with Ionomycin along with (PMA). Remarkably, CD8 T cell degranulation was fully restored in FHLH4 T cells suggesting that defective SOCE primarily causes cytolytic defects in FHLH4 T cells. Of note, the patient exhibited all symptoms of FHLH4 disease and we observed strong suppression of SOCE (Figure 1I), IL-2 gene expression and secretion (Figure 4D-F). However, *in vitro* culture of FHLH4 patient CD8 T cells with IL-2 has been previously shown to partially or fully restore cytolytic defects by other groups [18, 26]. Therefore, considering previous reports, a relatively lower (50%) defect in CD8 T cells degranulation was not surprising [18, 26]. Taken together, the data show that defects in FHLH4 T cell degranulation as well as interleukin-2 gene expression originate from suppressed SOCE and not vesicle trafficking.

**Figure 4.**
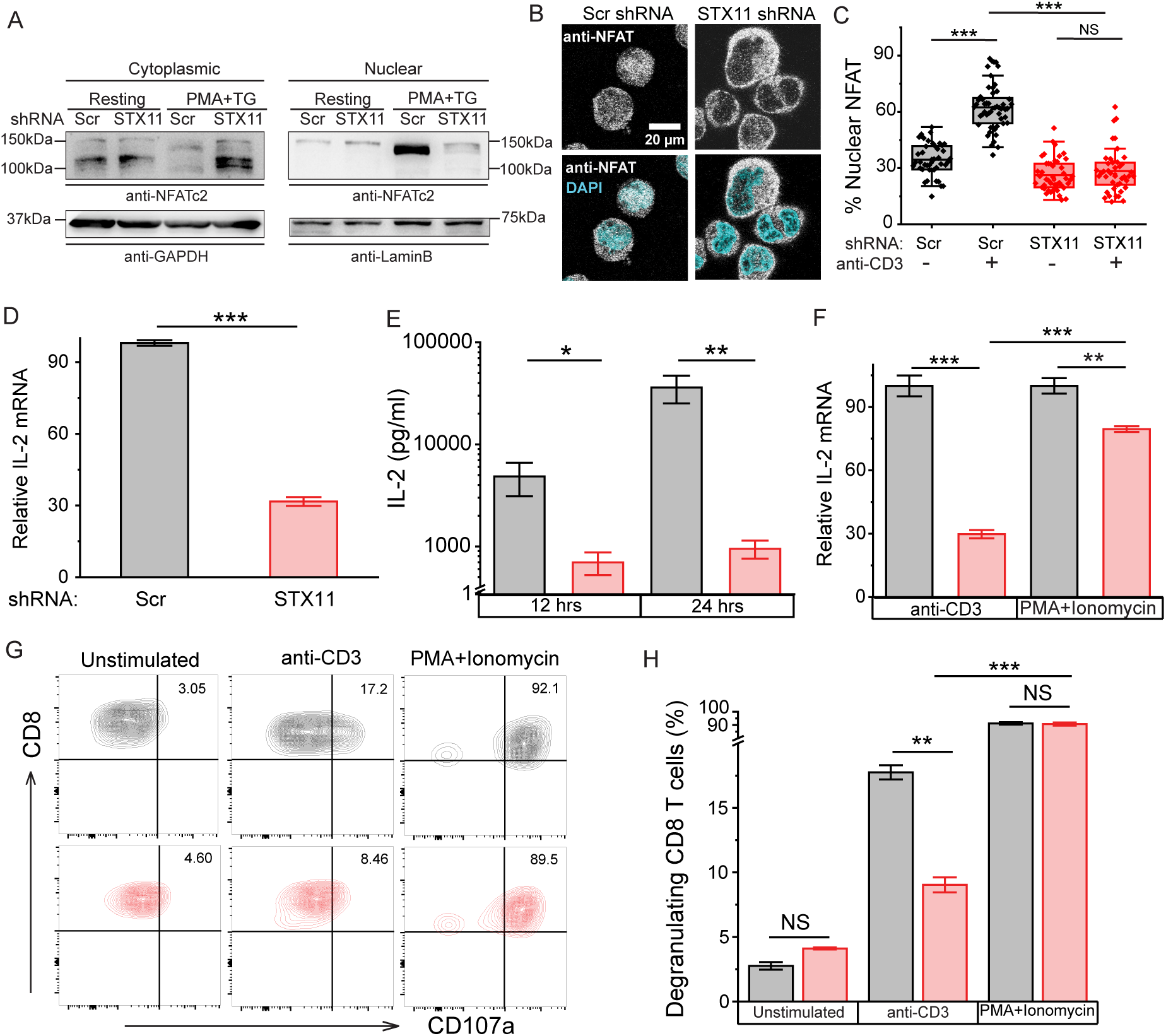
STX11 regulates NFAT activation, IL-2 gene expression and degranulation in FHLH4 patient T cells. (**A-C**) Estimation of nuclear translocation of NFAT. (**A**) Western blot showing nuclear translocation of NFAT in Jurkat T cells treated with scr or STX11 shRNA and stimulated with PMA+TG. N=3. (**B**) Representative confocal images of control and STX11 depleted Jurkat T cells stimulated with anti-CD3, immunolabelled with anti-NFAT1 antibody and counter-stained with DAPI. (N=3) (**C**) Box and whisker plot showing quantification of nuclear NFAT from cells populating 10 randomly chosen fields per group in (**B**). (**D**) Quantification of IL-2 mRNA in anti-CD3 stimulated Jurkat T cells treated with scr or STX11 shRNA using qPCR. The bars show mean ± SE of relative IL-2 mRNA. N=3. (E) Quantification of IL-2 EIA on supernatants of scr (black) or STX11 (red) shRNA treated Jurkat T cells stimulated with anti-CD3. (**F**) qPCR analysis of IL-2 mRNA expression in various stimulated HD (black) and FHLH4 T cells (red). (**G-H**) Granule release assay performed on HD (black) and FHLH4 CD8 T (red) cells (**G**) and its quantification (**H**).

Unlike most Q-SNAREs, STX11 lacks a transmembrane domain but harbors a stretch of cysteines close to the C-terminus and several basic residues, spread throughout the sequence (Figure 5-figure supplement 1A) [27–29]. The cellular localization of STX11 is not established because, unlike primary T cells, native STX11 is undetectable in Jurkat T and HEK293 cell lines using available commercial antibodies. Tagging STX11 on either N- or C-terminus result in partial degradation (Figure 5-figure supplement 1B) and complete mis-localization of the protein (Figure 5-figure supplement 1C). Therefore, we inserted an HA tag inside the N-terminal unstructured region of human STX11. The internal HA tagged STX11 as well as un-tagged STX11 localized in plasma membrane of most HEK293 cells (Figure 5A) and co-localized with Orai1-YFP in resting (Figure 5B) as well as store-depleted cells (Figure 5C). Figure 5D shows quantification of Orai1 and STX11 co-localization using Pearson’s correlation coefficient. Notably, despite loss of all terminal cysteines and a short terminal part of the SNARE domain, the FHLH4 mutant STX11 did not lose the ability to localize to PM (Figure 5E), therefore, terminal cysteines do not appear to target STX11 to the PM.

**Figure 5:**
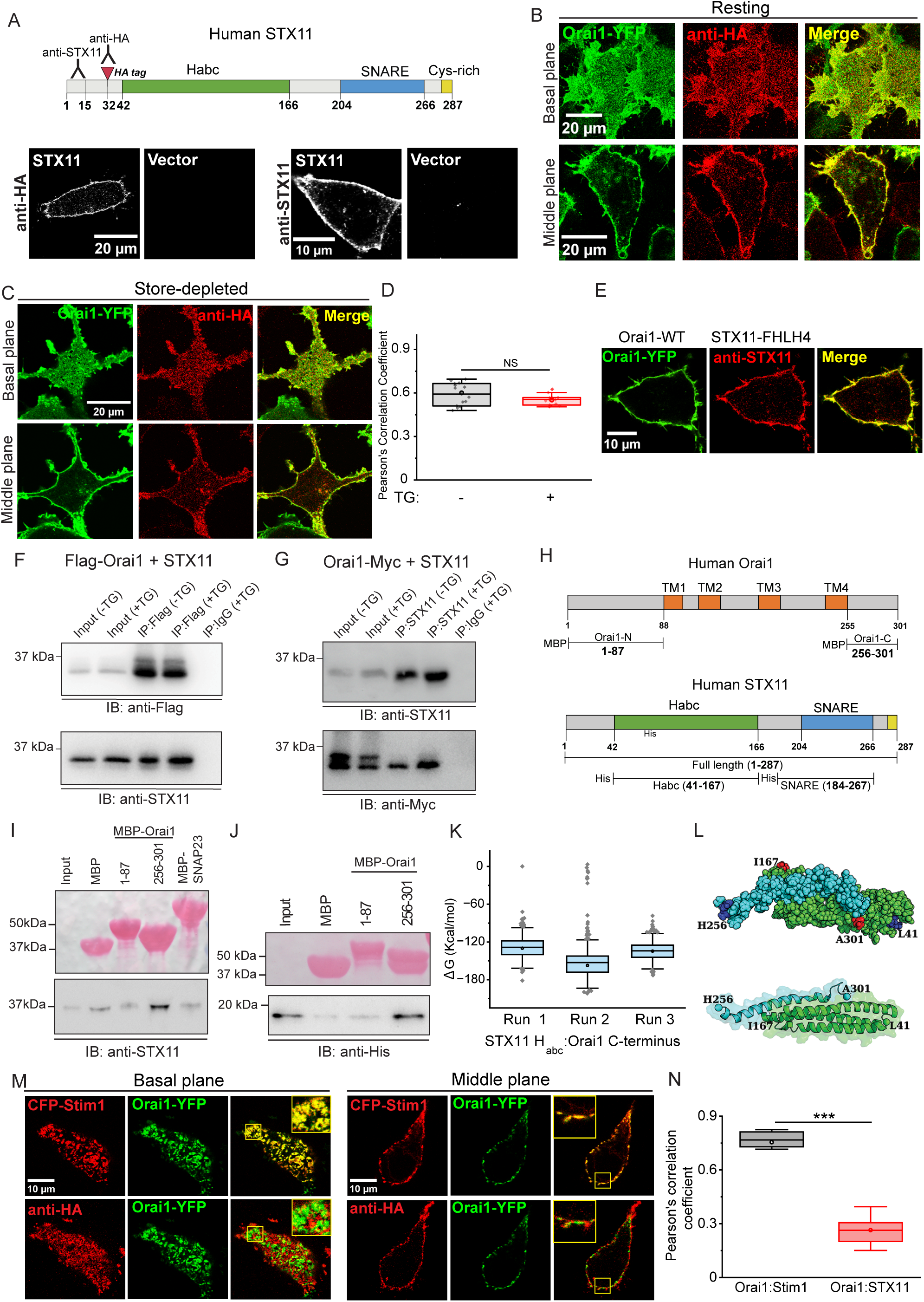
STX11 directly binds resting Orai1 in plasma membrane. Intracellular localization of STX11 and its association with Orai1. (**A**) Representative confocal images of HEK293 cells transfected with either HA-tagged STX11 or untagged STX11 and stained using anti-HA antibody or anti-STX11 antibody respectively. N=3 (**B-D**) Localization of HA-tagged STX11 with respect to Orai1-YFP in the basal as well as nuclear plane of resting **(B)** and store-depleted **(C)** HEK293 cells. (D) Pearson’s correlation coefficient showing Orai1 and STX11 colocalization from **(B)** and **(C)**. N=3. (**E**) Representative confocal images of FHLH4 mutant STX11 expressing HEK293 cells. (**F-G**) Co-IP to assess STX11 binding to Orai1. Whole cell lysates of resting and store-depleted HEK293 cells co-expressing either Flag-Orai1 and STX11 (**F**) or Orai1-Myc and STX11 (**G**) were subjected to IP and Western blot using anti-Myc, anti-Flag or anti-STX11 antibodies, as indicated. (N=3) **(H)** Schematic showing key domains of STX11 and Orai1 used for *in vitro* pull-down assays. (**I**) Pull-down assay showing *in vitro* binding of His-tagged STX11 to Orai1. (Top) Ponceau S staining showing the input of MBP alone or MBP-tagged Orai1 fragments. (Bottom) Western blot using anti-STX11 antibody. Input:1/5^th^ of the protein. N=4. (**J**) Pull-down assay showing *in vitro* binding of His-tagged H_abc_ domain of STX11 to Orai1 fragments. (Top) Ponceau S staining showing the input of MBP alone or MBP-tagged Orai1 fragments. Input:1/5^th^ of the protein. (Bottom) Western blot using anti-His antibody. N=3. (**K**) Binding free energy distribution of STX11-H_abc_ and Orai1 C-terminus interactions. (**L**) Sphere and cartoon representation of the structure of STX11 H_abc_ (green) and Orai1 C-terminus (cyan) complex after MD simulation. The N-termini are highlighted in blue, and C-termini in red. (**M**) Representative confocal images showing localization of STX11 with respect to Orai1:Stim1 puncta in store-depleted cells. HEK293 cells expressing Orai1-YFP, CFP-Stim1 were transfected with HA-tagged STX11, store-depleted and stained using anti-HA antibody. (N=3) (**N**) Quantification of the localization of STX11 with respect to Stim1:Orai1 puncta in store-depleted cells.

To test whether STX11 and Orai1 formed a complex, we co-expressed STX11 with either Flag- or Myc-tagged Orai1 in HEK293, prepared whole cell lysates (WCLs) and subjected them to co-immunoprecipitation (co-IP) followed by western blot using either anti-Flag, anti-Myc or anti-STX11 antibodies. Both Orai1 and STX11 could co-IP each other (Figure 5F-G). To further assess whether STX11 directly bound Orai1, we expressed and purified from *E. coli*, MBP-tagged N- and C-terminal cytosolic tails of Orai1 (Figure 5H). *In vitro* pull-down assay performed by incubating MBP-tagged Orai1 cytoplasmic tails with full length soluble His-tagged STX11 showed that the C-terminus of Orai1 directly bound STX11 (Figure 5I). SNAREs typically utilize their SNARE domain for associating with other SNAREs [30]. To identify the domain of STX11 involved in the regulation of SOCE, we expressed His-tagged H_abc_ and SNARE domains of STX11 (Figure 5H) in *E. coli*, purified and assessed their binding to the MBP-tagged Orai1 tails using pull-down assay. We found that the H_abc_ domain of STX11 bound to the C-terminus of Orai1 (Figure 5J) but the SNARE domain of STX11 did not show any binding to Orai1 and only a faint binding to SNAP23 (Figure 5-figure supplement 1D). We next used AlphaFold3 (AF3) to obtain deeper insights into the domain interactions of STX11 and Orai1. Of all the possible combinations, the complex of STX11 H_abc_ and Orai1 C-terminus resulted in a significantly high prediction score (Table 1). We, therefore, executed AF3 predictions with different initial seeds to generate multiple models of STX11 H_abc_ with Orai1 C-terminus. We calculated the contact frequency of interface residues in different models and found that multiple residues showed frequency greater than 80% (Figure 5-figure supplement 2A). For further analysis, we considered the predicted model with the best score (Table 2). We next performed all-atom MD simulation in aqueous environment (Movies 1-3) to assess the conformation, interaction stability of the STX11 H_abc_ and Orai1 C-terminus complex, which was stable throughout the simulation time as suggested by RMSD (Figure 5-figure supplement 2B), and binding free energy (ΔG) values (Figure 5-figure supplement 3) (Figure 5K). The individual trajectories were concatenated and clustered to obtain a centroid structure for this complex. There were 46 clusters, and the largest cluster, with 52 members, showed an elaborate protein-protein interface where STX11 H_abc_ was found to be oriented in an anti-parallel orientation to the Orai1 C-terminus (Figure 5L).

**Table 1:** AF3 predictions for STX11:Orai1 complex formation.

| Proteins/Domains | ipTM | pTM |
| --- | --- | --- |
| Full-length Orai1 - Full-length STX11 | 0.11 | 0.35 |
| STX11-Habc_41-167 and Orai1_N-term_1-87 | 0.27 | 0.59 |
| **STX11-Habc_41-167 and Orai1_C-term_256-301** | **0.53** | **0.79** |
| STX11-SNARE_183-267 and Orai1_N-term_1-87 | 0.08 | 0.34 |
| STX11-SNARE_183-267 and Orai1_C-term_256-301 | 0.18 | 0.45 |
| STX11-Habc_41-167 and full-length Orai1 | 0.24 | 0.38 |
| STX11-SNARE_183-267 and full-length Orai1 | 0.08 | 0.37 |
| Orai1_N-term_1-87 and full-length STX11 | 0.15 | 0.5 |
| Orai1_C-term_256-301 and full-length STX11 | 0.16 | 0.54 |

**Table 2:**
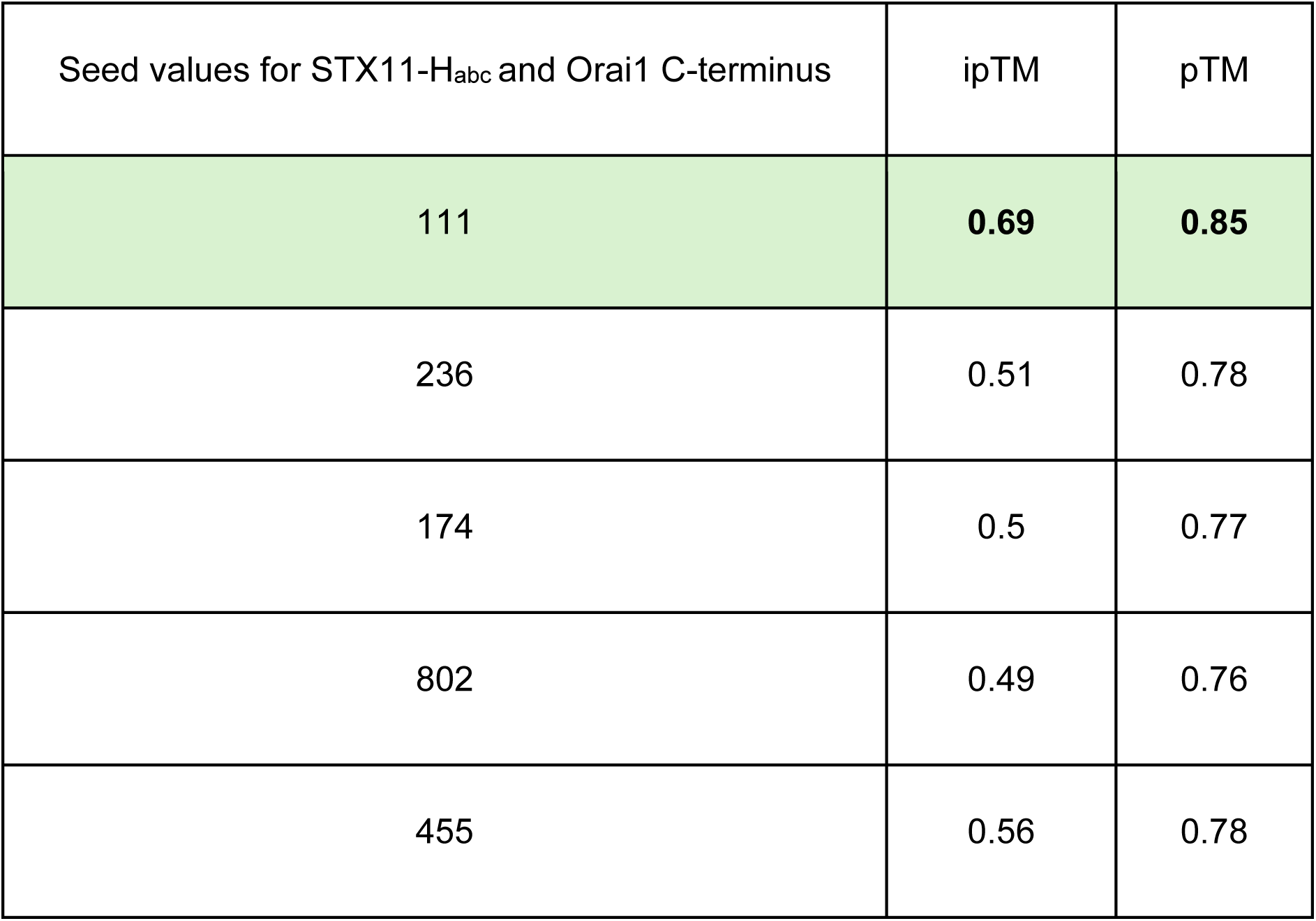
AF3 predictions for STX11-H_abc_ and Orai1 C-terminus with different seeds.

Because STX11 binds the C-terminus of Orai1 and the region largely overlaps with the Stim1 binding domain, we next sought to determine whether STX11 clusters with Orai:Stim puncta. We over-expressed HA-tagged STX11 in a Orai1-YFP and CFP-Stim1 expressing stable HEK293 cell line, store-depleted with TG, and stained using anti-HA antibody. We did not observe co-clustering of STX11 in Orai:Stim puncta even 8 minutes post store-depletion (Figure 5M-N). Accordingly, STX11 did not co-IP with Stim1 in store-depleted cells (Figure 5-figure supplement 4A). These data suggest that STX11 comes off Orai1 C-terminus prior to Stim1 dependent clustering of Orai1. Since co-IP studies suggested sustained interaction between STX11 with Orai1 (Figure 5F-G), we quantified the fraction of Orai1 that resides inside Stim:Orai puncta w.r.t total Orai1 in PM and found that post TG addition, nearly 60% of Orai1 remains outside of Stim1 puncta even in unmanipulated cells (Figure 5-figure supplement 4B). We next used AF3 to generate the complex of Stim-Orai activating region (SOAR) domain of Stim1 and Orai1 (Figure 5-figure supplement 5A). Given that SOAR is a dimer under resting state, we used two copies each of SOAR and Orai1 [31] [32]. The prediction scores for their complex are shown in Table 3. The best scoring model among multiple runs of AF3 was considered for MD simulations on the complex of Orai1 C-terminus (256-301) and SOAR. The ΔG values for the complex are shown in (Figure 5-figure supplement 5B-C). Three independent MD simulations were run for the complex (Movies 4-6) which showed an initial conformational change which stabilized thereafter (Figure 5-figure supplement 6A). All three simulation trajectories were concatenated and clustered to obtain a centroid structure for this complex (Figure 5-figure supplement 6B). There were 43 clusters, and the largest cluster had 77 structures. Collectively, these data suggest that STX11 directly binds Orai1 with an affinity that is physiological but relatively poorer when compared to the binding of SOAR dimers to Orai1. Therefore, presence of high density of SOAR dimers in ER-PM junctions of store-depleted cells could potentially outcompete STX11:Orai1 interactions by progressively allowing formation of SOAR:Orai1 oligomers. It is also possible that another, yet to be characterized, molecular step and/or protein actively catalyzes the removal of STX11 from Orai1 prior to its entrapment in ER-PM puncta. Future studies will be able to systematically assess these possibilities.

**Table 3:** AF3 predictions for SOAR and Orai1 in 2:2 ratio.

| 2 copies of SOAR and 2 copies of<br>Orai1_C-term_256-301 | ipTM | pTM |
| --- | --- | --- |
| Seed_101 | 0.73 | 0.78 |
| Seed_102 | 0.73 | 0.78 |
| Seed_103 | 0.73 | 0.78 |
| Seed_104 | 0.73 | 0.78 |
| **Seed_105** | **0.74** | **0.79** |

The major interactions predicted between STX11 and Orai1 from all-atom MD simulations are highlighted in (Figure 5-figure supplement 3B-C). To assess the significance of these interactions, we performed site-directed mutagenesis of two key residues of STX11 predicted to be involved in binding to Orai1 (Figure 6A). Ectopic expression of untagged N147A_E150A STX11 showed PM localization (Figure 6A) as well as stable expression (Figure 6B) but failed to rescue SOCE in STX11 depleted Jurkat T cells (Figure 6C-D). We next expressed His-tagged wildtype and N147A_E150A mutant STX11 in *E. coli.*, purified (Figure 6E) and assessed their ability to bind MBP-tagged Orai1 C-terminal tail in an *in vitro* pull-down assay (Figure 6F-G). Compared to wildtype STX11, N147A_E150A mutant STX11 showed reduced binding to Orai1 C-terminus. Further, we introduced four mutations, R289A_E272A_E275A_E278A, in the Orai1 C-terminus (Figure 6H), tagged with YFP and expressed in HEK293 cells. The Orai1 mutant localized to PM (Figure 6H) but failed to rescue SOCE in Orai1 deficient cells (Figure 6I-J). We expressed and purified MBP-tagged R289A_E272A_E275A_E278A mutant Orai1 C-terminus from *E. coli* and assessed its ability to bind purified wildtype as well as N147A_E150A mutant STX11 in a pull-down assay (Figure 6K-L). Binding of Orai1 mutant to both was significantly compromised. Because Stim1 binds an overlapping region of the Orai1 C-terminus, we next tested whether mutation of Orai1 residues affected Stim1 binding. We co-expressed YFP-tagged soluble CRAC activation domain (CAD) of Stim1 (YFP-CAD) (Figure 6M) with either CFP-tagged wildtype or mutant Orai1 in HEK293 cell line. YFP-CAD localizes to the cytosol in HEK293 cells but in cells overexpressing Orai1, a significant fraction of it localizes to plasma membrane due to its association with Orai1 [10]. We quantified relative YFP-CAD levels in the plasma membrane of both groups but found no difference in the PM recruitment of CAD (Figure 6N) suggesting that the inability of R289A_E272A_E275A_E278A mutant Orai1 to support SOCE does not result from defective binding to CAD/SOAR. However, CAD was unable to stimulate constitutive calcium influx from R289A_E272A_E275A_E278A mutant Orai1 in the above experiment (Figure 6O-P). Collectively, these data demonstrate a direct role of STX11 in binding and regulation of Orai1.

**Figure 6.**
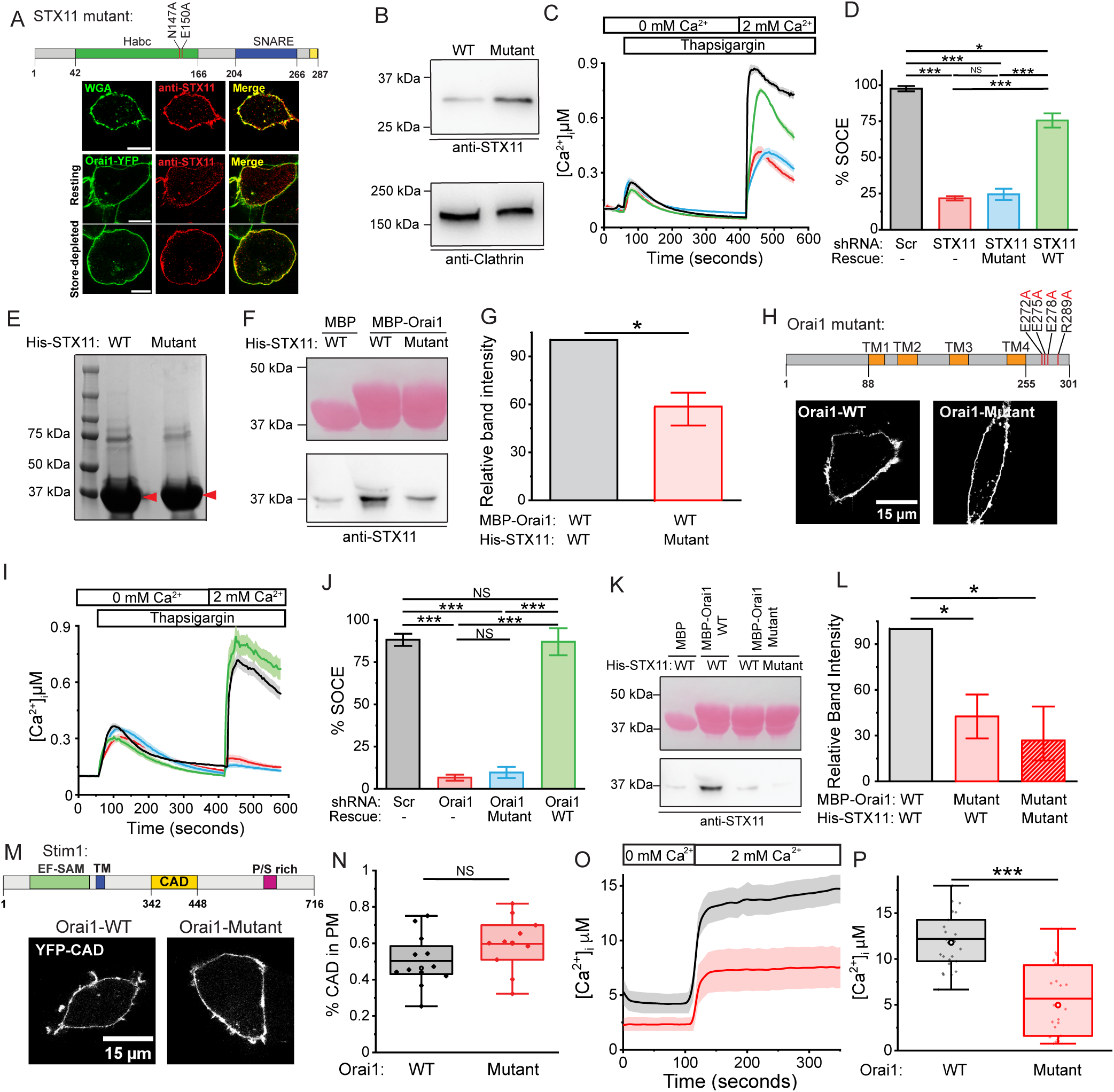
Residues involved in Orai1:STX11 complex formation are crucial for SOCE. Analysis of STX11 and Orai1 mutants. (**A**) Schematic showing STX11 mutations (Top). STX11 N147A_E150A mutant expressing HEK293 co-labelled with Alexa Fluor-488 conjugated wheat germ agglutinin (WGA) or co-transfected with Orai1-YFP and immunolabelled with anti-STX11 to assess co-expression in PM of resting and store-depleted cells (Bottom). Scale bar 10μm. (**B**) Western blot to compare total expression of mutant and wildtype STX11 using anti-STX11 antibody. (**C-D**) Representative Fura-2 calcium imaging assay (**C**) and quantification of repeats (**D**) measuring rescue of SOCE in Jurkat cells treated either with scr shRNA (black) or STX11 shRNA expressing empty vector (red), wildtype STX11 (green) or N147A_E150A mutant STX11 (blue). (**E**) Coomassie blue stained SDS-PAGE showing expression and purification of His-tagged wildtype or N147A_E150A mutant STX11 from *E. coli.* (**F-G**) Pull-down assay measuring their binding to MBP-tagged Orai1 C-terminus. (Top) Ponceau S staining showing the input of MBP-tagged Orai1 fragments. (Bottom) Western blot using anti-STX11 antibody. N=4 (**G**) Quantification of STX11 band intensities in (F). (**H**) Schematic showing Orai1 mutations (Top). Confocal images of HEK293 cells expressing YFP-tagged wildtype or R289A_E272A_E275A_E278A mutant Orai1 (Bottom). **(I-J)** Representative Fura-2 calcium imaging assay (**I**) and quantification of repeats (**J**) measuring rescue of SOCE in HEK293 cells treated either with Scr (black) or Orai1 shRNA expressing empty vector (red), wildtype Orai1 (green) or R289A_E272A_E275A_E278A mutant Orai1 (blue). (**K-L**) Pull-down assay measuring the binding of MBP-tagged wildtype or R289A_E272A_E275A_E278A mutant Orai1 C-terminus to wildtype or N147A_E150A mutant STX11. (Top) Ponceau S staining showing the input of MBP-tagged Orai1 fragments. (Bottom) Western blot using anti-STX11 antibody. N=4. (**L**) Quantification of STX11 band intensities in (K). (**M-N**) Representative confocal images (**M**) and quantification (**N**) of PM localized YFP-CAD in CFP-tagged wildtype or R289A_E272A_E275A_E278A mutant Orai1 expressing HEK293 cells. (**O-P**) Representative Fura-2 calcium imaging assay (**O**) and quantification of repeats (**P**) of constitutive calcium flux from cells imaged in panels **M-N**.

Upon store depletion, Stim1 localizes to ER-PM junctional regions where it traps and co-clusters with Orai1. To further understand the mechanism, using confocal microscope we first imaged HEK293 cells expressing CFP-Orai1, Stim1-YFP under resting conditions and quantified the levels of Orai1 and Stim1 in ER-PM junctions (puncta) of store-depleted control and STX11 depleted cells. The distribution of Orai1 and Stim1 was normal in resting cells (Figure 7A). However, in store-depleted cells, clustering of CFP-Orai1 was visibly defective in STX11 RNAi treated cells (Figure 7B), even though the intensities (Figure 7C) and area of Stim1-YFP puncta were normal (Figure 7D).

**Figure 7:**
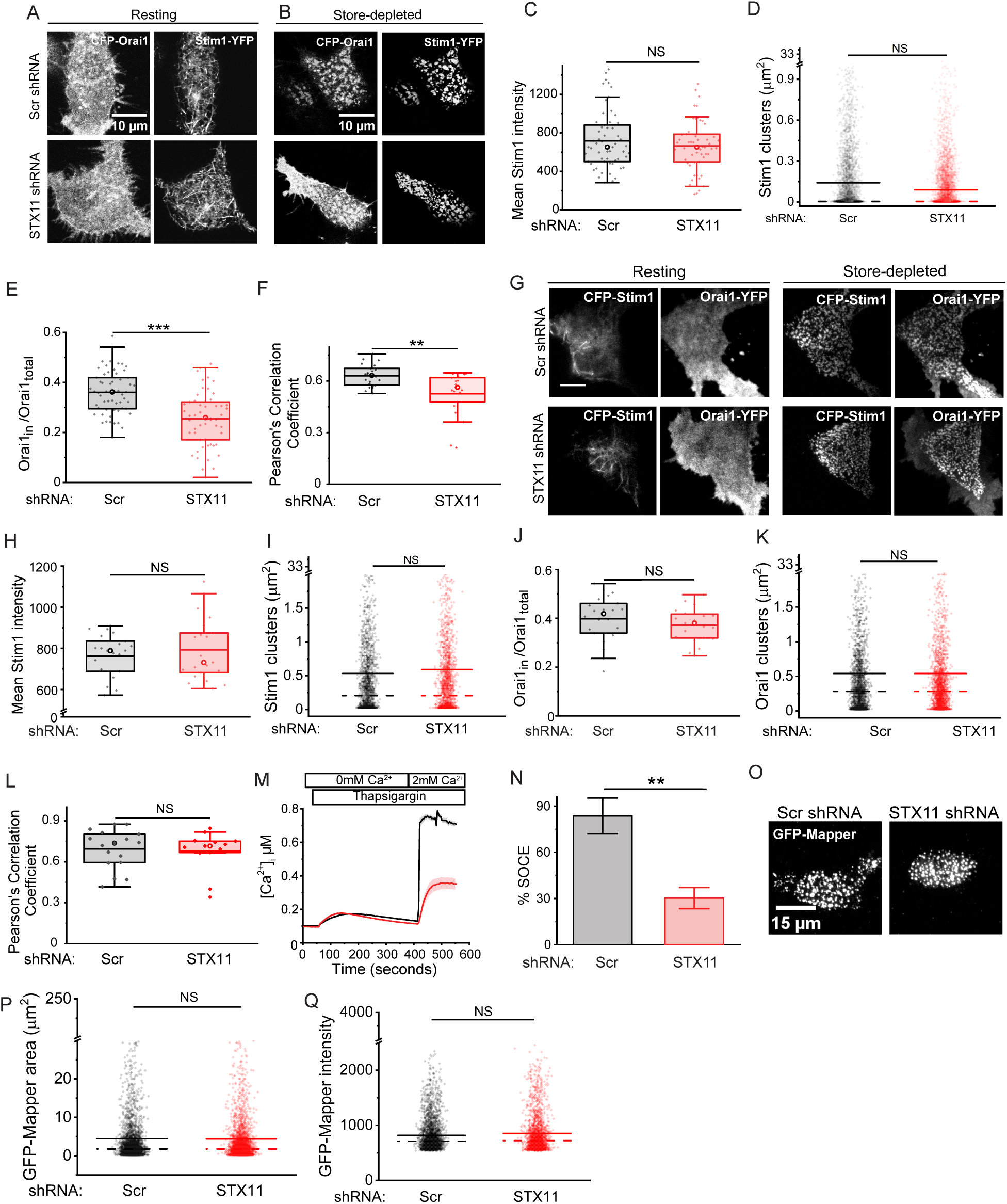
Non-functional clustering of Orai1 in STX11 depleted cells. **(A-L)** Quantification of Orai1 and Stim1 intensities in the ER-PM puncta of store-depleted cells. (**A-B**) Representative images of resting (**A**) and store-depleted (**B**) shRNA treated HEK293 cells expressing N-terminal CFP tagged Orai1 (CFP-Orai1) and C-terminal YFP tagged Stim1 (Stim1-YFP) from confocal microscopy. (**C-F**) Mean Stim1-YFP intensity (**C**) and area (**D**) of Stim1 clusters. (**E**) Fraction of CFP-Orai1 intensities inside Stim1-YFP clusters of shRNA treated cells, post store-depletion. (**F**) Pearson’s correlation coefficient of Stim1 and Orai1 co-localization inside Stim1 puncta. (**G-J**) Quantification of Orai1 and Stim1 intensities in the ER-PM puncta of C-terminal YFP-tagged Orai1 (Orai1-YFP) and N-terminal CFP-tagged Stim1 (CFP-Stim1) expressed in HEK293 cells and imaged using TIRF microscopy. **(G)** Representative TIRF images of resting and store-depleted shRNA treated HEK293 cells. (**H-K**) Mean CFP-Stim1 intensity (**H**) and area (**I**) of Stim1 clusters. Fraction of Orai1-YFP intensities (**J**) and area (**K**) of Orai1 clusters in control and STX11 depleted cells. **(L)** Pearson’s correlation coefficient of Stim1 and Orai1 co-localization inside Stim1 puncta (**M-N**) Representative Fura-2 calcium imaging assay (**M**) and quantification of repeats (**N**) measuring TG induced SOCE in Orai1-YFP and CFP-Stim1 expressing HEK293 cells treated with scr (black) or STX11 (red) shRNA. (**O-Q**) Representative TIRF images of GFP-Mapper in store-depleted scramble and STX11 shRNA treated HEK293 cells (**O**) and quantification of intensity (**P**) and area of GFP-MAPPER puncta (**Q**). n=15.

Quantification of Orai1 intensities confirmed a partial defect in entrapment with relatively lower fraction of Orai1 present inside Stim1 puncta in STX11 depleted cells (Figure 7E-F). To determine whether reduced clustering of Orai1 resulted from structural defects within ER-PM junctions, we reversed the direction of fluorescent tags on Orai1 as well as Stim1 and repeated the experiment from Figure 7A-F in Figure 7G-L, using TIRF microscopy. Interestingly, the defect in Stim1 mediated entrapment of Orai1 could be rescued (Figure 7G-L) but SOCE remained significantly inhibited despite co-overexpression of Orai and Stim (Figure 7M-N) [33] [34] [35], suggesting ‘nonfunctional’ co-clustering of Orai1 and Stim1. We further assessed binding of full length YFP-Stim1 to mutant R289A_E272A_E275A_E278A Orai1-CFP and found that Stim1 was able to co-cluster with mutant Orai1 (Figure 7-figure supplement 1A-D).

Stim1 can itself serve as a marker of ER-PM junctions in store-depleted cells and quantification of area and mean intensity of Stim1-YFP or CFP-Stim1 puncta showed no defect in STX11 depleted cells imaged using confocal (Figure 7C,D) or TIRF microscopy (Figure 7H,I). To still assess whether STX11 depletion altered the structure and/ or proximity of junctional ER to PM, we expressed an alternate marker of ER-PM junctions, GFP-MAPPER, in STX11 depleted cells [36]. Quantification of the area and intensity of GFP-MAPPER puncta imaged using TIRF microscopy also did not show any significant difference in store-depleted, STX11 shRNA treated HEK293 cells (Figure 7O-Q). Collectively, these data show that STX11 binding to resting Orai1 has implications for CRAC channel function beyond mere co-entrapment by Stim1.

Defects in ‘functional’ entrapment of Orai1 suggest absence of a necessary molecular transition within Orai1 which precedes its interaction and gating by Stim1. To test whether STX11 induced structural changes within CRAC channel pore subunit, Orai1, we co-expressed Orai1-CFP along with Orai1-YFP in scramble control or STX11 depleted HEK293 cells and measured Orai:Orai FRET. We found significantly higher Orai1:Orai1 FRET in STX11 depleted resting cells in the absence of ectopically expressed Stim1 (Figure 8A). We have previously shown that store-depletion induces on-site assembly of Orai1 dimers into oligomers [14]. Notably, store-depletion induced a similar level of increase in Orai1:Orai1 FRET in scramble control cells. To confirm these findings, we performed acceptor photobleaching analysis on Orai1-CFP (donor), Orai1-YFP (acceptor) co-expressing HEK293 resting cells where we had stably knocked down the expression of endogenous Orai1 and Stim1. Although the trends were largely similar, the differences in Orai1:Orai1 FRET were much more significant in these cells (Figure 8B-C). These data demonstrate a previously unsuspected, novel molecular transition in Orai1 necessary for CRAC channel activation, which we define as ‘priming’. Further, we show that priming precedes the interaction of Orai1 with Stim1 in ER-PM junctions. To determine whether ‘priming’ simply induced a structural change in Orai1 protomers or facilitated correct assembly of Orai1 multimers, we expressed Flag-Orai1 in control or STX11 depleted resting cells, mechanically lysed to isolate membranes, crosslinked Orai1 oligomers using BS3 and resolved using SDS-PAGE (Figure 8D) (Figure 8-figure supplement 1). We found a greater percentage of higher order oligomers in STX11 depleted cells suggesting stoichiometric changes (Figure 8E).

**Figure 8:**
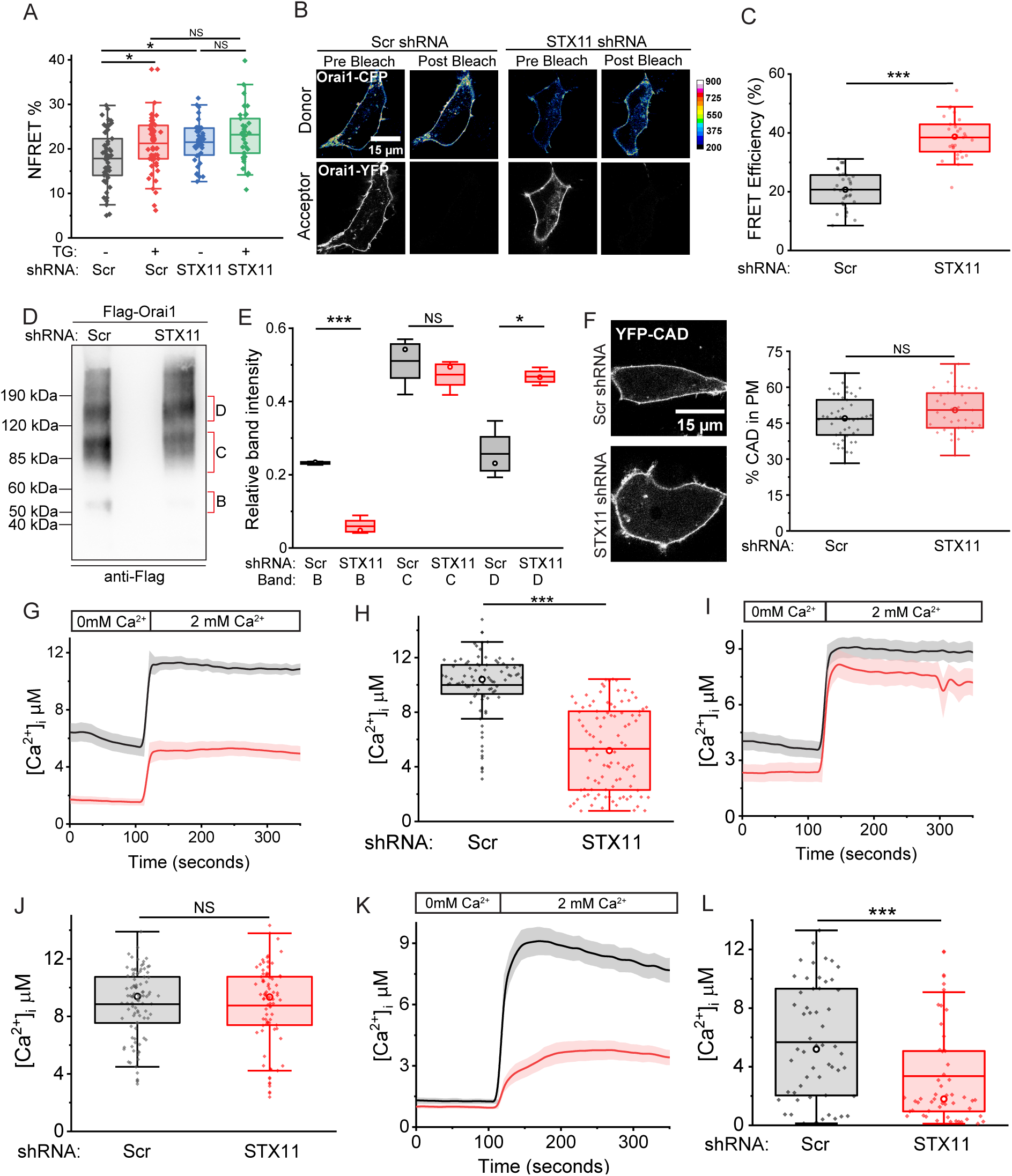
STX11 primes Orai1 for optimal on-site assembly. Oligomerization of Orai1. (**A**) N-FRET analysis of scramble and STX11 shRNA treated HEK293 cells co-expressing Orai1-CFP and Orai1-YFP before and after store-depletion. **(B-C)** Representative images (**B**) and quantification (**C**) of FRET analysis using acceptor photobleaching of Scr and STX11 shRNA treated Orai1, Stim1 double knockdown resting HEK293 cells co-expressing Orai1-CFP (donor) and Orai1-YFP (acceptor). **(D-E)** Representative western blot (**D**) and quantification of band intensities from repeats (**E**) showing distribution of BS3 crosslinked Flag-Orai1 oligomeric bands in control and STX11 depleted HEK293 cells. N=3. (**F**) Representative confocal image and quantification (right) of YFP-CAD localization in Scr and STX11 shRNA treated HEK293 cells expressing Orai1-CFP. **(G)** Fura-2 calcium imaging assay (**G**) and quantification of repeats (**H**) to measure constitutive calcium influx in Orai1-CFP and YFP-CAD expressing, control (black) or STX11 depleted (red) HEK293 cells. n=80-90, N=3 (**I-J**) Fura-2 calcium imaging assay (**I**) and quantification of repeats (**J**) to measure constitutive calcium influx in Orai1-H134S mutant expressing control (black) and STX11 depleted (red) HEK293 cells. n=60-70, N=3 (**K-L**) Fura-2 calcium imaging assay (**K**) and quantification of repeats (**L**) to measure constitutive calcium influx in Orai1-ANSGA mutant expressing control (black) and STX11 depleted (red) HEK293 cells. n=50-60, N=3

Increasing the density of Stim1 has been earlier proposed to bypass the requirements for accessory factors in SOCE. To determine whether increasing the levels of Stim1 could compensate for STX11 depletion, we expressed Orai1 tethered to two Stim-Orai activating region (SOAR) domains of Stim1 and GFP (Orai1-S-S-GFP) (Figure 8-figure supplement 2A-B), which has been previously shown to constitutively activate Orai1 in HEK293 cells [31]. In STX11-depleted Orai1-S-S-GFP expressing cells, the magnitude of calcium entry was significantly smaller (Figure 8-figure supplement 2C-D). To determine whether reduced calcium entry resulted from a defect in the ability of SOAR to bind Orai1, we expressed YFP-CAD, in control or STX11 depleted Orai1-CFP expressing HEK293 cell line (Figure 8F). Quantification of PM localized YFP-CAD as a fraction of total CAD showed no defect in STX11 depleted cells. Yet, as seen with Orai1-S-S-GFP, calcium entry was significantly reduced (Figure 8G-H). We next expressed a constitutively active H134S mutant of Orai1 (Figure 8-figure supplement 3A) as it harbors an open pore and the C-terminal cytosolic tails of Orai1 are unlatched, pointing towards the cytosol [37]. Remarkably, expression of H134S Orai1 completely restored constitutive calcium influx in STX11 depleted cells (Figure 8I-J) and showed similar PM expression when compared to wildtype Orai1 in all the groups (Figure 8-figure supplement 3B). To further establish whether binding with STX11 induces straightening of Orai1 tails or molecular transitions within transmembrane helices, we expressed in STX11 depleted cells the constitutively active ANSGA (261-265) mutant of Orai1 (Figure 8-figure supplement 3C), which harbors 4 consecutive mutations in the nexus region close to the Orai1 C-terminus resulting in the straightening of the C-terminal tail along with constitutive activation of CRAC currents [38]. The constitutive calcium entry in STX11 deficient Orai1 ANSGA mutant expressing cells was significantly suppressed (Figure 8K-L) even though the PM expression of the ANSGA Orai1 mutant was unchanged in STX11 depleted cells (Figure 8-figure supplement 3D). Taken together, these data show that STX11 induces a molecular switch in the pore-forming subunit, Orai1, which precedes and primes it for optimal on-site assembly into multimers and subsequent gating by Stim1. Such SNARE dependent priming of Orai1 is crucial and cannot be compensated by merely increasing the density of Stim proteins bound to Orai or by introducing mutations that alter the orientation of Orai1 tails. Figure 9A and Figure 9B show the proposed models depicting unprimed and primed Orai1 and its role in T cell function.

**Figure 9:**
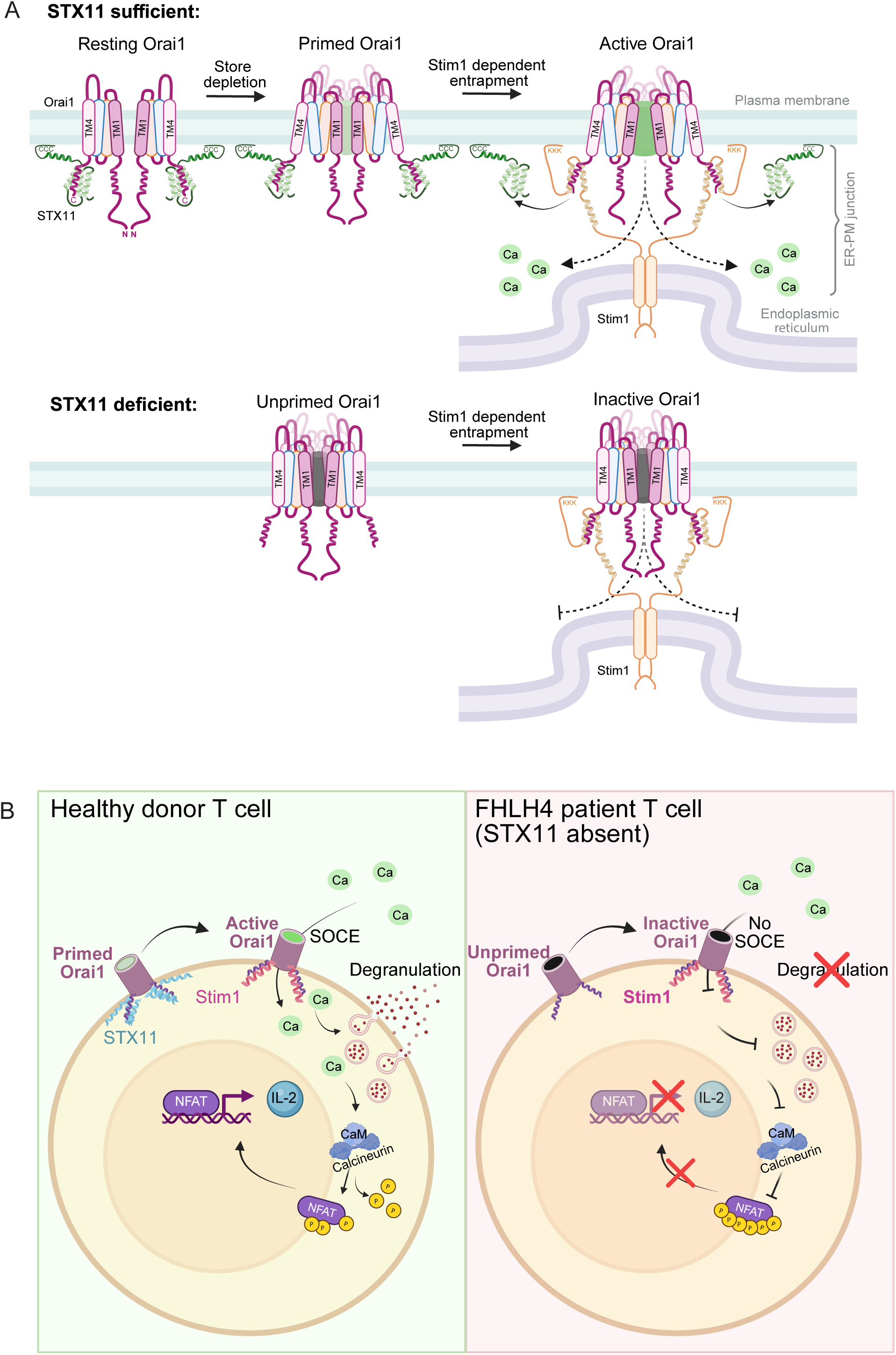
Models depicting STX11 mediated priming of Orai1 and its role in T cell signaling and function. (**A**) STX11 unbound (unprimed) and bound (primed) Orai1 in resting and store-depleted cells. For simplicity, STX11 interaction with only two subunits of Orai1 has been shown. (**B**) Regulation of T cell SOCE, NFAT activation and effector function by STX11.

FHLH is a heterogeneous immune disorder where diverse genes such as *Prf1*, *STX11*, *UNC13D*, *Munc18-2*, *MAGT1, XIAP, LYST, AP3B1, RAB27A*, *ITK*, amongst several others, are broadly grouped together based on disease symptoms, although the age of onset and severity varies across the spectrum [39, 40]. Congenital immunodeficiency involving defects in the CTL and NK cell cytotoxicity appear to be the common factors underlying the development of FHLH, although the individual pathways involved are highly diverse. Using shRNA, we screened 6 additional genes, *Perforin1*, *RAB27A*, *UNC13D* and *Munc18-2*, *Synaptotagmin11* and *13*, that have been either associated with vesicle fusion or routinely grouped together with STX11 in diagnosing FHLH. SOCE was not affected by depletion of most genes (Figure 1-figure supplement 3). Future studies will evaluate the role of others.

## Discussion

CRAC channels conduct a small, highly specific calcium current in response to the depletion of intracellular calcium stores. According to the prevailing view, Orai multimers reside in the PM and depend on the ER resident store sensor, Stim, for all structural transitions leading up to their activation [10]. Unlike other ion channels, the process of activation of CRAC currents is exceptionally slow. For instance, following break-in or store-depletion, on average, it takes ∼5 minutes for Stims to cluster in the ER-PM junctions and for measurable CRAC currents to flow from Orai1. During this time, Stims undergo intramolecular transitions and slowly trap and gate freely diffusing Orai. It is, therefore, believed that no molecular transitions take place within the pore forming subunit, Orai, until its entrapment by Stim in the ER-PM junctions.

We have shown that in STX11 deficient cells, the basal Orai:Orai FRET is significantly higher even in the absence of Stim1 or store-depletion. These data establish that Orai1 undergoes a, hitherto unrecognized, SNARE dependent molecular transition which precedes trapping and gating by Stim1. We have earlier shown that store-depletion induces dynamic on-site assembly of Orai1 dimers into oligomers [14]. In STX11 depleted resting cells, the ∼25-50% increase over basal Orai:Orai FRET largely results from a change in the oligomerization of Orai1 because the proportion of BS3 crosslinked Orai1 oligomers was also found to be shifted towards higher molecular weight. It is very likely that a change in the overall conformation of Orai1 protomers facilitated this change in store-dependent Orai assembly and contributed to the overall change in Orai:Orai FRET. This conclusion is also supported by the observation that STX11 depletion induced change in Orai:Orai FRET which not only persisted but increased in Orai1, Stim1 double deficient cells.

Furthermore, STX11 bound a region of Orai1 C-terminus that has been previously shown to bind Stim1. Therefore, STX11 induced molecular transitions in Orai are independent of and precede its association and clustering by Stims. Because tagging of STX11 with fluorescent tags on either end results in mis-localization and degradation, our study could not demonstrate the real-time removal of STX11 from the Orai1-C-terminus to allow for Stim1 binding in live cells. It is also unclear whether binding or removal of STX11 induced the proposed molecular transition defined as ‘priming’ of Orai1. Further, it remains to be determined whether STX11 removal is actively induced by an additional chaperone or passive displacement due to formation of relatively higher avidity oligomeric associations between Orai1:Stim1 in ER-PM junctions. Future studies will be able to systematically evaluate each of these possibilities.

We have shown that the C-terminal cytosolic tail of Orai1 directly couples with STX11 in resting cells. Although binding appeared reasonably strong in co-IPs, STX11:Orai1 interaction appeared somewhat weak in pull-downs. This is likely due, in part, to the instability of purified His-STX11 in solution which necessitated shorter co-incubation times during pull-down assays to prevent precipitation of STX11. Instability could arise from the absence of a stabilizing co-chaperone, such as one of the known binding partners of SNAREs [41] [42] or membranes to which STX11 has been proposed to attach in live cells. Together, these factors appear crucial for the stabilization of purified STX11 *in vitro*.

The individual relevance of Orai1 N-*versus* C-terminus in the trapping *versus* gating of Orai1 remains unclear [43]. Though crucial for trapping and gating, Orai1 tails were truncated in the early structures of *Drosophila* Orai [37]. A previous NMR structure of truncated Orai1 C-terminal tails suggested that the tails of two adjacent Orai1 subunits, bend, pair with each other in an antiparallel fashion and sit closely apposed to PM [44]. However, in recent structures of constitutively active H134 mutant Orai, the C-terminal tails were found to orient away from the membrane [37]. Although C-term tails in ANSGA Orai1 mutant are also proposed to be unlatched, STX11 depletion inhibited ANSGA Orai1 induced constitutive calcium influx [38]. In the absence of direct structural evidence that establishes conformational similarity between constitutively active H134 *versus* ANSGA mutant Orai1, Orai1+CAD and Orai1-S-S, it is likely that Orai1 can assume several different open conformations/ assemblies [14, 37, 45–47]. Since most mutants showed sensitivity to STX11 depletion, it reinforces our conclusions that STX11 induced conformational shifts precede gating and involve Orai1 TMs.

Using Fura-2 imaging assays in STX11 depleted cells, we have conclusively established that STX11 is necessary as well as limiting not only in live and unperturbed HEK, U2OS, Jurkat and primary T cells but also HEK293 cells co-overexpressing Orai1 and Stim1. Previous whole cell patch clamp analysis of HEK293 cells co-overexpressing Orai1 and Stim1 showed up to 50-100-fold amplification of CRAC currents whereas Fura-2 based imaging of live cells routinely shows only 2-fold amplification [33] [34] [35]. The patch clamp studies performed on Stim, Orai over-expressing cells are often extrapolated to conclude that all other accessory factors that have since been identified are unnecessary for native CRAC currents. A major problem with this interpretation is that even though amplification of CRAC currents was shown, none of the previous patch clamp studies established whether higher currents resulted from a greater number of active channels or unchecked conductance per channel by performing single channel recordings. Direct single-channel recordings of CRAC are challenging. As a result, estimates of open probability and channel number often rely on fluctuation (noise) analysis rather than direct measurements of single-channel events [48]. The fact that recording of I_CRAC_ in the whole cell configuration involves use of 20mM extracellular calcium and dialysis of cells with high concentrations of BAPTA, a fast calcium chelator, the calcium dependent inactivation of CRAC channels, a key mechanism of regulation of I_CRAC_ magnitude in live cells is disrupted. Furthermore, hyperpolarizing potentials and voltage clamping used to record CRAC currents prevents the progressive depolarization of cells due to influx of cations which would otherwise dampen the driving force for CRAC currents in intact cells. Given these limitations, Fura-2 assays offer a more realistic and physiological reflection of SOCE amplitudes in live cells as they allow for these regulatory mechanisms, which are necessary for the regulation of native currents, to be still operational. While patch clamp is a direct method of verification of CRAC currents, exercising logic in the extrapolation of amplitudes of I_CRAC_ using patch clamp is imperative.

Most human mutations in the *STX11* coding region result in the complete loss or severe depletion of STX11 protein levels [28]. The FHLH4 disease, therefore, results from reduced STX11 expression in most reported patients. Targeted mutagenesis of the STX11:Orai1 interface lining residues based on the AlphaFold 3 model, however, didn’t destabilize or mis-localize either N147A_E150A STX11 or R289A_E272A_E275A_E278A Orai1 and yet both mutants showed reduced binding to their respective wildtype counterparts. Importantly, the R289A_E272A_E275A_E278A mutant of Orai1 retained the ability to bind and cluster with Stim1.

Most importantly, we have shown that deficiency of a Q-SNARE, STX11, causes a, vesicle trafficking independent, severe defect in SOCE, IL-2 gene expression and degranulation in FHLH4 T cells. Mechanistically, STX11 directly binds and induces a molecular switch in Orai1 necessary for its assembly into ‘functional’ multimers which we call a ‘primed’ state. We suggest that ion channel priming is a novel, direct and primary role of specific SNAREs, independent of vesicle trafficking. In accordance with this hypothesis, early SNARE-like proteins have been found in bacteria, which lack an endomembrane system but express a variety of ion channels [49–51]. SNAREs are now thought to have evolved from a common archaeal precursor found in the genomes of Asgard and members of *Legionella* [52].

Analyses of FHLH4 patient T cells suggest that SOCE defects significantly contribute to the pathogenesis of FHLH4 disease. We, therefore, propose that pharmacological induction or promotion of primed intermediate state of Orai1 could potentially provide an alternative to immunosuppressants and bone marrow transplants for FHLH4 as well as SCID patients, with minimal risk of autoimmunity.

## Materials and Methods

### Plasmid constructs and transfection

Stim1 CAD was sub-cloned in eYFPC1 vector after amplification from YFP-Stim1 plasmid to generate YFP-CAD. pLKO.1, psPAX2 and pMD2.G were purchased from Addgene. pLKO.1 cloned shRNA sequences targeting the genes of interest, were either purchased from Horizon Discovery, UK or designed and cloned in-house using the Broad Institute portal (https://portals.broadinstitute.org/gpp/public/gene/search). Full-length human STX11 was cloned from human cDNA prepared from HEK293 cells and subcloned into pcDNA3.1(+), pcDNA/4TO/Myc-HisA (Invitrogen, Grand Island, NY), pMSCV-IRES-mcherry (Addgene), pEF1alpha-IRES (Clontech) and pET28b vector [16] with an N-terminal 6XHis-tag. The fragments of STX11 were cloned in pET28b vector with an N-terminal 6xHis tag for expression in *E. coli*. HA tag was inserted between H30 and G31 in STX11 cloned in pMSCV-IRES-mCherry construct using PCR. H134S and ANSGA mutants of Orai1 were generated using site directed mutagenesis and PCR, respectively. The fragments of Orai1 (1-87, 1-47, 48-103, 256-301 and 272-292) were amplified from full length constructs and cloned into pMAL-c5X vector (New England Biolabs), in-frame with MBP protein coding sequence, as described previously [16]. SNAP23, SNAP25 and SNAP29 cDNA cloned in pCMV-Sport6.1 vectors were purchased from Dharmacon and subcloned into pcDNA4/TO/Myc-HisA. SNAP23 was subcloned in pMAL-c5X vector in-frame with MBP. All plasmid DNA transfections in human cell lines and primary cells were done using Lipofectamine 2000 (Invitrogen)/ Lipofectamine 3000 (Invitrogen) or Amaxa nucleofection kit (Lonza, Basel, Switzerland) respectively, as per manufacturer’s protocol.

### Cell Lines

HEK293-FT cell line was cultured in high glucose DMEM with 10% FBS and 10mM HEPES, 1X Penicillin Streptomycin, 1X GlutaMax and 1X non-essential amino acid and transfected with the appropriate plasmids to generate viral supernatants. Lentiviral shRNA transduction experiments were performed in HEK293, U2OS or Jurkat (ATCC, Manassas, VA) cell lines cultured in low glucose DMEM (Hyclone, Logan, UT) or RPMI (Hyclone) respectively, with 5-10% fetal bovine serum (Hyclone), 1X Penicillin Streptomycin, and GlutaMax (Gibco, Grand Island, NY). Stable cell lines generated using HEK293 and U2OS parent lines have been described previously [9]. To generate HEK293 stable cell line expressing Orai1-BBS-YFP construct, the cells were nucleofected (SE Cell Line 4D-Nucleofector X kit_Amaxa _V4XC) and YFP positive cells were sorted. All cell lines were tested for mycoplasma contamination twice every year and found to be negative.

### Lentiviral transductions

For the generation of lentiviral supernatants, shRNAs cloned in pLKO.1 were co-transfected with psPAX2 packaging and pMD2.G envelope plasmids into HEK293-FT cells using the calcium phosphate method of transfection. 48- and 72-hours post-transfection, viral supernatants were collected, pooled and stored at −80°C till further use. For transduction of HEK293 and Jurkat cells, 0.1 million cells were plated in 6- or 24-well plates either the day before (HEK293) or the same day (Jurkat). Viral supernatants were added to the cells along with 8μg/ml polybrene and cells were spun at 2500 RPM, 30°C for 90 mins. 24 hours or 48 hours post-spinfection, Puromycin was added at a final concentration of 1μg/ml to HEK293 and Jurkat cells, respectively. Cells were analyzed 3-5 days post transduction.

### Immunocytochemistry and confocal imaging

For STX11 localization, resting or store-depleted (1μM Thapsigargin) WT HEK293 or HEK293 cells stably expressing Orai1-YFP were either plated in 6 well plates and spinfected with viral supernatants generated from pMSCV-STX11(HA)-IRES-mCherry, pMSCV-IRES-mCherry (empty vector control) or plated in carbon coated, glow discharged 35mm glass bottom dishes (IBDI) and transfected with pEF1alpha-STX11-IRES-mCherry. 24 hr post-spinfection, cells were plated in 35mm glass bottom dishes. For immunolabelling, cells were washed with Ringer’s buffer, fixed with 4% PFA and blocked with 3% BSA containing 0.1% NP40 for 1.5 hrs. Post-fixation and permeabilization, cells were incubated either with anti-HA (CST) or rabbit anti-Stx11 (SySy/ Proteintech) primary antibodies O/N at 4°C, washed and incubated with either Alpaca VHH anti-rabbit AF488, donkey anti-rabbit A647 or goat anti-rabbit A647 secondary antibodies for 1 hour at room temperature (RT). Cells were counter stained with DAPI for 10 min at RT and imaged using Olympus FV3000 laser scanning confocal microscope. Images were acquired sequentially using the following parameters: DAPI (DM405/488 dichroic, 405nm excitation, 430-470nm emission); anti-HA Alexa488 (DM405/488, 488nm excitation, 500-590nm emission); STX11 (DM405/488/561/640, 640nm excitation, 650-750nm emission); Orai1-YFP (DM405/488/561/640, 488nm excitation, 521-591nm emission); CFP-Stim1 (DM405-445/514/594, 405nm excitation, 448-510nm emission). For STX11-N147A-E150A mutant, HEK293 cells were spinfected with viral supernatants and imaged as described above where WGA (Wheat germ agglutinin) conjugated with Alexa 488 was used to mark plasma membrane. For immunolabelling of ER and Golgi, HEK293 cells were treated with Scr or STX11 shRNA, washed, fixed, blocked and incubated with anti-KDEL and anti-GM130 primary antibodies overnight at 4°C. Following incubation, the cells were washed with 1X PBS containing 0.1% NP40, incubated with Donkey anti-mouse A647 secondary antibody for 30 mins at RT, washed and counter-stained with DAPI for 10 mins. Images were acquired using Olympus FV3000 laser scanning microscope as mentioned above.

### Quantification of Orai1 and Stim1 intensities inside/outside ER-PM puncta

HEK293 cells stably or transiently expressing wildtype Orai1-YFP or E272A-E275A-E278A-E289A Orai1-YFP with CFP-Stim1 or CFP-Orai1 with Stim1-YFP were plated in 6 well plates and transduced with scramble or STX11 shRNA. 48hr post transduction, cells were trypsinized and plated in carbon coated 35mm dishes. Images were acquired using a TIRF microscope setup described before ^14^. Briefly, resting cells were imaged first and positions were marked. Store-depleted images were acquired after incubation with 1 μM Thapsigargin (TG) and 10mM EGTA for ∼6 mins. Stim1 and Orai1 clusters were defined using Otsu local thresholding and Pearson’s correlation coefficient was measured using colocalization finder in ImageJ software. Stim1 and Orai1 intensity (AU), area in µm^2^ and Pearson’s correlation coefficient values values obtained from the analysis were plotted using Origin software.

### Orai:Orai FRET

Scramble and STX11 shRNA transduced HEK293 cells were plated on freshly carbon coated, glow discharged 35mm glass bottom dishes (IBDI) and transfected with 0.5 μg each of Orai1-CFP and Orai1-YPF plasmid DNA using Lipofectamine 3000. 16 hours post transfection, co-transfected cells were imaged using PlanApoN 60XO/1.42 NA objective with Olympus FV3000 laser scanning microscope. Orai1-CFP, Orai1-YFP and FRET images were acquired with the following excitation laser and emission bandwidth parameters using DM405-445/514 dichroic mirror: CFP Ex-405nm Em-460-630nm; YFP Ex-514nm Em-530-630nm; FRET Ex-405nm, Em-530-630nm. ROIs representing the plasma membrane rim were drawn around each cell, followed by local thresholding using Bernsen’s algorithm in Fiji software to generate an image mask which was applied to CFP, YFP, FRET images to extract mean intensity values. The background subtracted mean intensities of FRET (I_FRET_), CFP (I_Donor_) and YFP(I_acceptor_) images were measured for each cell. Cells singly expressing Orai1-CFP or Orai1-YFP only were used as CFP donor or YFP acceptor channel bleed-through (BT) controls. The BT was calculated using the following formula:

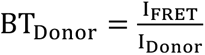 when the Orai1-CFP donor alone is expressed,

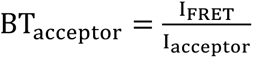 when the Orai1-YFP acceptor alone is expressed.

FRET and NFRET percentage was calculated by subtracting the contributions of CFP and YFP bleed through from the FRET signal using the following:

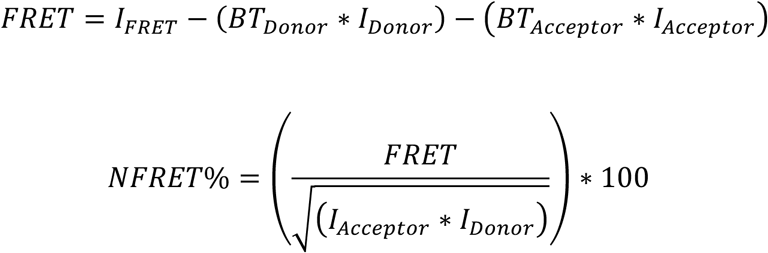

### Acceptor Photobleaching

Scramble and STX11 shRNA transduced, Orai1, Stim1 double knockdown HEK293 cells where were plated on Poly-D-Lysine coated glass bottom dishes (Cellvis) and transfected with 0.5 ug DNA each of Orai1-CFP and Orai1-YPF using Lipofectamine 2000. 16 hours post transfection, co-transfected cells were imaged using PlanApoN 60XO/1.42 NA objective with Olympus FV3000 laser scanning microscope. Images were acquired with the following excitation laser and emission bandwidth parameters using DM405-445/514 dichroic mirror: CFP Ex-405nm Em-460-508nm; YFP Ex-514nm Em-530-630nm, with 4% offset. The pre bleach images of Orai1-CFP (the donor) and Orai1-YFP (the acceptor) were captured, sequentially. This was followed by bleaching of the acceptor, Orai1-YFP, using 100% 514 laser power for 2 mins and then acquiring the donor and acceptor images post bleach. ROIs were drawn around the cell periphery using the Orai1-CFP image. Local thresholding was performed using Bernsen’s algorithm to generate binary masks, which were then applied to the CFP images to measure mean intensities. The background subtracted mean intensities of CFP images were measured for each cell and the FRET efficiencies were calculated as follows:

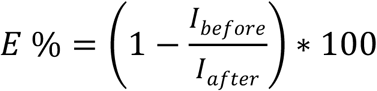

I_before_ and I_after_ denotes the average fluorescence intensity of the donor in the plasma membrane before and after acceptor photobleaching.

### Alpha-bungarotoxin binding assay

U2OS or HEK293 cell lines stably expressing Orai1-BBS-YFP construct were either transduced with scramble or STX11 shRNA and harvested 3 days post-transduction. Jurkat cells treated with Scr and STX11 shRNA were transiently transduced with pMSCV-Orai1-BBS-YFP 48 hrs prior to harvest. For surface labeling, cells were incubated on ice with AF647 conjugated alpha-bungarotoxin (BTX-A647) (1μg/ml) dissolved in 1X HBSS with 2% FBS 30 mins, washed and fixed with 4% PFA (∼20 min at RT). For flow-cytometry, U2OS and HEK293 cells were analyzed on Cytoflex FACS analyzer (Beckman Coulter) and data were analyzed using Flow Jo software. For microscopy, the shRNA treated HEK293 cells were plated in 35mm glass-bottom dishes, one day prior to harvest, whereas labeled Jurkat cells were plated on poly-D-lysine coated coverslips, mounted onto glass slides and sealed. Imaging was performed using Olympus FV3000 confocal microscope.

Line profiles of Orai1-YFP and BTX-A647 were drawn on raw confocal images of HEK293 cells. Quantifications of Orai1-YFP and BTX-A647 were calculated using the following formula:

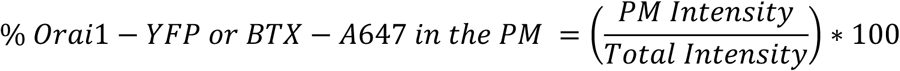

### Alkaline phosphatase (AP) assay

HEK293 cells were transduced with Scr and STX11 shRNA and 48 hrs post-transduction the cells were transfected with pAP tag plasmid (Genhunter) in low glucose DMEM without phenol red. Supernatants from control and transfected cells were collected at 48hour post transfection and triplicates of undiluted supernatant or those diluted at 1:1, 1:2,1:4 using DMEM without phenol red were added to 96-well plates and mixed in 1:1 ratio with 1 mg/ml pNPP (Disodium-4-Nitrophenyl Phosphate Hexahydrate) substrate in AP Assay Buffer (100 mM Tris-Cl pH-9.0, 100 mM NaCl and 5 mM MgCl2) to obtain final AP:pNPP ratios of 1:1, 1:2, 1:4 and 1:8. Supernatants from untransfected cells were used as blank controls. After 25 mins of incubation at 37°C in the dark, the assay was terminated by adding 0.4M NaOH and absorbance at 405 nm was measured using Softmax Pro version 5.4 in FlexStation 3.

### Measurement of GFP-Mapper cluster size and intensity

HEK293 cells treated with Scr and STX11 shRNA were transfected with GFP-Mapper plasmid, 48 hrs post-transduction. The cells were store-depleted with 1 μM Thapsigargin (TG) and 10mM EGTA for ∼6 mins, fixed with 4% PFA, washed and imaged using TIRF microscopy. To analyze the images, local thresholding was performed in ImageJ using the Bernsen algorithm, and the resulting mean intensity and area (μm^2^) of the GFP-Mapper clusters were plotted using Origin software.

### Validation of STX11 knockdown using qPCR

Scramble and STX11 shRNA treated HEK293 and Jurkat T cells were counted and equal number of cells were used for lysis. The lysates were homogenized using QIA shredder columns (Qiagen) and RNA isolation was done using RNeasy Plus Mini kit (Qiagen). Total RNA isolated from both the groups was used for cDNA synthesis using oligo dT primers and Superscript IV as per manufacturer’s guidelines. The concentration of synthesized cDNA was estimated using Qubit ssDNA assay kit and STX11 TaqMan probes were used to perform the qPCR using LightCycler 96 (Roche). Beta-actin and RPL30 were used as housekeeping control.

### Single cell Ca^2+^ imaging

Cells were plated in carbon-coated glass bottom dishes (one day prior or 30 mins before the assay for HEK293, U2OS and Jurkat T cells respectively) and loaded with 1μM Fura-2 AM dye in HBSS (CaCl_2_ 1.8mM, KCl 5.36mM, MgSO_4_ 0.81mM, NaCl 136.89 mM, Na_2_HPO4 0.335mM, D-Glucose 5.55mM) for 30 mins, at 37°C in the dark. After incubation, cells were washed and incubated in Ringer’s buffer (135mM NaCl, 5mM KCl, 1.8mM CaCl_2_, 1mM MgCl_2_, 5.6mM Glucose, 10mM HEPES) for an additional 10 mins, washed and imaged in Ringer’s buffer or Calcium-free Ringer’s buffer (135mM NaCl, 5mM KCl, 1mM MgCl_2_, 5.6mM Glucose, 10mM HEPES at pH7.5), as indicated. Olympus IX-71 inverted microscope equipped with a Lamda-LS illuminator (Sutter Instrument), Fura-2 (340/380) filter set (Chroma, Bellows Falls, VT), a 10X 0.3NA objective lens (Olympus, UPLFLN), and a Photometrics Coolsnap HQ2 CCD camera was used to capture images at a frequency of ∼1 image pair every 2 or 4 seconds interval. Data were acquired, analyzed and plotted using MetaFluor (Molecular Devices), Microsoft Excel, and Origin softwares. Approximately 30-40 cells were imaged per group in each experiment unless otherwise stated. *In-vitro* calibration of Fura-2 was performed using calibration kit (Molecular Probes_F6774) according to the manufacturer’s protocol. Briefly, calcium calibration buffers spanning a range of free calcium concentrations (0, 0.017, 0.038, 0.065, 0.1, 0.15, 0.225, 0.351, 0.602, 1.35, and 39 µM) and containing 15 µM microsphere beads were loaded onto cleaned glass slides, sealed with a coverslip and imaged. A standard linear calibration curve was constructed by plotting the 340/380 ratios (y-axis) against the known free calcium concentrations (x-axis) and used to quantify the absolute intracellular free calcium concentrations using simple linear regression equation y=mx+c.

### Quantification of SOCE

Quantification of mean SOCE ± SE from three independent experiments was performed as follows. The mean of 5 peak amplitude values after the addition of 2mM calcium from one of the scramble shRNA groups, amongst the three experiments, was normalized to the baseline values (5 values just before the addition of 2mM calcium) and was set at 100%. The relative percent responses of all other Scr and test shRNA groups were normalized with respect to this group.

### Constitutive calcium influx assay

For constitutive calcium influx assay, shRNA treated HEK293 cells were either transfected or nucleofected with Orai1-S-S-EGFP, Orai1(H134S)-YFP or Orai1(ANSGA)-YFP and ∼12-14 hour post transfection (∼4-5 hours post nucleofection), cells were loaded with Fura-2 AM, washed and imaged in Ringer’s buffer with 0 mM calcium to acquire baselines and 2 mM calcium thereafter. To measure CAD induced constitutive calcium influx, Orai1-CFP expressing stable HEK293 cells were nucleofected with YFP-CAD and analyzed ∼4 hour later as described above. To measure CAD induced calcium influx in E272A-E275A-E278A-R289A mutant Orai1-YFP expressing cells, Orai1-shRNA treated HEK293 cells were transfected with wildtype or mutant Orai1-YFP constructs along with YFP-CAD. To identify cells expressing Orai1 mutants and fusion proteins in the plasma membrane, cells were imaged using a 20X 0.7NA water objective lens (Olympus, UApoN340, Japan). Images were acquired at a frequency of ∼1 image pair every 10 seconds interval to avoid photobleaching and analyzed as described above. Quantification of constitutive calcium influx using Orai1 mutants was done by plotting the average of five consecutive data points immediately following addition of 2mM calcium across three independent repeats. Fura 2 calibration was done separately for these experiments as described above.

### Transferrin recycling assay

HEK293 and Jurkat T cells treated with Scr or STX11 shRNA were plated on 35-mm glass-bottom dishes and washed three times with serum-free DMEM to remove residual FBS. The cells were then incubated with Alexa Fluor 647-conjugated human holo transferrin (20ug/ml) in serum-free DMEM, for 30 min at 37°C (pulse), placed on ice for 5 min and subjected to three rapid washes with acidified ice-cold serum-free DMEM (pH 3.5-5.5) to remove transferrin bound to cell surface receptors. Cells were then chased for indicated time points at 37°C, washed with HBSS, fixed with 4% PFA, washed again with PBS and imaged using an IX83 widefield microscope equipped with a 60X/1.25 NA objective lens (Olympus UPFLN). Fluorescence images were collected on a Hamamatsu Orca-Fusion camera using the same settings across all conditions. Brightfield images were used to define cell boundaries, and per-cell mean fluorescence intensities were quantified using ImageJ software.

### Dextran-FITC uptake assay

HEK293 cells treated with Scr or STX11 shRNA were plated on 35-mm glass-bottom dishes, washed with serum-free DMEM and incubated with FITC-conjugated dextran (1mg/ml) in serum-free DMEM, for 30 min at 37°C. The cells were then washed with HBSS, fixed with 4% PFA and imaged. To preserve FITC fluorescence post-fixation, the cells were treated with 20mM ammonium chloride prior to imaging. Images were acquired as described above for Transferrin recycling assay.

### Measurement of NFAT nuclear localization by Western blot

Jurkat T cells were transduced with scramble or STX11 shRNA to knock-down STX11 as described above. On day 4 post-transduction, cells were collected, spun down and resuspended in plain RPMI media and rested for 1hr at 37°C. Following this, cells were counted and divided into two equal groups. One group was resuspended in RPMI (unstimulated) and the other in RPMI media containing 1 μM TG + 10 ng/ml PMA (Phorbol Myristate Acetate) (stimulated) and incubated at 37°C for 30 mins. Following incubation, cells were pelleted and the nuclear and cytosolic protein fractions were separated using NE-PER kit according to the manufacturer’s guidelines and subjected to SDS-PAGE and western blot using the mouse anti-NFATc2 primary antibody (Santa Cruz) followed by Donkey anti-mouse secondary antibody.

### Estimation of nuclear translocation of NFAT by immunolabeling

Control and STX11 depleted Jurkat T cells were rested in plain RPMI for ∼1 hour prior to the assay. ∼100,000 cells per group were plated on freshly carbon-coated coverslips for 40 minutes and stimulated for 1 hour at 37°C with 5 μg/ml anti-CD3 antibody, diluted in plain RPMI. Following this, cells were washed and fixed using 4% PFA diluted in 1X PBS for 20 minutes at RT, washed twice and incubated with 30 mM Glycine for 10 minutes at RT. For permeabilization and blocking, cells were incubated with 0.1% Saponin, 3% BSA diluted in 1X PBS for 1 hour at RT, washed and incubated with anti-NFAT primary antibody (anti-NFAT1, CST) at 4°C, overnight. Following primary antibody application, cells were washed with 1X PBS and incubated with anti-Rabbit AF647 secondary antibody for 1 hour, washed and stained with DAPI for 5 minutes followed by additional washes. Images were acquired in the FV3000 confocal microscope using a 100X objective lens. DAPI was used to identify the nuclear area. Nuclear *versus* whole cell (total) NFAT mean intensity ratio was plotted across different groups.

### Whole cell lysates (WCLs), Western blot and Co-immunoprecipitation (Co-IP)

HEK293 cells transfected with the desired plasmids were lysed using buffer containing 50 mM Tris-Cl (pH 8.0), 150 mM NaCl, 1% NP-40, 1 mM PMSF, and protease inhibitor cocktail. The whole cell lysates were centrifuged at 21000 g for 15 minutes and supernatants were subjected to SDS-PAGE, and western blot. For immunoprecipitations, lysates were divided into two equal parts. To the first tube, the appropriate anti-mouse or anti-rabbit primary antibody was added and to the second, same amount of the respective IgG control antibody was added. The antibody-lysate mixtures were incubated overnight at 4°C. Following this, Protein A/G Mag Sepharose beads were added and incubated with the antibody-lysate mixtures for 4 hours, washed with the lysis buffer containing 0.1% NP-40, boiled with 1X Laemmli buffer + 120 mM DTT and subjected to SDS-PAGE and western blot analysis. Typically, 1/10^th^ of the whole cell lysate (WCL) was loaded in the input lane of the co-IP blots. The full-length blots pertaining to each figure have been deposited in Zenodo https://doi.org/10.5281/zenodo.21663510

### *E. coli* expression and *in-vitro* binding assays

Full-length His_6_-tagged Stx11 and truncation mutants were cloned in pET28b, expressed in Lemo21 (DE3) *E. coli* cells and induced with 1 mM IPTG (Isopropyl β-d-1-thiogalactopyranoside) for 14-18 hours at 18°C. The cell pellets were lysed in buffer containing 50 mM Tris-Cl (pH 8.0), 150 mM NaCl, 10% Glycerol, 1 mM PMSF, 1% Sarkosyl, 0.1 mg/ml Lysozyme, protease inhibitor cocktail and sonicated on ice. DNase I was added after sonication and the lysates were further incubated for ∼60 minutes before centrifugation at 21000 g for 40 mins. The supernatants were subjected to SDS-PAGE to confirm expression by Coomassie staining and subsequently used for pull-down assays. MBP-tagged Orai1 constructs were expressed in Lemo21 (DE3) *E. coli* cells, induced with 0.3mM IPTG (Isopropyl β-d-1-thiogalactopyranoside) for 14-18 hours at 18°C. The cell pellets were lysed in buffer containing 50 mM Tris-Cl (pH 8.0), 150 mM NaCl, 5% Glycerol, 1 mM PMSF, 0.1 mg/mL Lysozyme and protease inhibitor cocktail (1:200) and sonicated on ice. DNase I was added after sonication and the lysates were further incubated for ∼60 minutes before centrifugation at 21000 g for 40 mins. The supernatants were collected and subjected to SDS-PAGE to confirm expression by Coomassie staining and used for pull-down assays. Following lysis, the His_6_-tagged Stx11 proteins were purified using Talon Beads and MBP-tagged Orai1 proteins using Dextrin Sepharose/Amylose resin according to the manufacturer’s guidelines. For *in-vitro* binding assays, lysates prepared from cells expressing MBP or MBP-tagged Orai1 fragments were incubated with 25 μL Dextrin Sepharose Beads and incubated for 2 hours at 4°C. After incubation, beads were washed thrice and ∼50-125 ng of purified His-tagged full length Stx11, His-tagged SNARE or His-tagged H_abc_ domains were diluted in buffer containing 50mM Tris-Cl (pH8), 150 mM NaCl, 5% Glycerol and 0.1% NP-40 and added to the beads. Following 1 hour of incubation at 4°C, the beads were washed thrice, re-suspended and boiled in the binding buffer containing 1X Laemmli and 120 mM DTT. The eluate was subjected to SDS-PAGE and Western blot. MBP and MBP-tagged proteins were detected with Ponceau staining, Full length STX11 using Rabbit anti-STX11 primary antibody (Thermo) followed by Donkey anti-rabbit HRP and His-tagged STX11 fragments were detected or mouse anti-6XHis primary antibody (Invitrogen) followed by Donkey anti-mouse HRP secondary antibody. Densitometric quantification of the relative band intensities was performed using ImageJ software and plotted in Origin software. To control for variation in experimental repeats, the band intensity of the control complex for wild-type (MBP-Orai1-WT:His-STX11-WT) for individual repeats was set to 100 percent and relative intensities of bands from other test groups were calculated with respect to this.

### BS3 crosslinking of Flag-Orai oligomers

Scramble and STX11 shRNA transduced HEK 293 cells were transfected with Flag-Orai1, harvested, washed, pelleted and subjected to one cycle of snap freeze-thaw. The pellet was resuspended in ice cold lysis buffer (1X HBSS, 1mM PMSF and protease inhibitor cocktail), sonicated for 30 secs (Branson 2510) and mechanically homogenized using dounce homogenizer. The homogenized supernatant was spun at 1500 x g for 10 mins at 4° C, to pellet the cell debris. The supernatant was ultracentrifuged at 105,000 x g for 1 hour at 4° C (Optima MAX-XP ultracentrifuge, Beckman Coulter). The membrane fraction pellet was resuspended in lysis buffer and re-homogenized using dounce homogenizer. The protein concentration was estimated using BCA assay and BS3 cross-linker was added to the resuspended membranes, at a final concentration of 1mM, and incubated for 30 mins at room temperature. BS3 was quenched using 50mM Tris-HCl pH 7.5 for 15 mins at room temperature, followed by addition of 1X Laemmli and 100 mM DTT and incubation at 37° C for 10 mins. The samples were resolved using 4-15% gradient SDS-PAGE and subjected to Western blot using anti-Flag antibody.

### Quantification of Flag-Orai oligomers using Western blot

Equal size ROIs were drawn over corresponding protein bands of same size, using ImageJ. The total intensity of each band was measured by calculating the area under the intensity curve and its relative proportion was calculated by dividing the raw intensity of each band by the sum of intensities of all the bands in the respective lanes.

### qPCR to estimate gene expression in Jurkat T cells

Scramble (scr) and STX11 shRNA treated Jurkat T cells were rested for ∼1 hour prior to the assay, stimulated with soluble anti-CD3 (5-10 μg/ml) for 3 hours and washed with HBSS. Total RNA was isolated using RNeasy Mini kit as per manufacturer’s instructions. cDNA was synthesized using random hexamers and Invitrogen Superscript IV kit and quantified using Qubit ssDNA assay. Taqman probes for IL-2 and beta-actin housekeeping control were used for performing the qPCR in triplicates. Data analysis was done by calculating the double delta C_t_ values.

### EIA to measure IL-2 levels in supernatants of Jurkat T cells

Scramble (scr) and STX11 shRNA treated Jurkat T cells were stimulated with plate-coated anti-CD3 (10 μg/ml) and the culture supernatants were collected at 12 and 24 hour post-stimulation. EIA plates were coated with purified mouse anti-human IL-2 capture antibody (1 μg/ml) overnight at 4°C, washed with 0.05% PBST and blocked with 2% BSA for 2 hours. After blocking, the plates were incubated with the culture supernatants for 2 hours at RT, washed and incubated with the detection antibody (Biotin mouse anti-human IL-2, 1 μg/ml) for 2 hours. After washing, Streptavidin HRP (0.1 μg/ml) followed by TMB reagent were added. 250 mM HCl was added to stop the reaction and absorbance was measured at 450 nm.

### Isolation and culture of human PBMCs

Whole blood freshly collected in Heparin or Citrate Phosphate Dextrose Adenine (CPDA) solution was subjected to density gradient centrifugation using Ficoll-Paque PLUS media. The buffy coat was separated, washed twice with HBSS and cultured with IL-2 (50 ng/ml). Cells were stimulated once per week with PHA (2 μg/ml) for 72 hours and rested in IL-2 for the remaining 72-96 hours. Unless specified, all the assays were performed following 24-48 hours of rest, post stimulation.

### Isolation of genomic DNA and sequencing

Genomic DNA was extracted from PBMCs using Phenol-Chloroform-Isoamyl alcohol. STX11 genomic region flanking the mutation was PCR amplified using primers: 5’ Forward -CATGCACGACTACAACCAGGC and 3’ Reverse - GGGACAGCAGAAGCAGCAGAGGG. The resulting PCR products were separated on 2% agarose gel, excised and extracted using the Macherey-Nagel Nucleospin columns and subjected to Sanger sequencing using the 5’ Forward PCR primer to confirm the mutation.

### Measurement of SOCE in human PBMCs

PBMCs in culture were washed, rested and allowed to adhere to freshly carbon-coated glass bottom (IBDI) dishes for 1 hour in plain RPMI at 37°C. The cells were washed with HBSS and incubated with 1 ml of 1 μM Fura-2-AM diluted in Ringer’s buffer for 40 minutes, washed and incubated for an additional 10 minutes to allow de-esterification of the dye. The assay was started with 1 ml of Calcium free Ringer’s buffer in the imaging dish and images were captured every 4 seconds. After capturing the baseline for ∼60 seconds, stores were depleted by the addition of 1 μM Thapsigargin. In other assays, 10 μg/ml of anti-CD3 and 5 μg/ml of the secondary antibody were used to cross-link the TCRs and thereby induce store depletion. ∼5 minutes post-store-depletion, Calcium Chloride (CaCl_2_) was added back to the dish at a final concentration of 2 mM to estimate the magnitude of store operated calcium entry.

### Degranulation assay

PBMCs were cultured in RPMI containing 10% FBS and 50 ng/ml IL-2 and stimulated with PHA (2 μg /ml) 48hrs before the assay. 24hrs prior to the assay, IL-2 was washed off but PHA was re-added. On the day of the assay cells were washed twice to remove PHA and any growth factors. To measure degranulation, cells were either left unstimulated, stimulated with a combination of anti-CD3 (10 μg/ml plate-coated), anti-CD28 (2 μg/ml soluble) and anti-CD49d (2 μg/ml soluble) or a combination of Ionomycin (1 μM) and PMA (50 ng/ml) for 3.5 hours. CD107a-PE antibody (1:50 dilution) was added to each of the three groups at the start of stimulation. Following stimulations, cells were transferred to ice, washed with cold HBSS containing 2% FBS and incubated with anti-CD8 APC for 20 minutes, washed twice with cold HBSS and analyzed using Cytoflex (Beckman Coulter) flow cytometer.

### qPCR to estimate gene expression in human PBMCs

PBMCs were taken off IL-2 48 hours prior to the assay. On the day of the assay, cells were washed twice, left unstimulated and either stimulated with anti-CD3 (10 μg/ml, plate-coated), anti-CD28 (2 μg/ml) and anti-CD49d (2 μg/ml) or with Ionomycin (1 μM) and PMA (50 ng/ml) for 6 hours. To end the stimulation, cells were washed with cold HBSS, pelleted and used for RNA extraction. Total RNA was isolated using Qiagen RNeasy mini kit and cDNA was synthesized using random hexamers and Superscript IV (Invitrogen). cDNA was quantified using Qubit and subjected to QPCR analysis using Taqman probes for IL-2 and beta-actin housekeeping gene control in triplicates. Analysis was performed by calculating the double delta C_t_ values.

### Transduction of human PBMCs with pMSCV-STX11

For viral transduction of PBMCs, polystyrene non-TC treated 24-well plates were coated with retronection (20 μg/ml) overnight at 4°C, blocked with 2% BSA for 30 minutes at room temperature and washed twice with HBSS. The retroviral supernatants were added onto the coated wells and the plates were spun at 1800g for 2 hours at 30°C. Following spin, the wells were washed with the blocking solution. PBMCs cultured in RPMI containing 10% FBS were stimulated with PHA (2 μg/ml) and IL-2 (50 ng/ml) for 48 hours prior to transduction. Stimulated PBMCs were transferred to coated plates at a density of 0.25 million cells/well centrifuged at 400g for 40 minutes at 30°C. The cells were analyzed 48-72 hours post transduction.

### Patch clamp measurements

Patch clamp recordings were performed by using an Axopatch 200B amplifier (Axon Instruments, Foster City, CA) interfaced to an ITC-18 input/output board (Instrutech, Port Washington, NY) and an iMac G5 computer. Currents were filtered at 1 kHz with a four-pole Bessel filter and sampled at 5 kHz. Recording electrodes were pulled from 100-µl pipettes (VWR), coated with Sylgard, and fire-polished to a final resistance of 2 to 5 MΩ. Stimulation and data acquisition and analysis were performed by using in-house routines developed by RS Lewis (Stanford University) on the Igor Pro platform (Wavemetrics, Lake Oswego, OR). All data were corrected for the liquid junction potential of the pipette solution relative to Ringer’s in the bath (−10 mV) and for leak currents collected in 20 mM extracellular free calcium plus 10µM La^3+^. The holding potential was +30 mV. The standard voltage stimulus consisted of a 100-ms step to –100 mV followed by a 100-ms ramp from –100 to +100 mV applied at 1 s intervals. I_CRAC_ was typically activated by passive depletion of ER Ca^2+^ stores by intracellular dialysis of 8 mM BAPTA.

### Solutions and chemicals for patch clamp

The standard extracellular Ringer solution contained 130 mM NaCl, 4.5 mM KCl, 20 mM CaCl_2_, 10 mM tetraethylammoniumchloride (TEA-Cl), 10 mM D -glucose and 5 mM HEPES (pH 7.4 with NaOH). The standard divalent-free (DVF) Ringer solutions contained (in mM) 150 NaCl, 10 HEDTA, 1 EDTA, and 10 HEPES (pH 7.4). The internal solution contained: 135 mM Cs aspartate, 8 mM MgCl_2_, 8 mM Cs-BAPTA and 10 mM HEPES (pH 7.2 with CsOH).

### Patch clamp data analysis

Analysis of current amplitudes was typically performed by measuring the peak currents during the −100 mV pulse. Averaged results are presented as the mean value standard error of the mean (SEM). Reversal potentials were measured from the average of several leak-subtracted sweeps (4–6) in each cell. For datasets with two groups, statistical analysis was performed with two-tailed t test to compare between control and test conditions with a confidence level of 95%, and results with p<0.05 were considered statistically significant.

### STX11:Orai1 and SOAR:Orai complex prediction

Full length STX11 and its two domains, (H_abc_ 41-167 and SNARE 183-267) and Orai1 (N-terminus 1-87 and C-terminus 256-301) were used to generate the complex of STX11 and Orai1 using AlphaFold3 (AF3) in all combinations. For Stim1, SOAR, residue 344-444 were used along with Orai C-terminus 256-301. The resultant models were assessed using ipTM and pTM scores. The best scoring combinations were further considered to generate more models by changing the seed values. The best model in terms of highest ipTM and pTM values was considered for further analysis. Custom script was used to analyze the contact frequency of interacting residues of STX11-H_abc_ and Orai1 C-terminus complex as well as SOAR and Orai1 C-terminus, across all predicted models.

### Molecular dynamics (MD) simulation

The STX11 H_abc_:Orai1 C-terminus and SOAR:Orai1 C-terminus complexes were subjected to all-atom MD simulations. Initially, the complexes were prepared using protein-preparation module of Schrodinger which involves H-bond network optimization and restrained minimization of the initial structure. The prepared structures were solvated using TIP3P water model in an orthorhombic water box and neutralized by adding counter ions. OPLS4 force field was used, and simulation system was generated by specifying 150 mM salt (NaCl). A buffer distance of 10 Å beyond the solute in each direction was used. The solvated system was subjected to default relaxation protocol of Desmond followed by production run for 500 ns at 300K and 1 atm pressure in NPT ensemble. The default relaxation protocol includes several short simulation steps. (1) Brownian dynamics simulation for 100 ps at 10 K temperature in NVT with restraints on solute heavy atoms (2) Simulation in NVT ensemble at 10 K for 12 ps with restraints on solute heavy atoms (3) 12 ps simulation in NPT ensemble at 10 K with restraints on solute heavy atoms (4) Simulation in NPT ensemble for 12 ps with restraints on solute heavy atoms (5) Simulation in NPT ensemble for 24 ps without restraints. Three independent runs were executed with different initial seeds for both structures.

### Analysis of MD simulation

The MD runs were analyzed for the stability of the complex and interactions between the subunits. Simulation stability was assessed based on the protein backbone RMSD over simulation time. The RMSD was calculated using simulation interaction diagram (SID) module of Schrodinger. The first frame was used as reference frame for RMSD calculation. Script “analyze_trajectory_ppi.py” was used to calculate the interactions between STX11:Orai1 and Stim1:Orai1. The binding energy (ΔG) at each nanosecond across simulation was calculated using “thermal_mmgbsa.py” script of Schrodinger. The trajectories of the three runs of STX11:Orai1 and Stim1:Orai1 interactions have been deposited in Zenodo (https://doi.org/10.5281/zenodo.21663510). The representative structure was generated through trajectory clustering using the trajectory clustering tool in Schrödinger, which applies affinity propagation to the pairwise backbone RMSD-based similarity matrix. Within each affinity propagation run, convergence was defined as no change in the set of exemplar frames for 15 consecutive iterations, with a maximum of 400 iterations per run. If convergence was not reached, the damping factor was increased from 0.5 in increments of 0.01 until convergence. The last 200 ns frames from each run were combined and clustered using scripts “trj_merge.py” and “trajectory cluster” of Schrodinger. The cluster representative of the largest cluster was used to analyze the protein-protein interactions. PDB Sum was used to calculate the interactions in the cluster representative. Figures were generated using Pymol and MD movies using Maestro.

### Statistical Analysis

Statistical analysis was done using unpaired two-tailed Student’s *t* test where only two groups were being compared and ANOVA where more than two groups were present. Normality of distribution was checked in Origin. Mann Whitney U Test was used for datasets with non-normal (or skewed) distribution. P values for all significance tests were *P<0.05; **P<0.01; ***P<0.001.

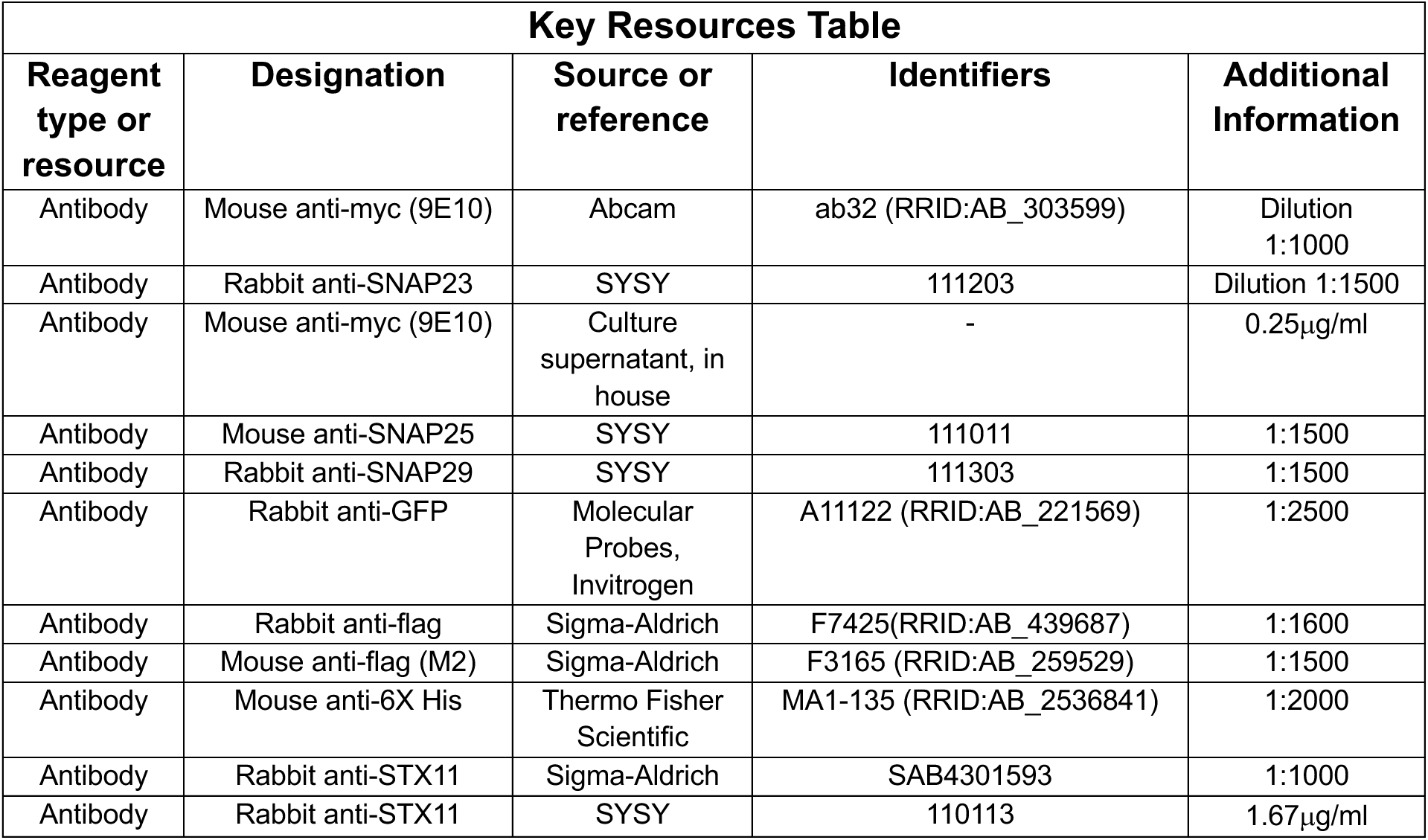
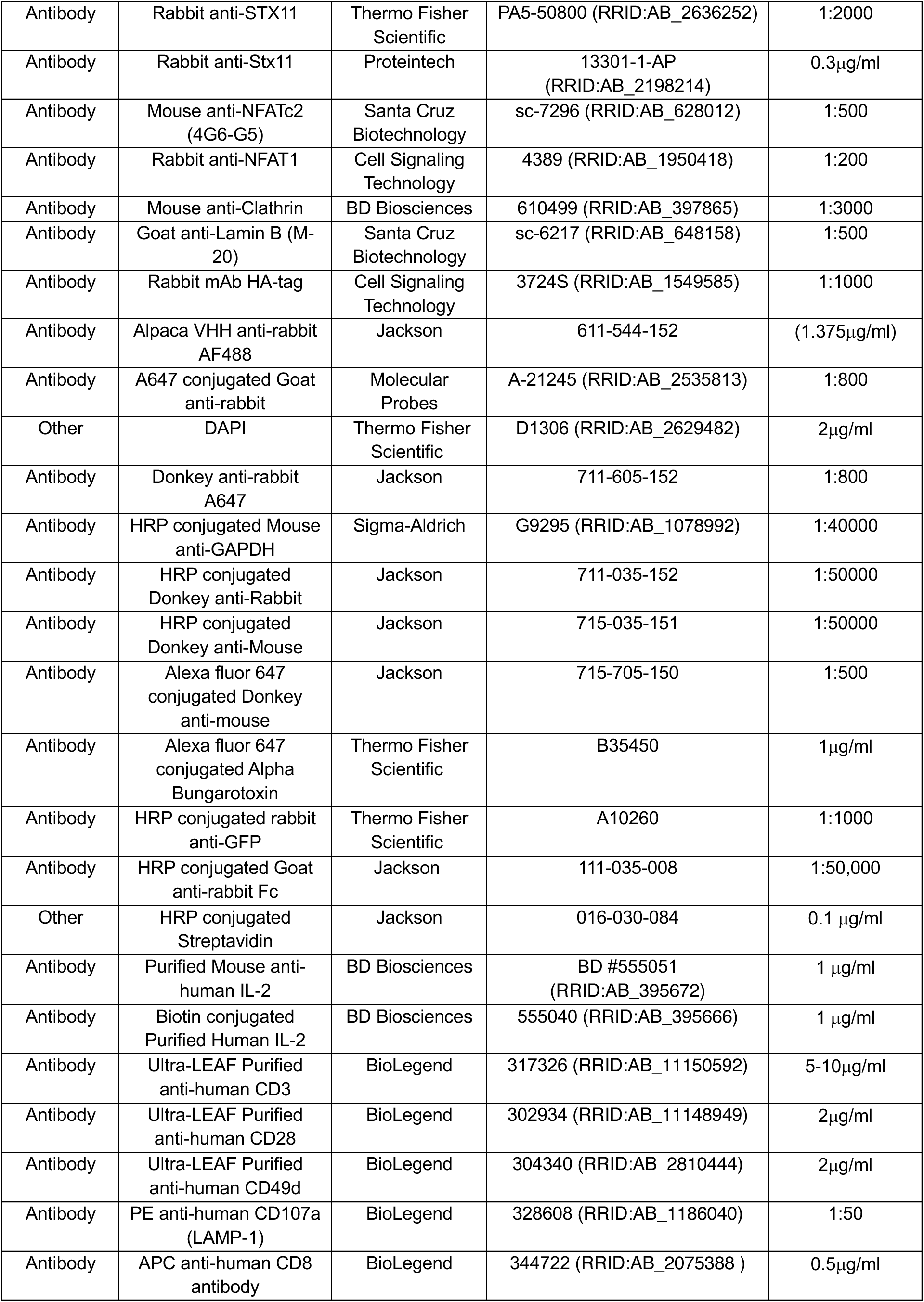
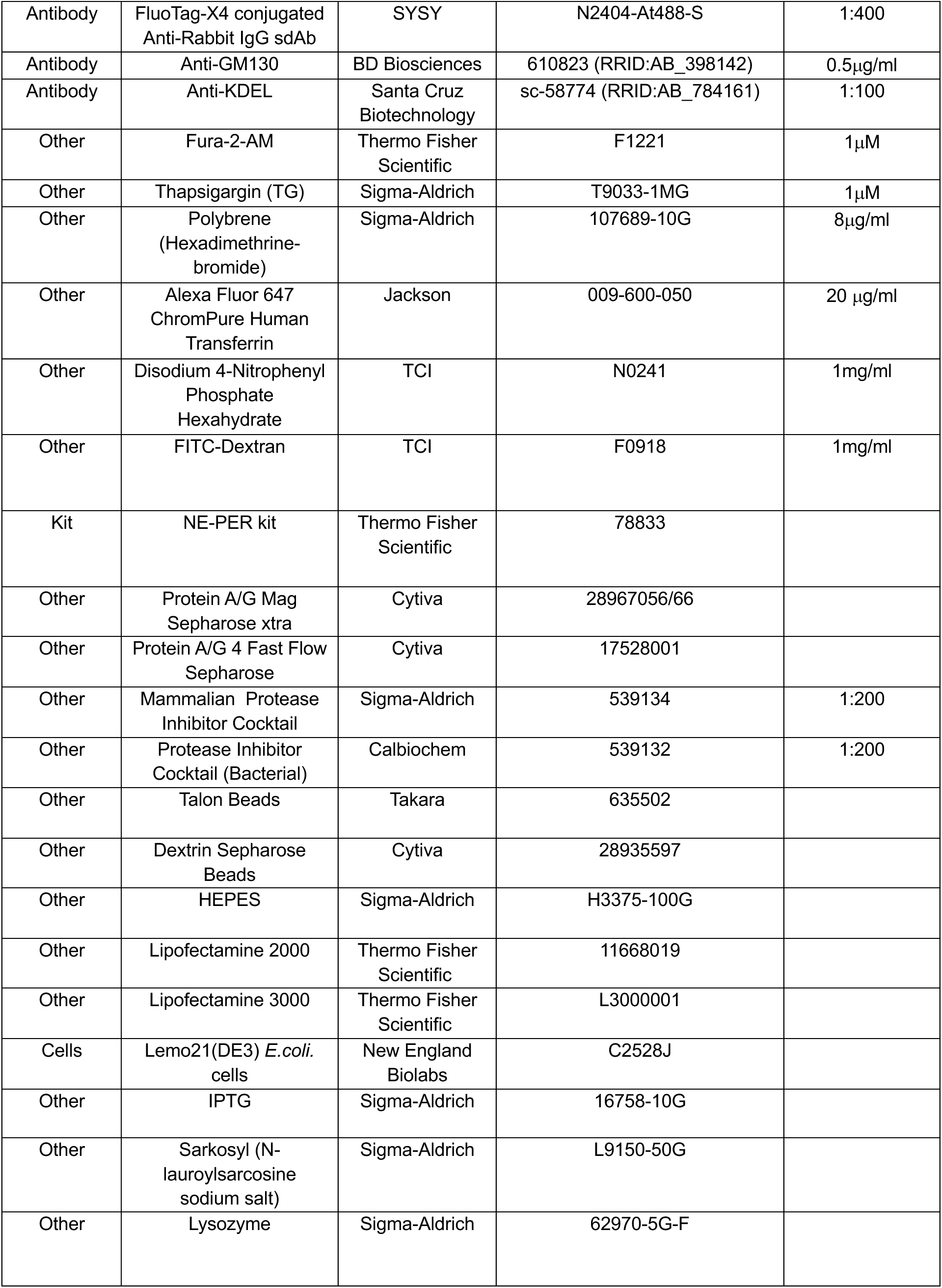
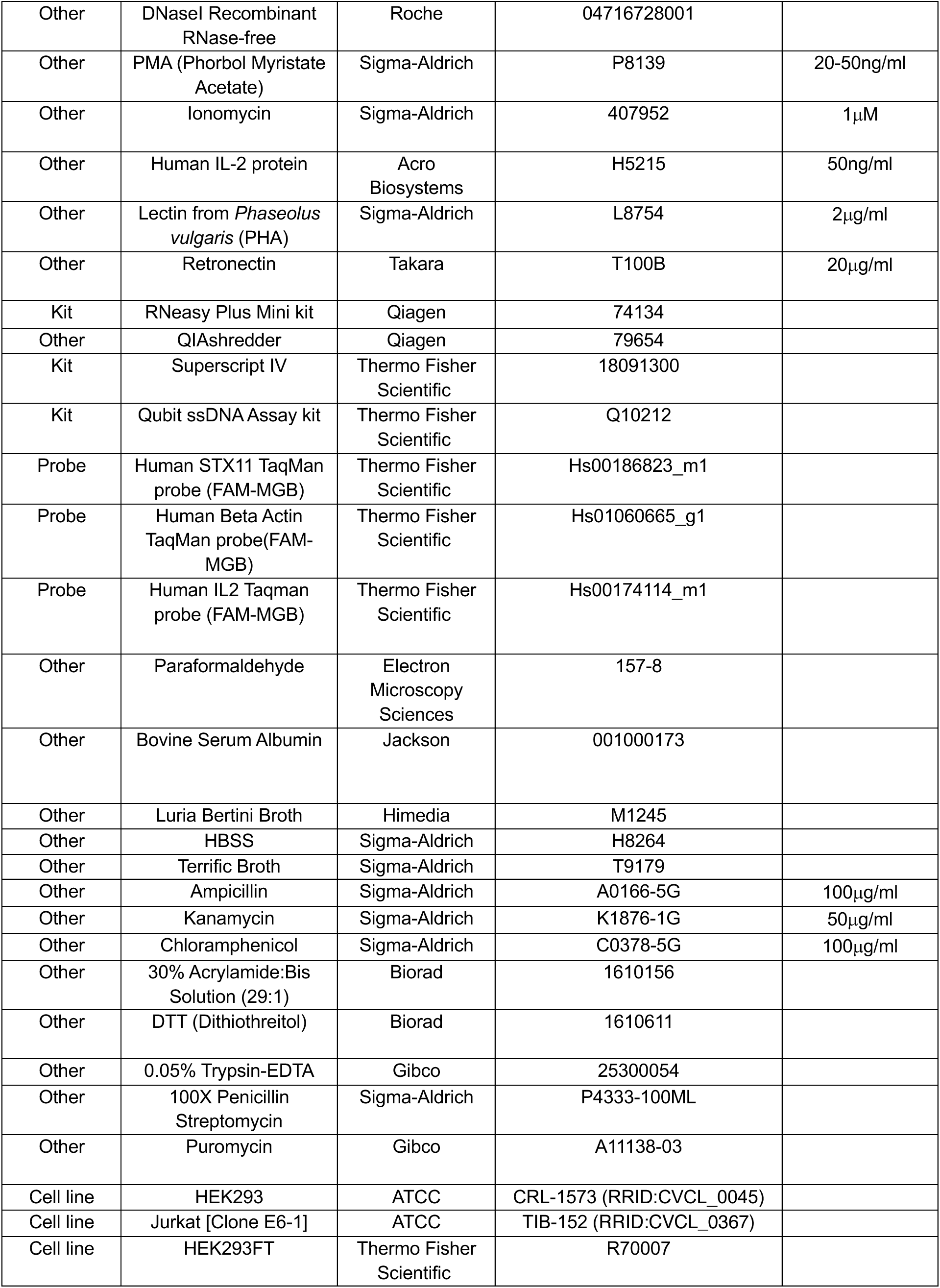
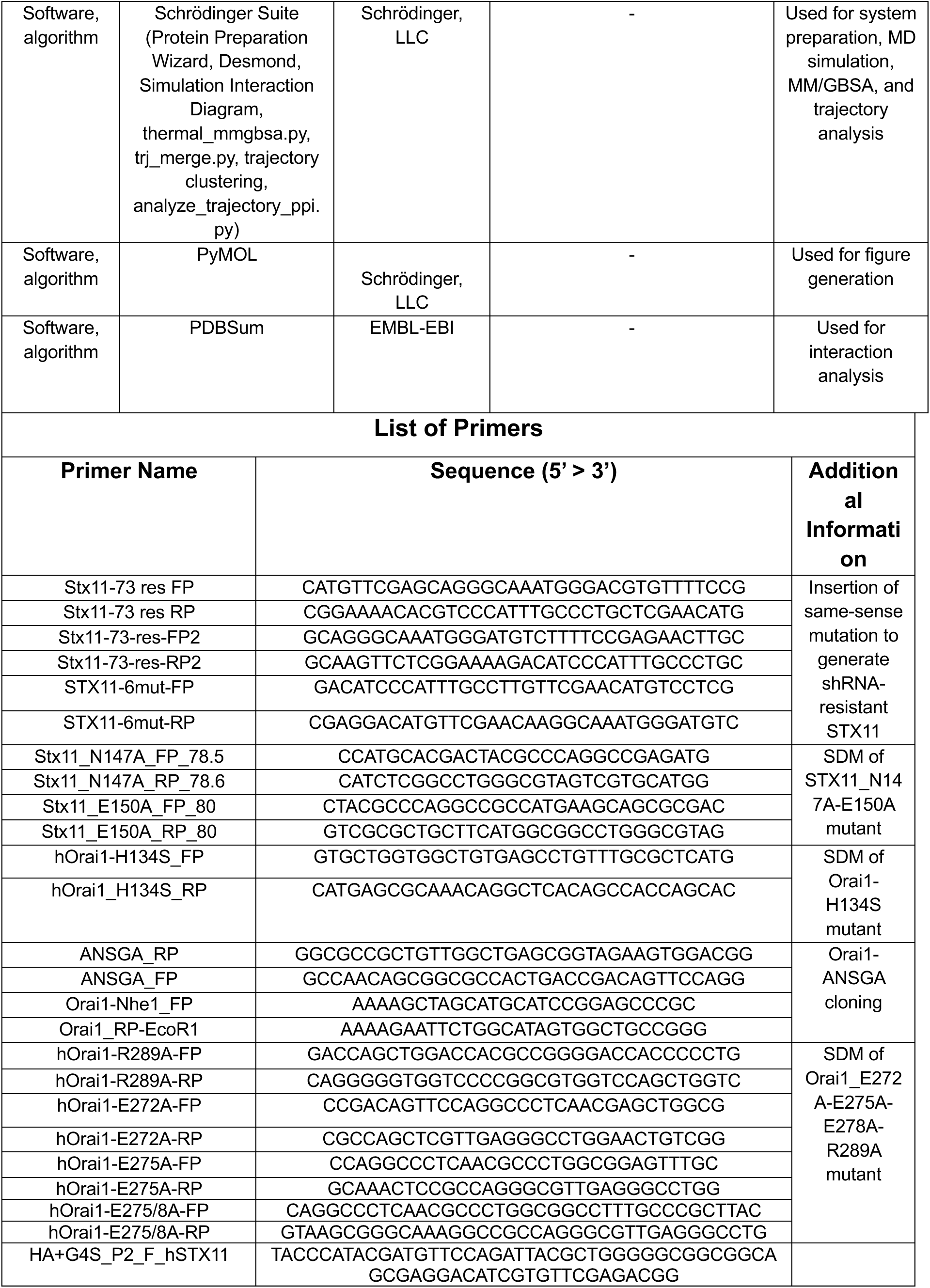
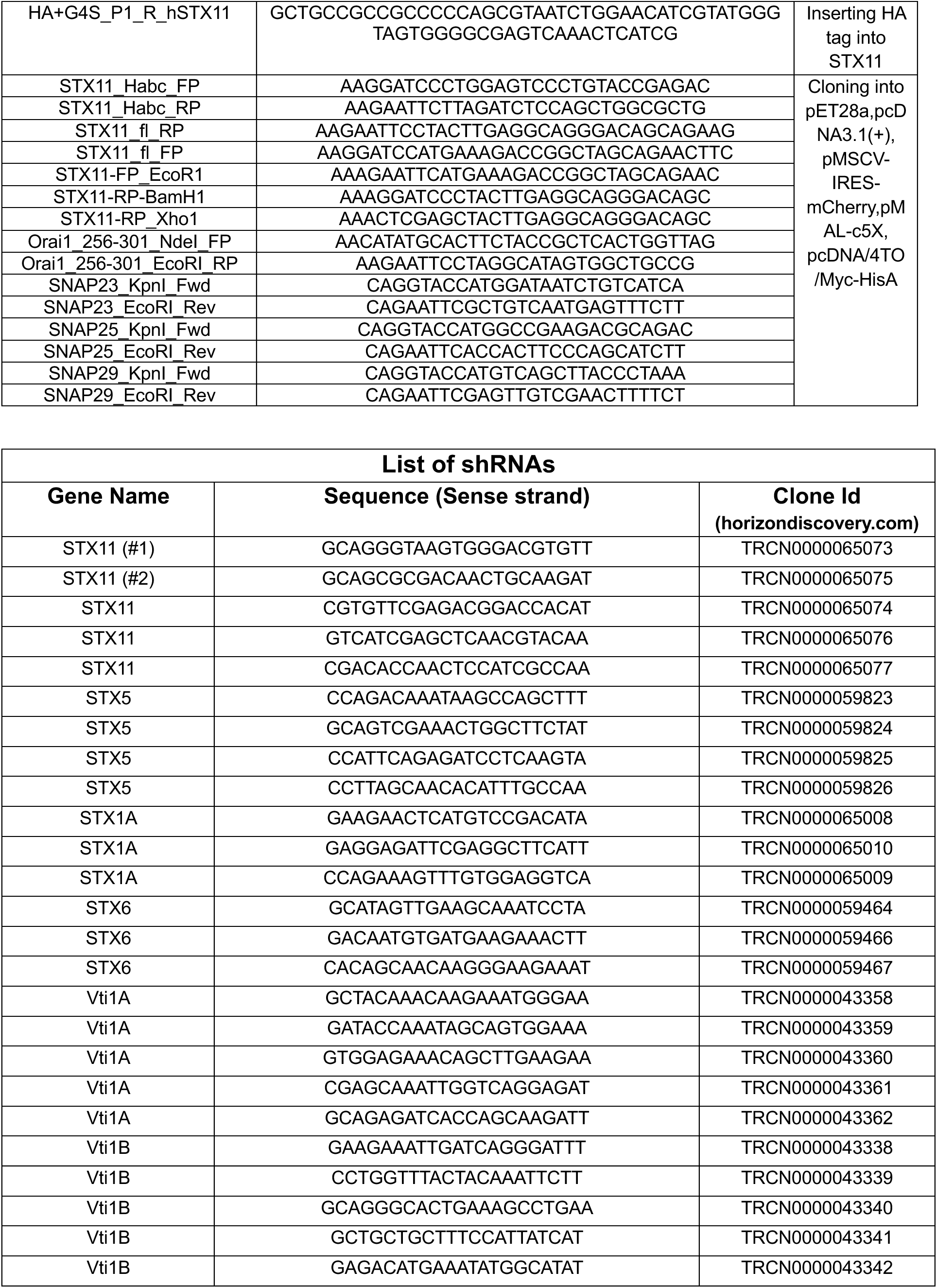
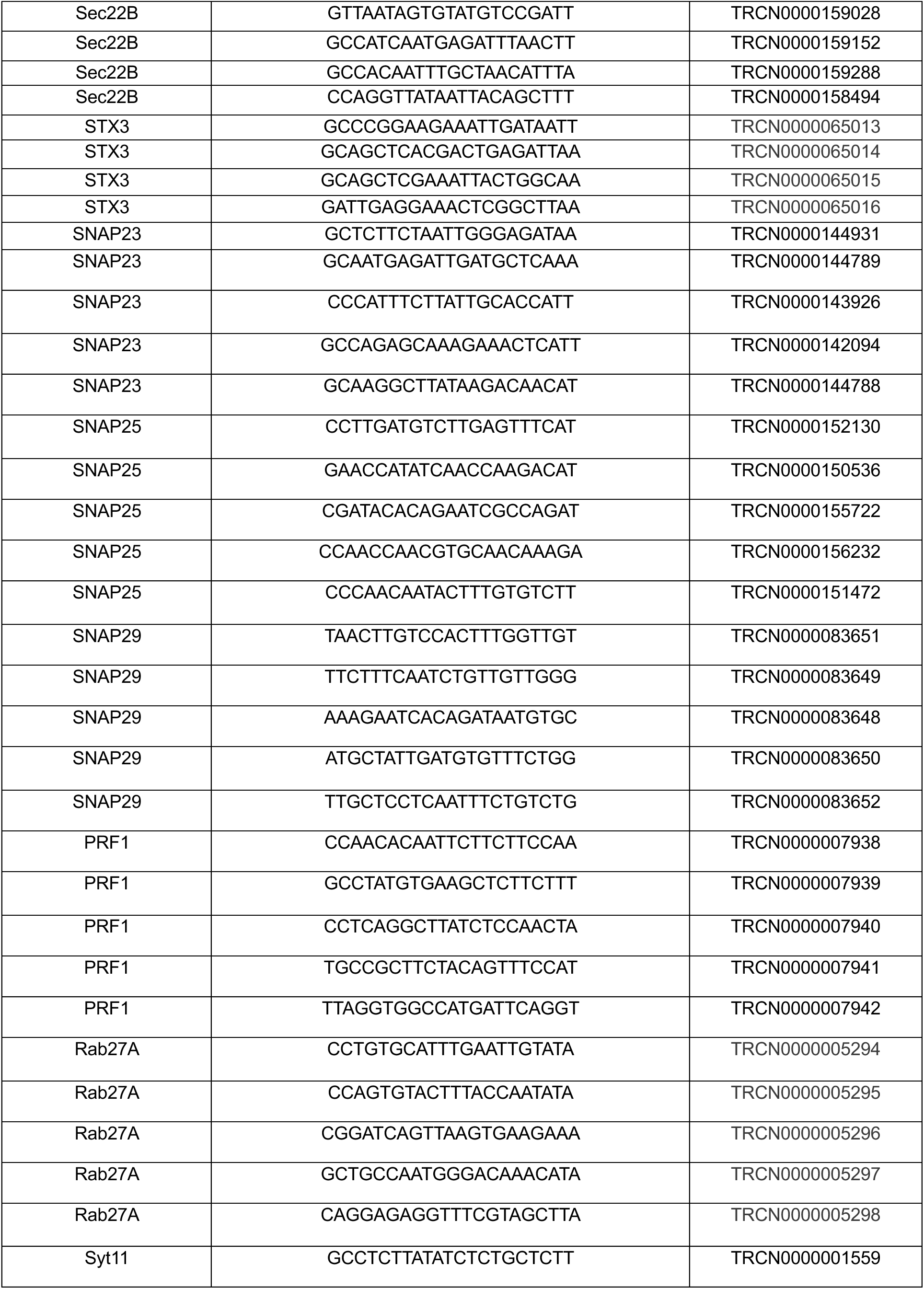
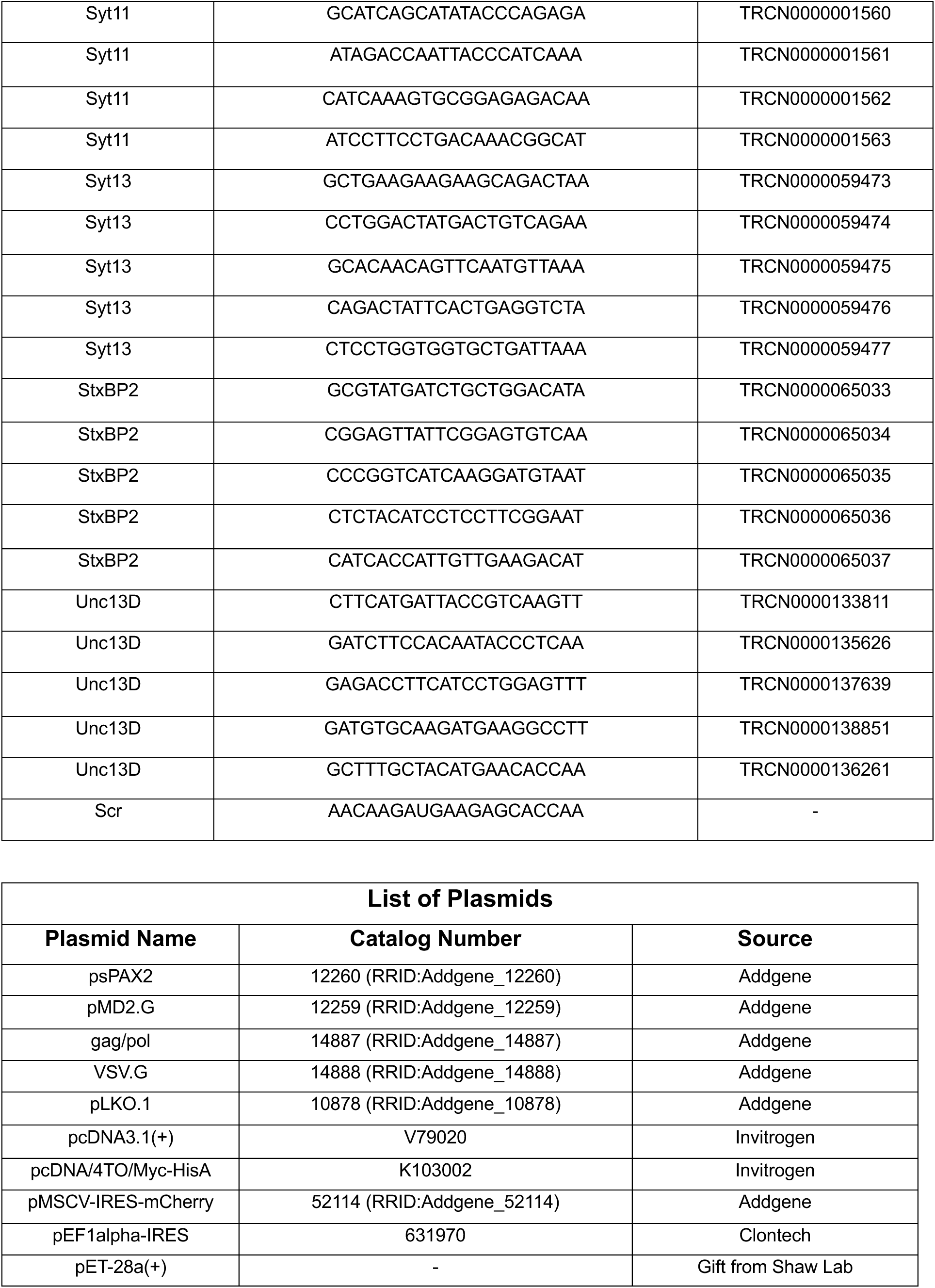
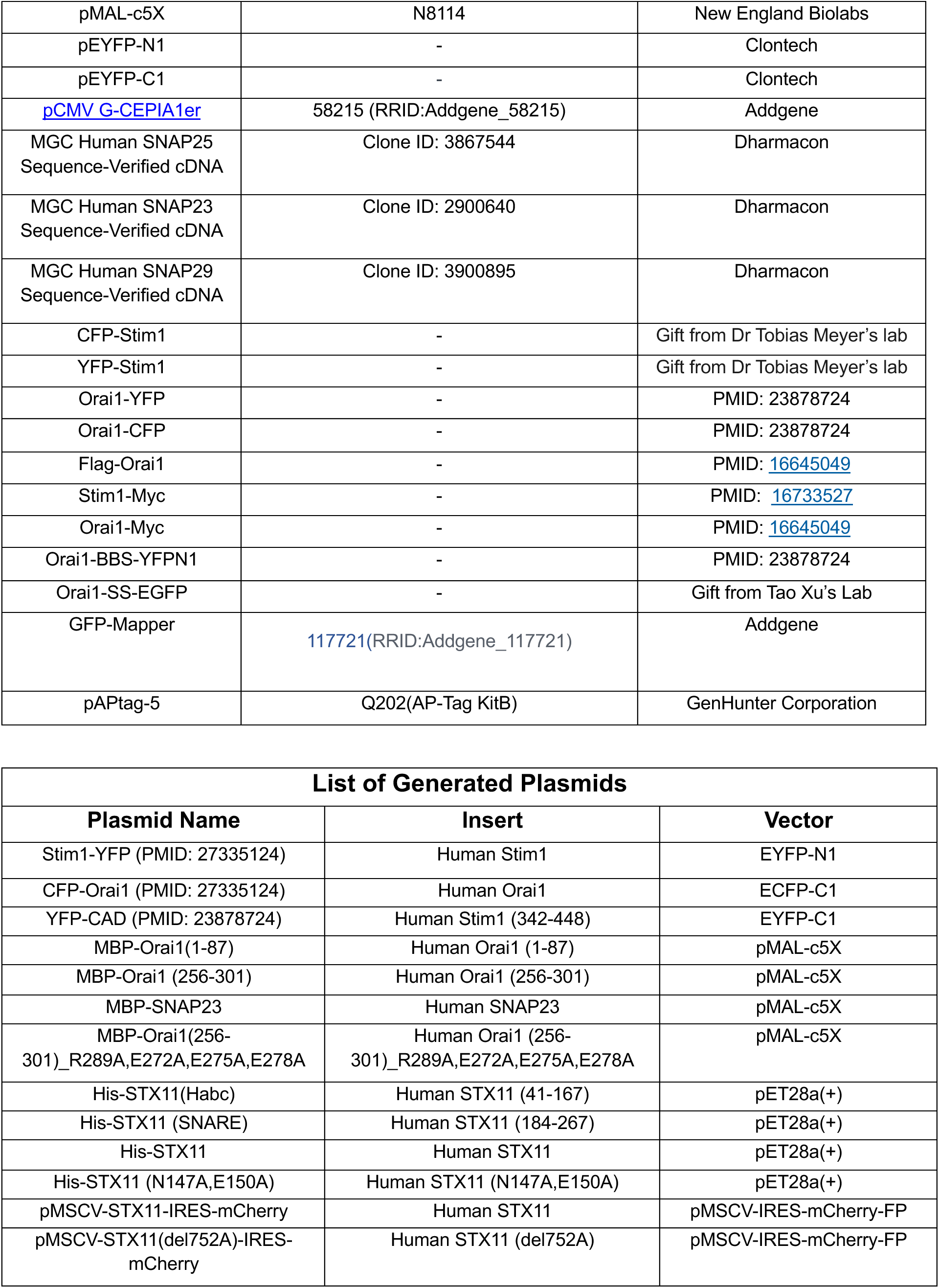
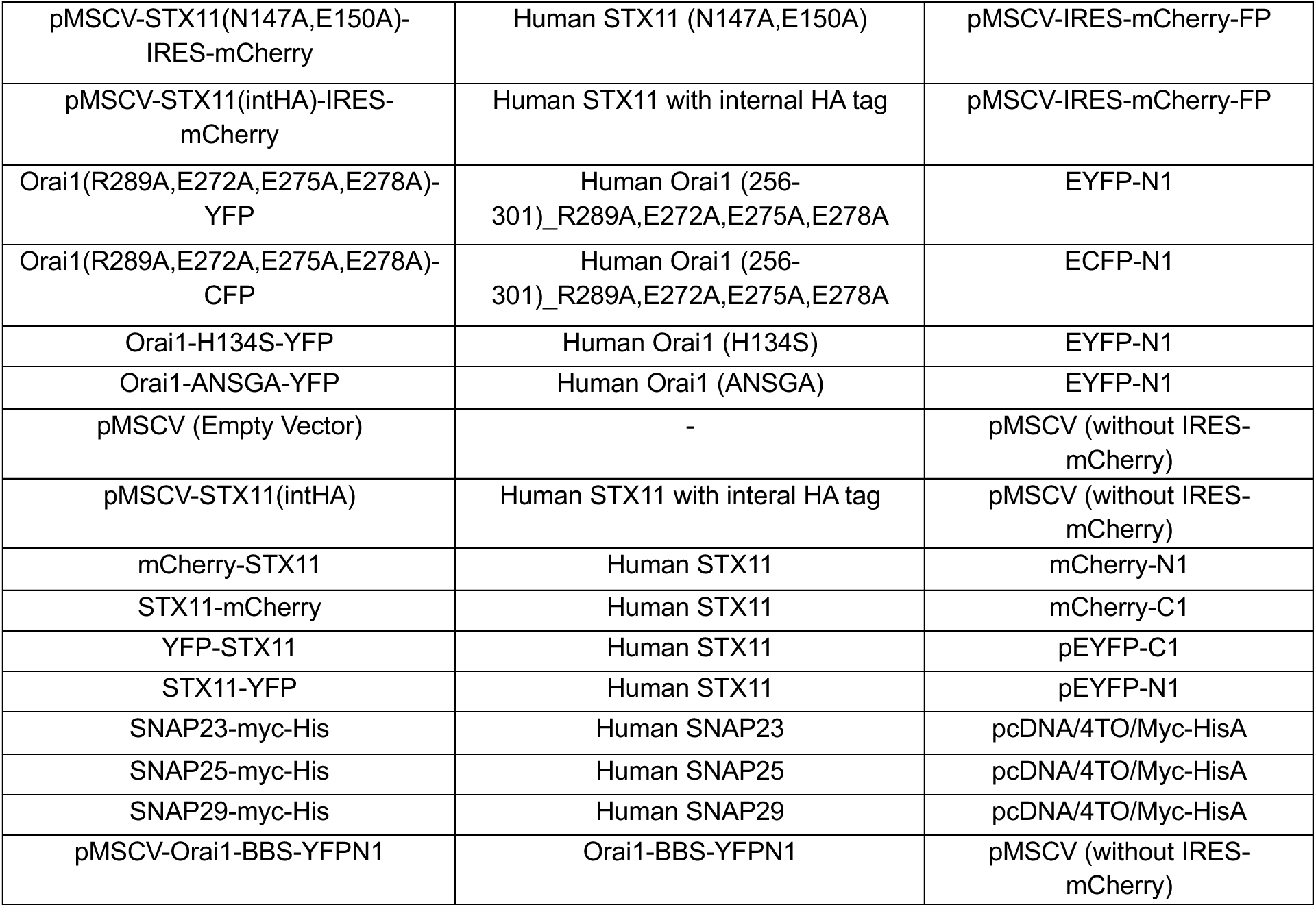

## Acknowledgements

We thank Vivien Beziat, Yenan Bryceson, Heinrich Schlums, Jelve Nejati Zendegani, Stephan Ehl, and Jasmin Mann for help with human PBMC handling, isolation, storage and shipping. Anand Vaidya for advice on STX11 purification. Jyoti Rohilla for technical support. Bio Render for the generation of models and IBS 2.0 for the generation of schematics shown in this manuscript. This work was supported in part by Department of Atomic Energy, Government of India under Project Identification No. RTI 4007, ICMR Grant No. EMDR/SG/11/2025-01-02427 and NIH-NIAID grant AI108636.

---

**Movies 1-3: Molecular dynamic simulations of Orai1 C-terminus interaction with the STX11_H_ABC_ domain.** https://doi.org/10.5281/zenodo.21663510

**Movies 4-6: Molecular dynamic simulations of Orai1 C-terminus interaction with Stim1 SOAR domain.** https://doi.org/10.5281/zenodo.21663510

**Supplementary Figure 1.**
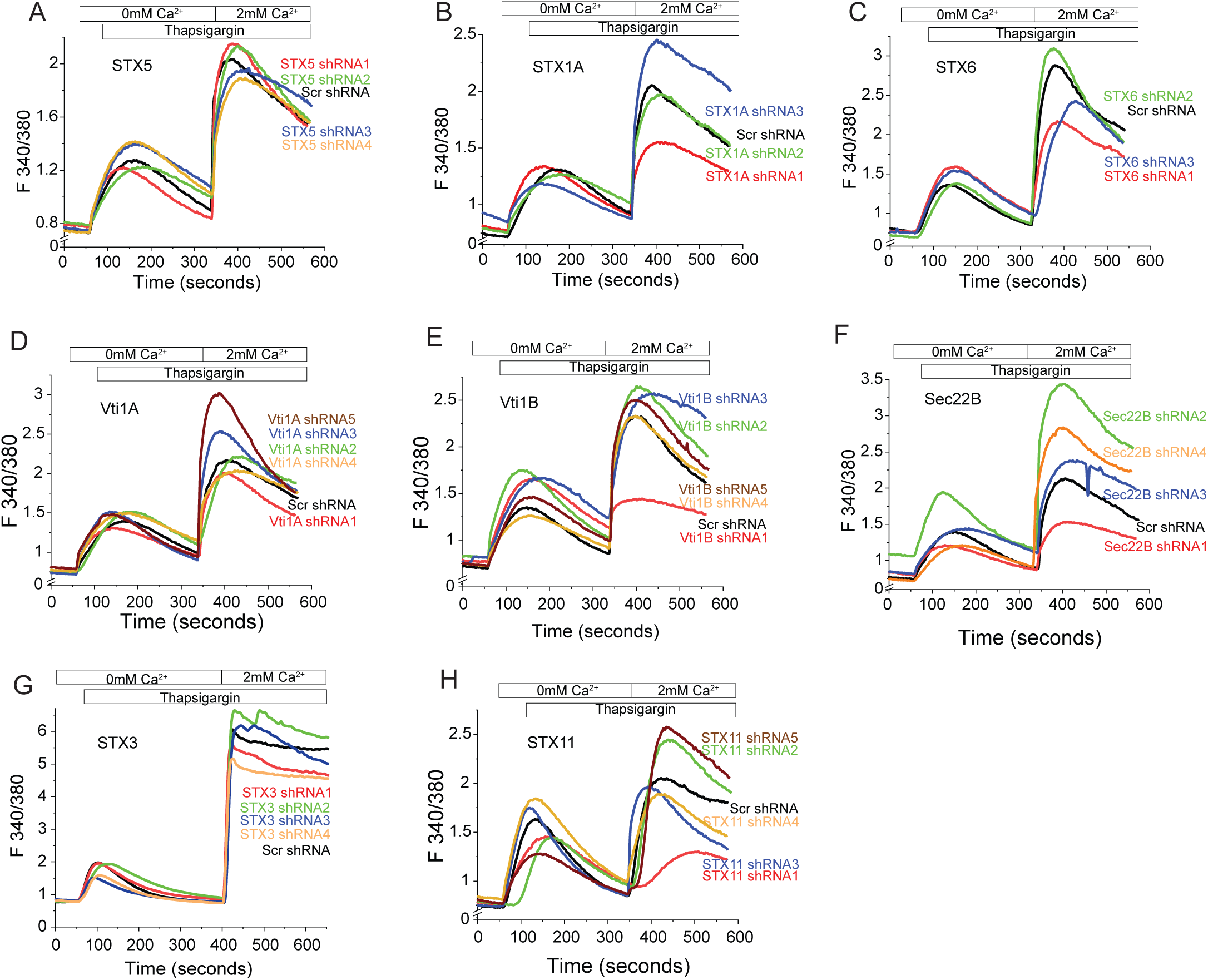
**(A-H)** Targeted RNAi screen for SNAREs involved in SOCE. Fura-2 calcium imaging assays measuring thapsigargin (TG) induced SOCE in HEK293 and/or Jurkat cell lines treated with scramble (scr) shRNA or different shRNA sequences against the gene of interest for 3-4 days. Shown here are the average traces from 30-40 cells per group.

**Supplementary figure 2.**
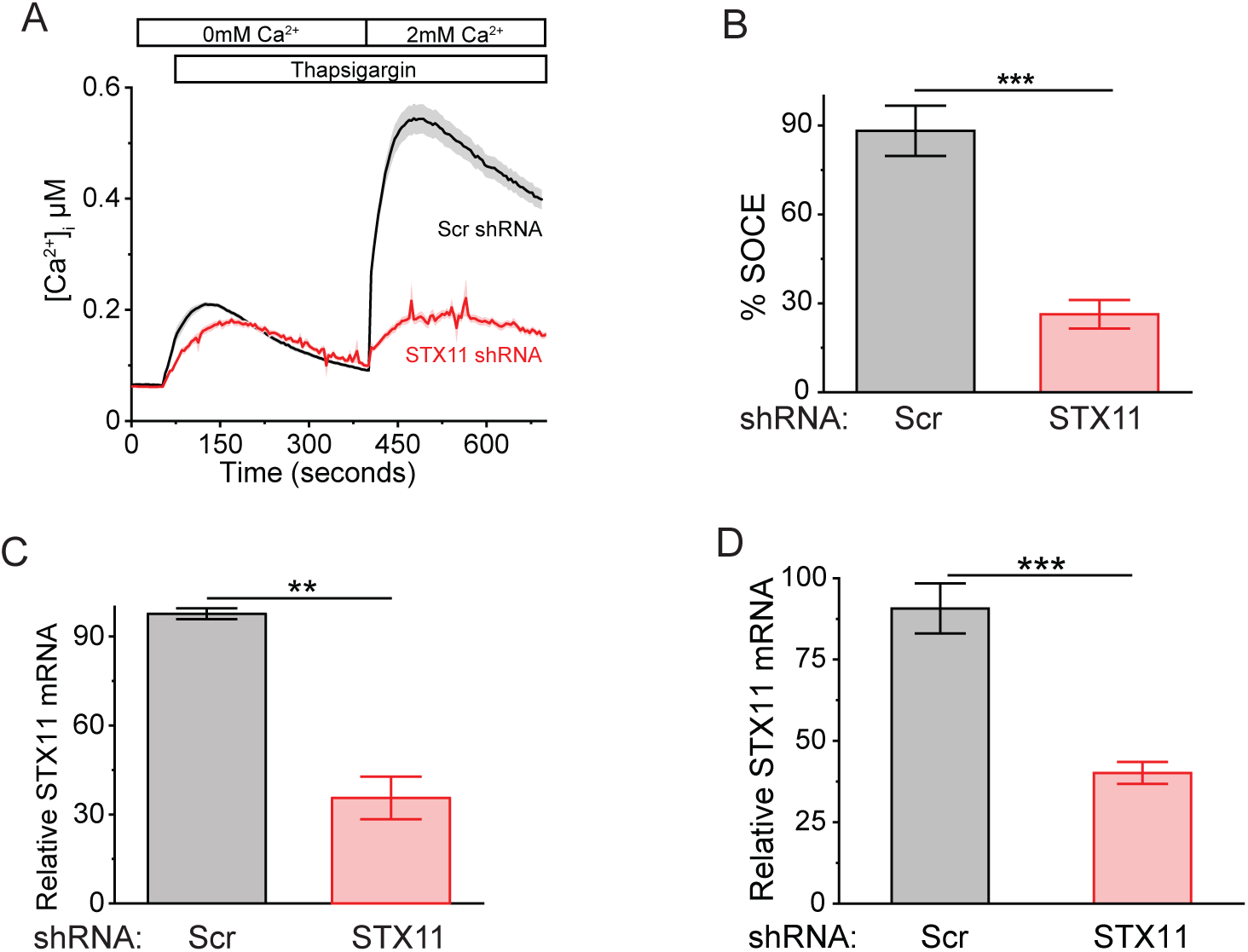
**(A-B)** Representative Fura-2 calcium imaging assay **(A)** and its quantification from repeats **(B)** measuring thapsigargin (TG) induced SOCE in HEK293 cells treated with scr or STX11 shRNA. Bars show relative SOCE ± SE from three independent experiments in **(A)** where SOCE from scramble shRNA treated group in one experiment was set at 100%. **(C)** Quantitative PCR to assess the efficiency of knockdown in shRNA treated HEK293 **(C)** and Jurkat T cells **(D)** from 3 independent repeats. Total RNA extracted from cells treated with STX11 shRNA was subjected to qPCR analysis using Taqman probes for STX11. Data were normalized to Beta-actin housekeeping control.

**Figure 1-figure supplement 3:**
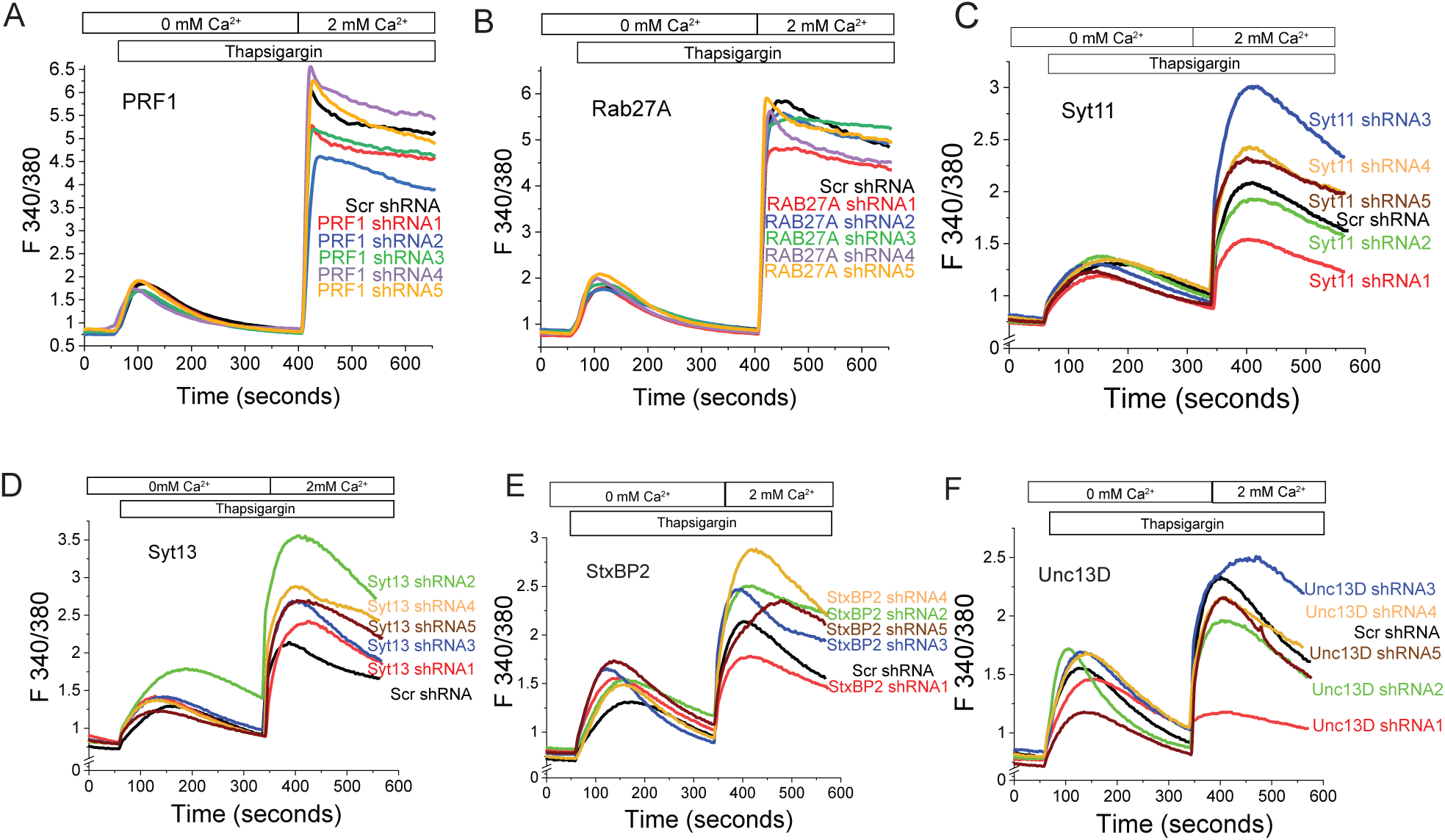
RNAi screen for genes involved in primary HLH and/or vesicle fusion for their role in SOCE. **(A-F)** Fura-2 calcium imaging assays measuring thapsigargin (TG) induced SOCE in HEK293 or Jurkat cells treated with scramble (scr) shRNA or different shRNA sequences against the gene of interest for 3-4 days. Shown here are the average traces from 30-40 cells per group.

**Figure 2-figure supplement 1:**
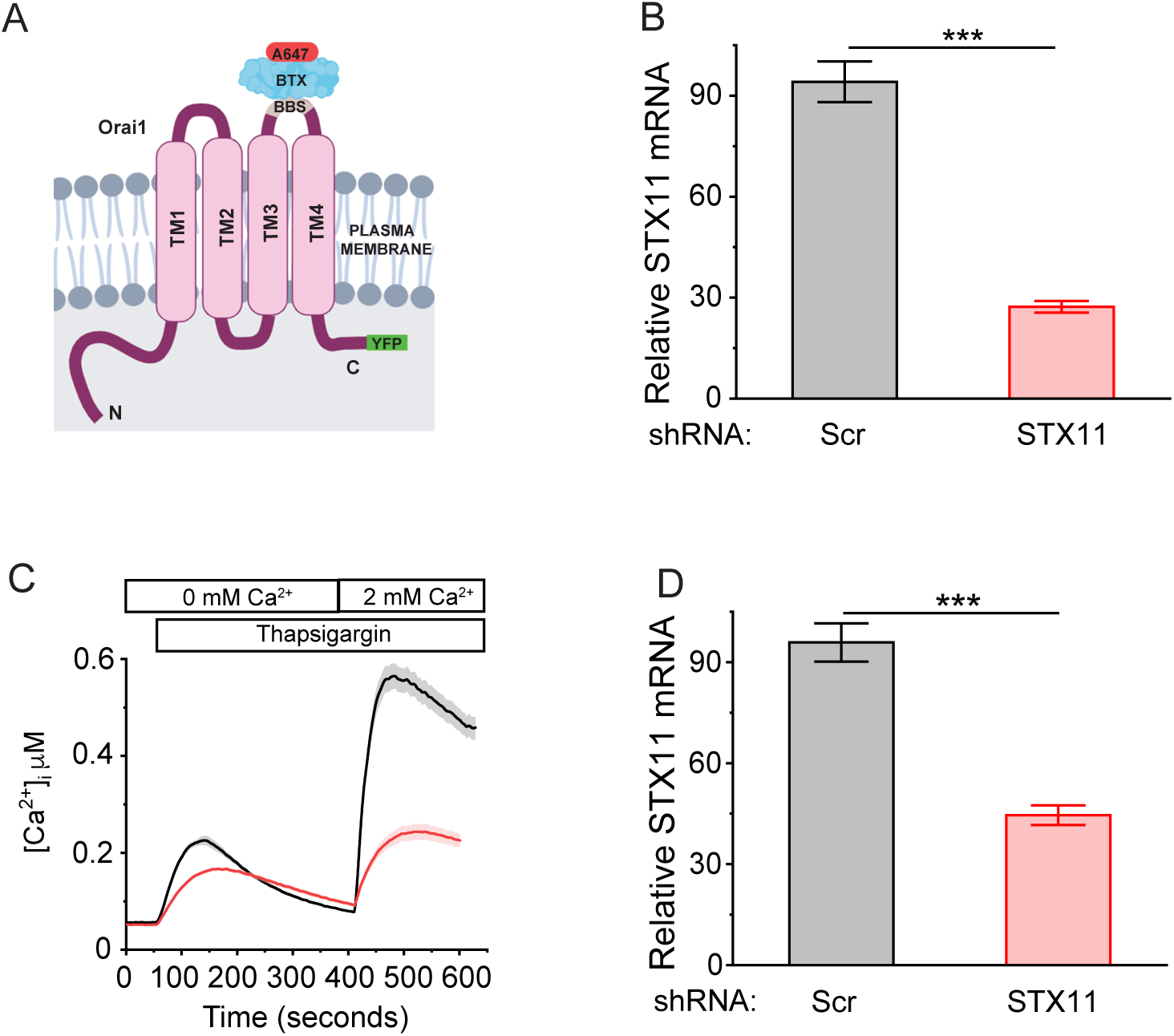
**(A)** Schematic showing the design of recombinant Orai1-BBS-YFP fusion protein. Shown here is the placement of the bungarotoxin binding site (BBS) inside the second extracellular loop and YFP tag in the C-terminus of Orai1 as well as binding of BTX-A647 to BBS. **(B)** Representative quantitative PCR to assess the efficiency of knockdown in shRNA treated HEK293 cells stably expressing Orai1-BBS-YFP. Total RNA extracted from cells treated with STX11 shRNA was subjected to qPCR analysis using Taqman probes for STX11. Data were normalized to RPL30 house-keeping control. **(C)** Representative Fura-2 calcium imaging assay measuring thapsigargin (TG) induced SOCE in Orai1-BBS-YFP expressing HEK293 cells treated with scr or STX11 shRNA. **(D)** Representative quantitative PCR to assess the efficiency of knockdown in shRNA treated Jurkat T cells expressing Orai1-BBS-YFP. Total RNA extracted from cells treated with STX11 shRNA was subjected to qPCR analysis using Taqman probes for STX11. Data were normalized to RPL30 housekeeping control.

**Figure 2-figure Supplement 2.**
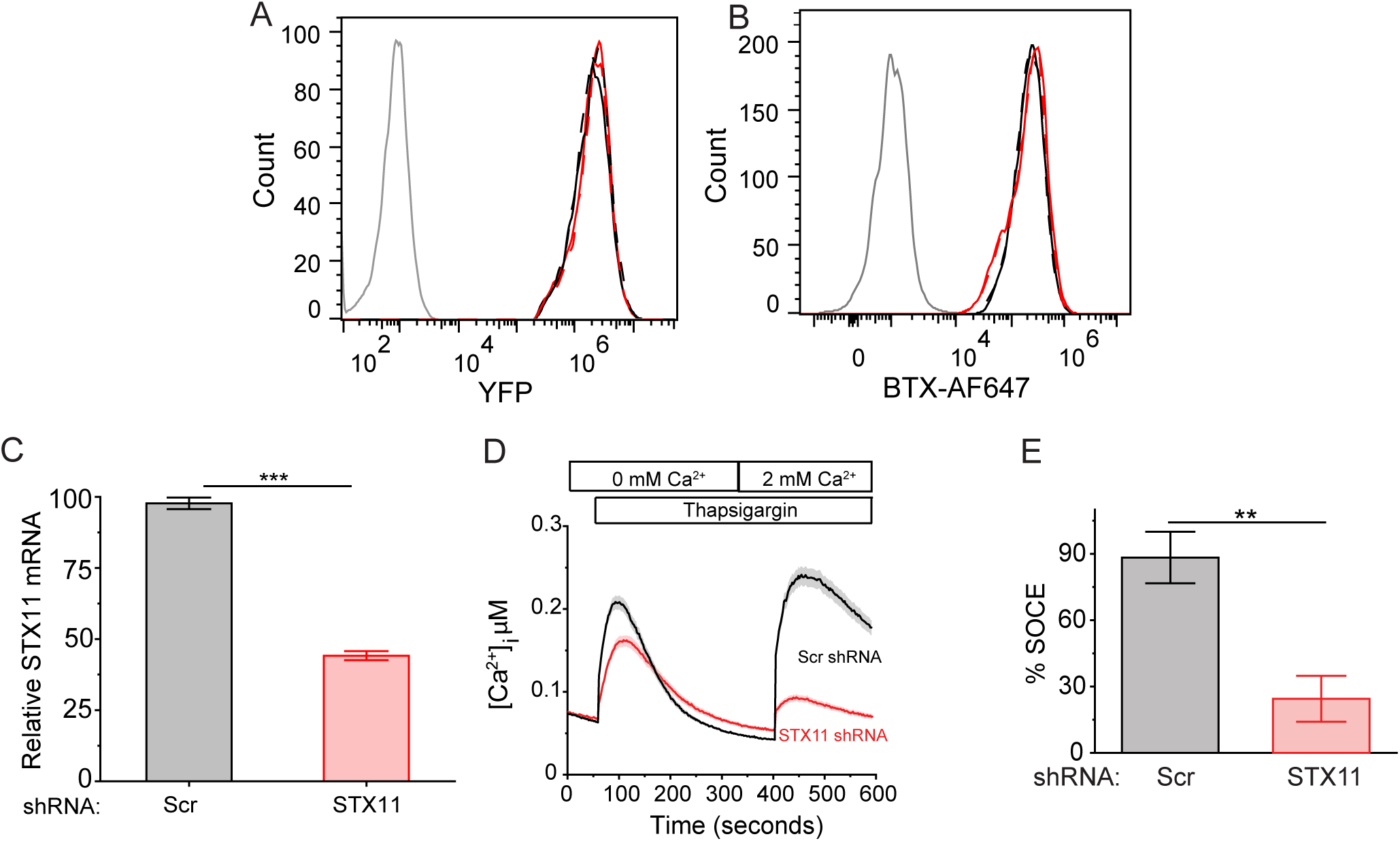
**(A-B)** Measurement of total Orai1 levels in the plasma membrane of STX11 depleted cells by flow-cytometry. U2OS cells stably expressing Orai1-BBS-YFP were transduced with Scr (black) or STX11 (red) shRNA, stimulated with 1uM TG (broken lines) or control (solid lines) and incubated with alpha-bungarotoxin alexa fluor 647 (BTX-AF647). Total Orai1-YFP levels **(A)** and BTX-AF647 binding to surface Orai1 **(B)** were measured where binding to wildtype U2OS cells was used as the background fluorescence. N=3 **(C)** Representative quantitative PCR to assess the efficiency of knockdown in shRNA treated Orai1-BBS-YFP expressing U2OS cells from **(A-B)**. Total RNA extracted from cells treated with STX11 shRNA was subjected to qPCR analysis using Taqman probes for STX11. Data were normalized to RPL30 housekeeping gene. **(D-E)** Measurement of thapsigargin (TG) induced SOCE in U2OS cells treated with STX11 RNAi in **(A-B)**. **(D)** Representative averaged traces of Fura-2 calcium imaging assay performed on cells in panels (A-B). **(E)** Bar plot showing relative mean % SOCE ± SE from three independent experiments in **(D)**.

**Figure 2-figure supplement 3.**
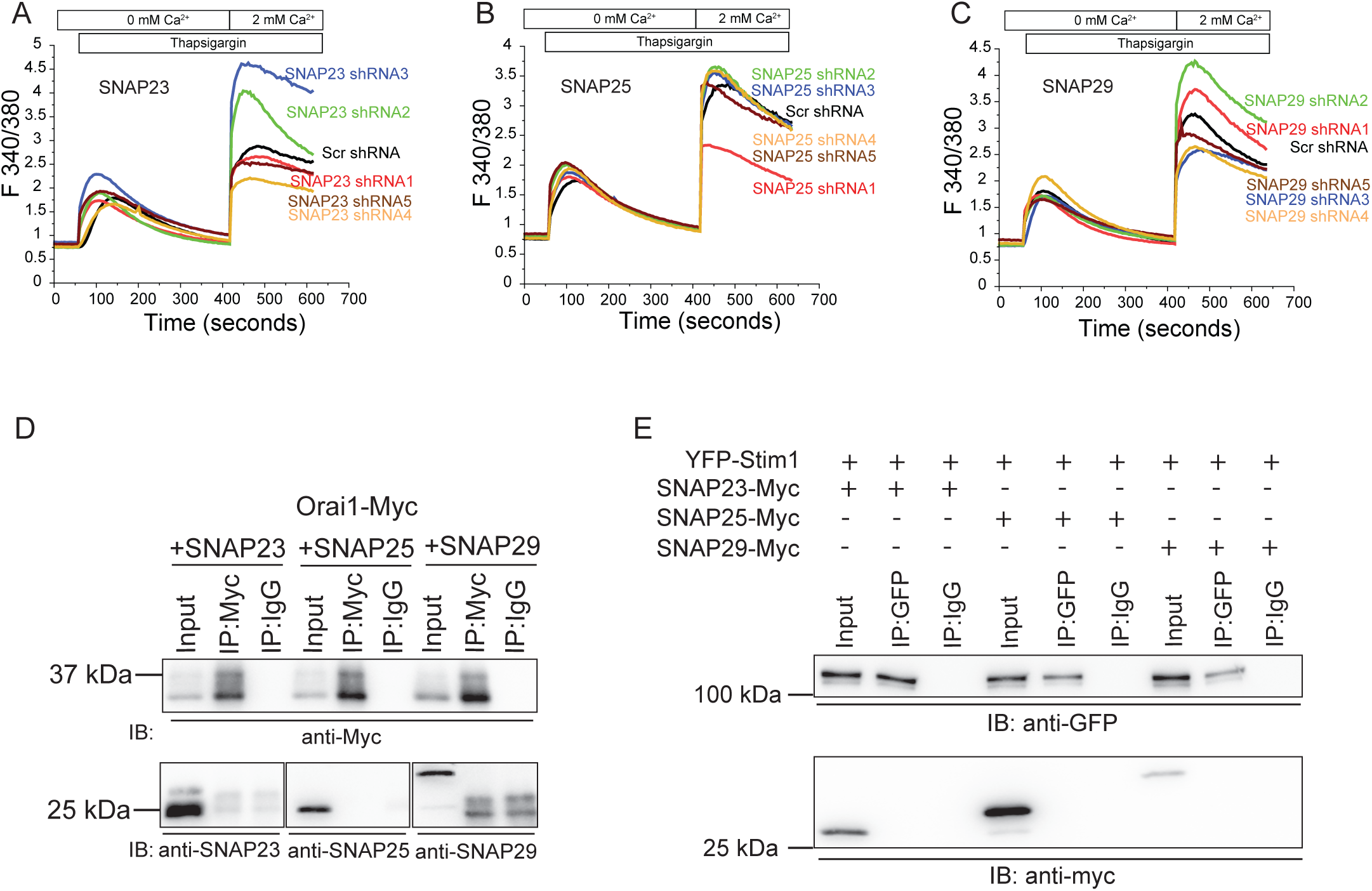
SOCE in RNAi treated Jurkat cells to assess the role of SNAPs. **(A-C)** Fura-2 calcium imaging assay measuring SOCE in cells treated with scramble (scr) shRNA or different shRNA sequences against SNAP23 **(A)** SNAP25 **(B)** SNAP29 **(C)**. Average traces from 30-40 cells per group are shown. **(D)** Western blot images of co-IPs to test the association of Orai1 with SNAP23/ SNAP25/ SNAP29. HEK293 cells were co-transfected with Orai1-Myc and untagged SNAP23/ SNAP25/ SNAP29, store-depleted, lysed and subjected to co-IP followed by western blot as indicated. (N=2) **(E)** Western blot images of co-IP to test the association of Stim1 with SNAP23/ SNAP25/ SNAP29. HEK293 cells were co-transfected with YFP-Stim1 and Myc-tagged SNAP23/ SNAP25/ SNAP29, store-depleted, lysed and subjected to co-IP followed by western blot as indicated. (N=2).

**Figure 5-figure supplement 1.**
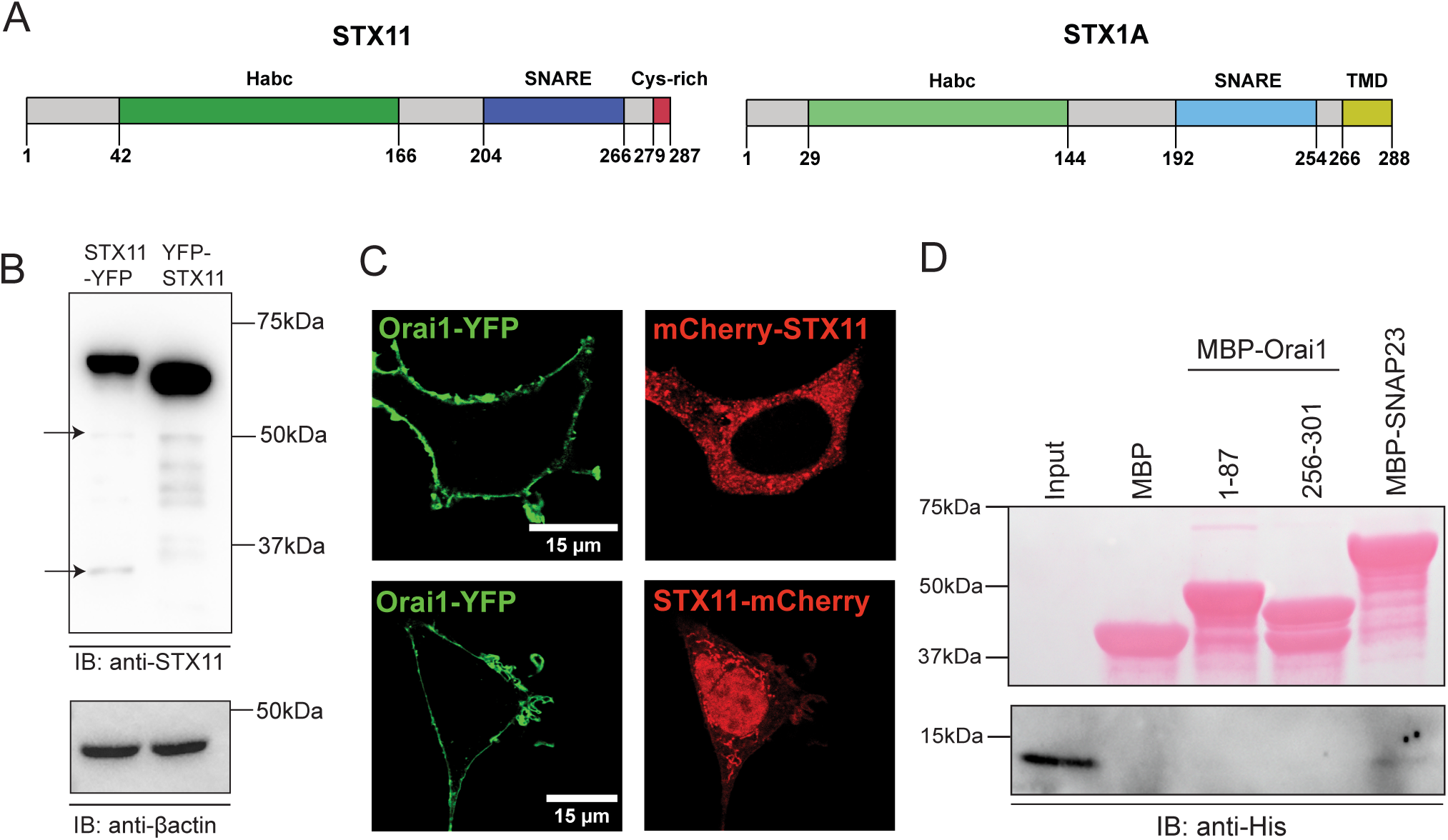
**(A)** Schematic showing a comparison of the key domains in STX11 and STX1A. **(B)** Western blot of whole cell lysates prepared from HEK293 cells expressing YFP-STX11 and STX11-YFP. Note the mismatch in molecular weight of the N-versus C-terminally tagged STX11 and the degradation products (arrows). N=3. **(C)** Confocal images showing localization of ectopically expressed mCherry-STX11 or STX-11-mCherry in HEK293 cells. N=3. **(D)** Pull-down assay showing *in vitro* binding of His-tagged SNARE domain of STX11 to MBP-tagged Orai1 N- and C-termini. (Top panel) Ponceau S staining showing the input of MBP alone or MBP-tagged Orai1 cytosolic tails. (Bottom panel) Western blot using anti-His antibody.

**Figure 5-figure supplement 2.**
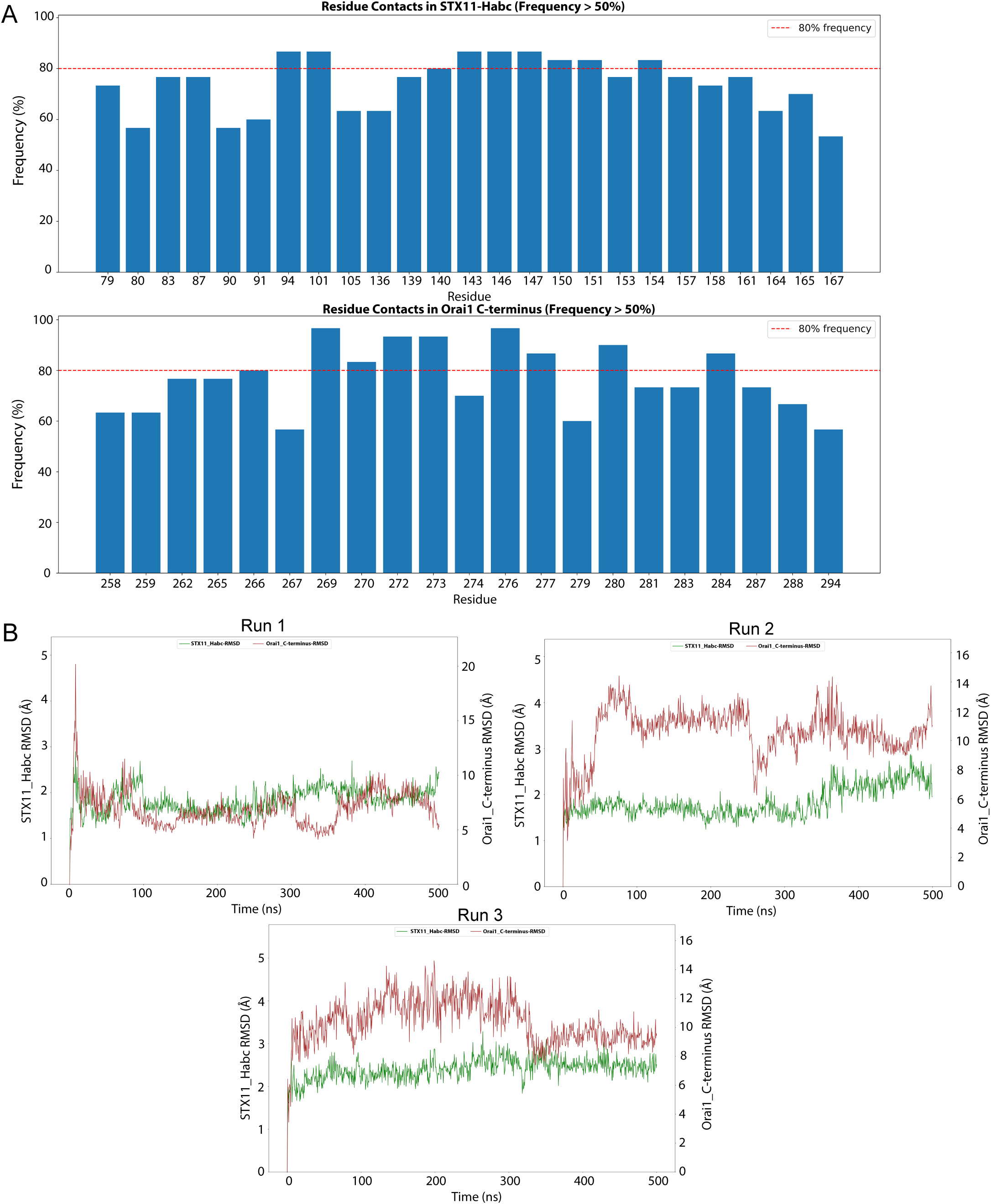
**(A)** Frequency of interface residues of STX11-H_abc_ and Orai1 C-terminus across AF3 predicted models. Residues with more than 50% frequency are shown. **(B)** RMSD of STX11 H_abc_ and Orai1 C-terminus upon superposition on the 0th ns frame. The STX11-H_abc_ RMSD indicates the backbone RMSD upon superposition of all frames on the STX11-H_abc_ backbone of 0th ns. The Orai1 C-terminus RMSD suggests the stability of Orai1 with respect to STX11 and is calculated for the Orai1 C-terminus after superposition of all frames on the STX11-H_abc_ backbone of 0th ns.

**Figure 5-figure supplement 3.**
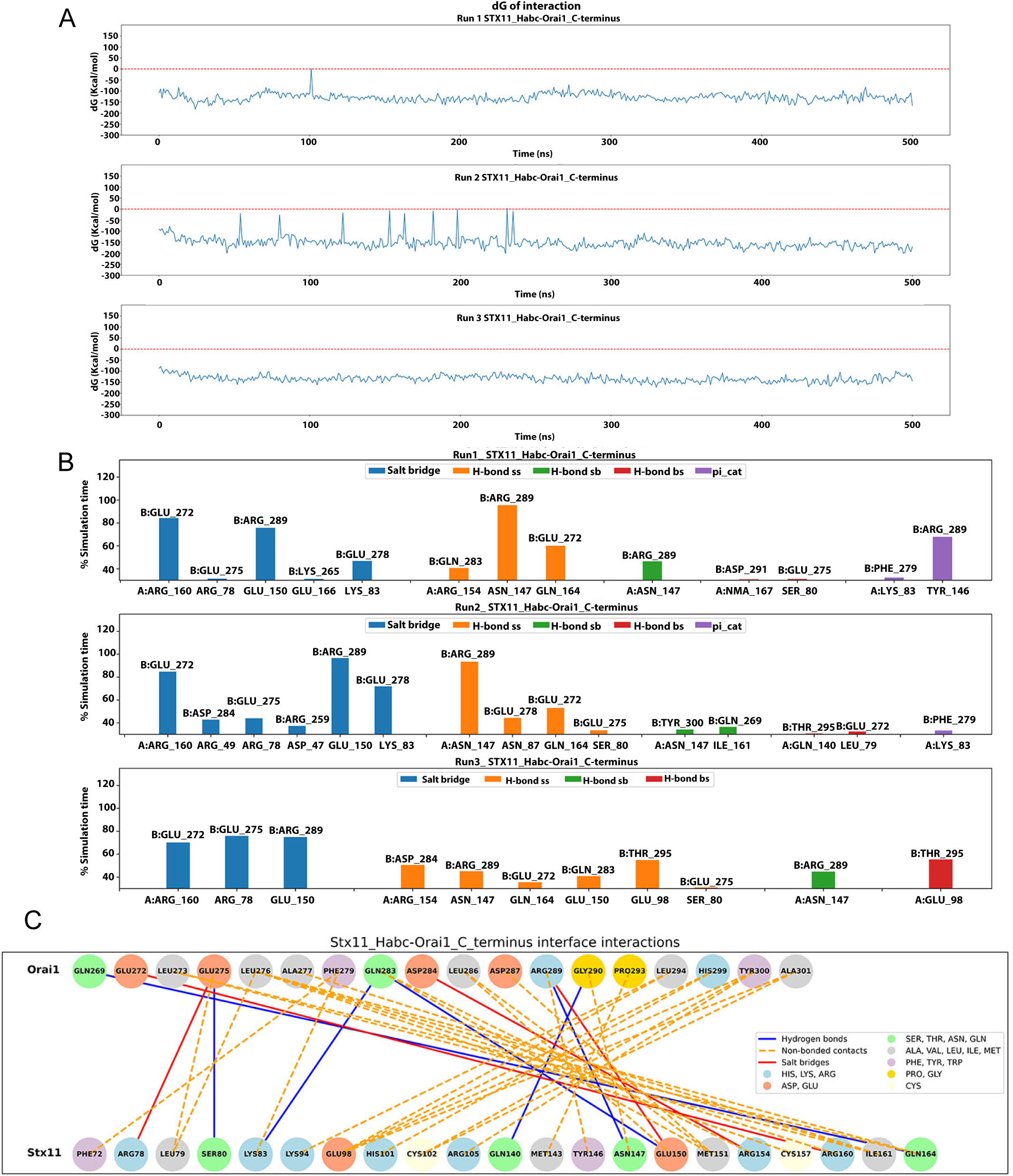
**(A)** Binding energy (ΔG) of STX11-H_abc_ and Orai1 C-terminus complex across the simulation time for all three replicates. **(B)** Interactions between STX11-H_abc_ and Orai1 C-terminus across the simulation time in three replicates. The interactions that persist for more than 30% of simulation time are shown and highlighted in different colors. H-bond has been categorized as sidechain-sidechain (ss), sidechain-backbone (sb) and backbone-sidechain (bs). **(C)** Residues predicted to be involved in interaction between STX11 H_abc_ and the Orai1 C-terminus identified from all-atom MD simulations.

**Figure 5-figure supplement 4.**
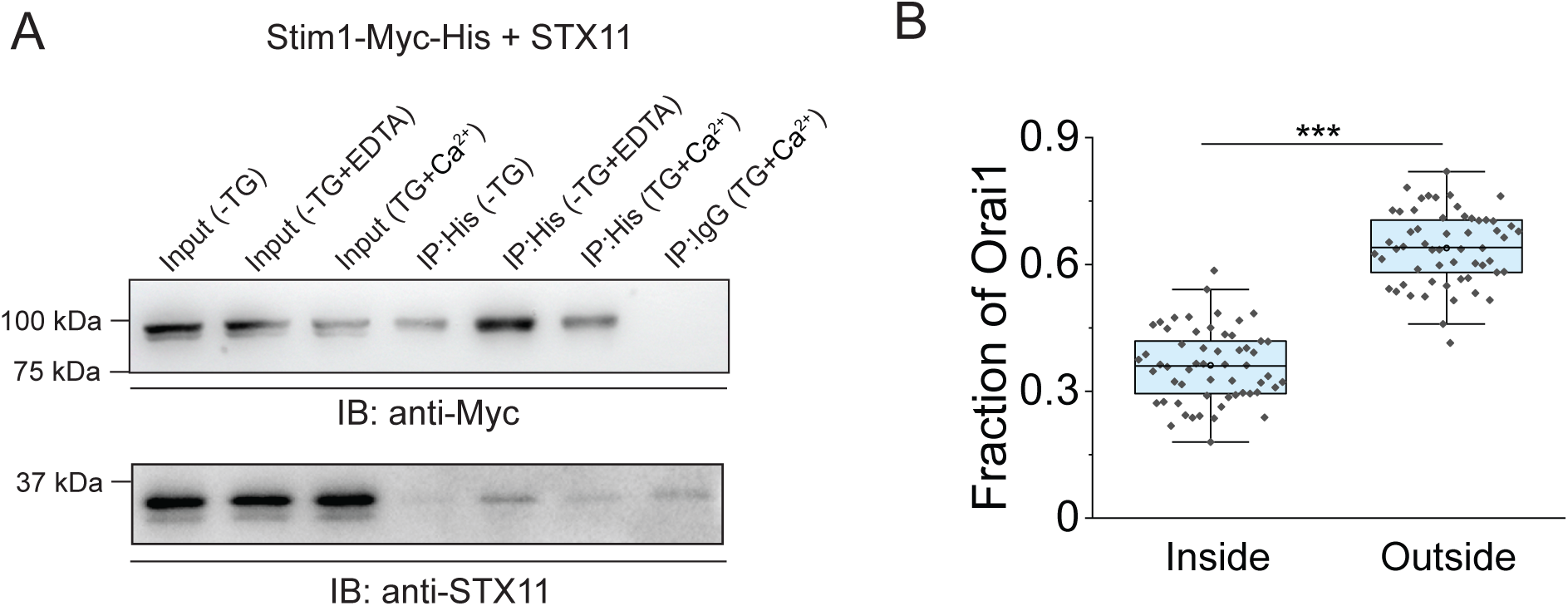
**(A)** Co-immunoprecipitation (co-IP) to test the association of Stim1 with STX11. HEK293 cells were co-transfected with Stim1-Myc-His and STX11, store-depleted, lysed and subjected to co-IP followed by Western blot as indicated. (N=2) **(B)** Quantification of fraction of Orai1 inside and outside Stim1:Orai1 clusters in store-depleted HEK293 cells expressing CFP-Orai1 and Stim1-YFP.

**Figure 5-figure supplement 5.**
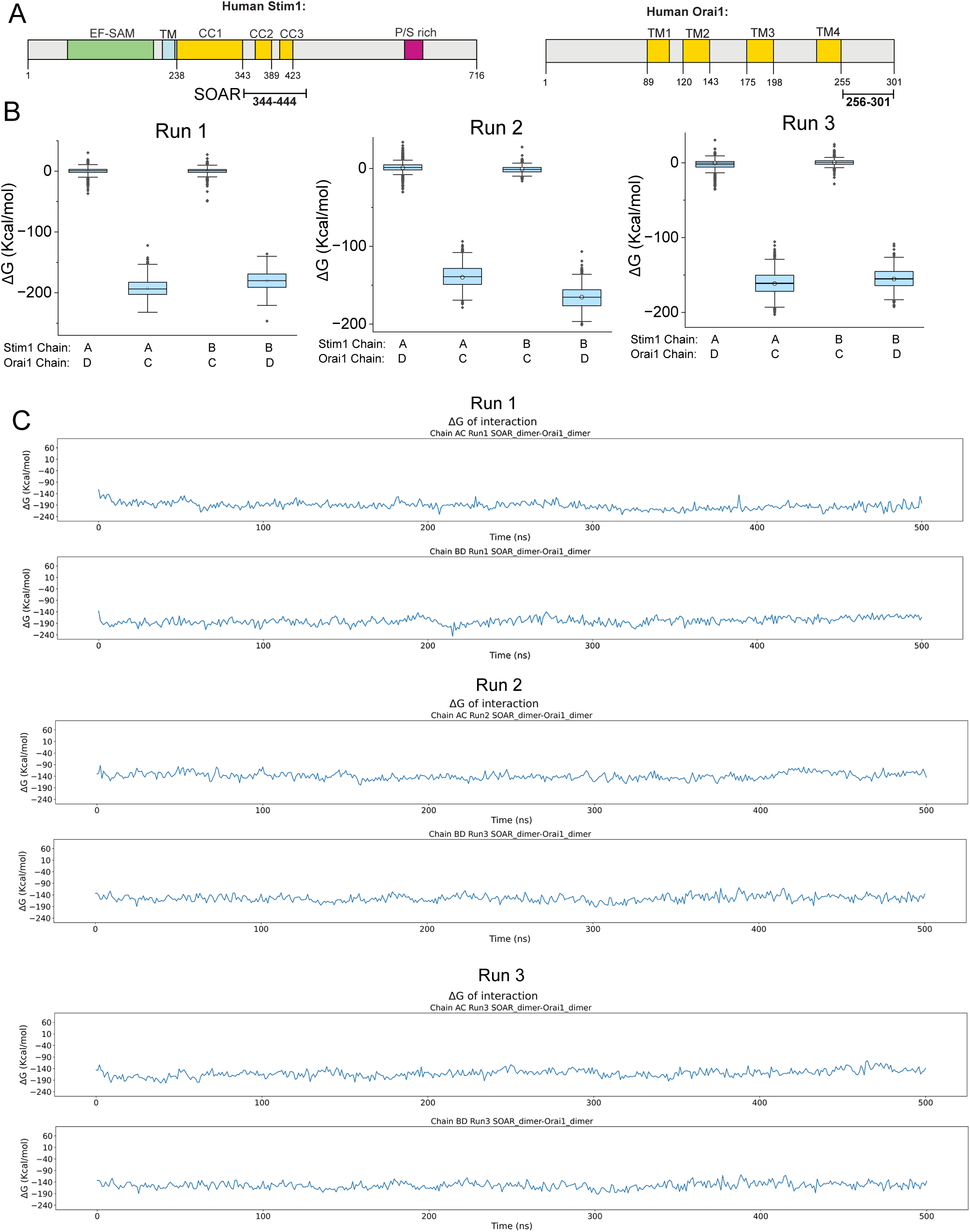
**(A)** Schematic of human Stim1 and Orai1 showing the domains used for MD simulation. TM:Transmembrane, P/S: Proline/Serine rich and CC: represent Coiled coil domains. **(B)** The binding free energy distribution of the three runs for the interactions between the SOAR dimer and Orai1 C termini. **(C)** Binding free energy (ΔG) of SOAR dimer and Orai1 C-termini complex across the simulation time for all three runs.

**Figure 5-figure supplement 6.**
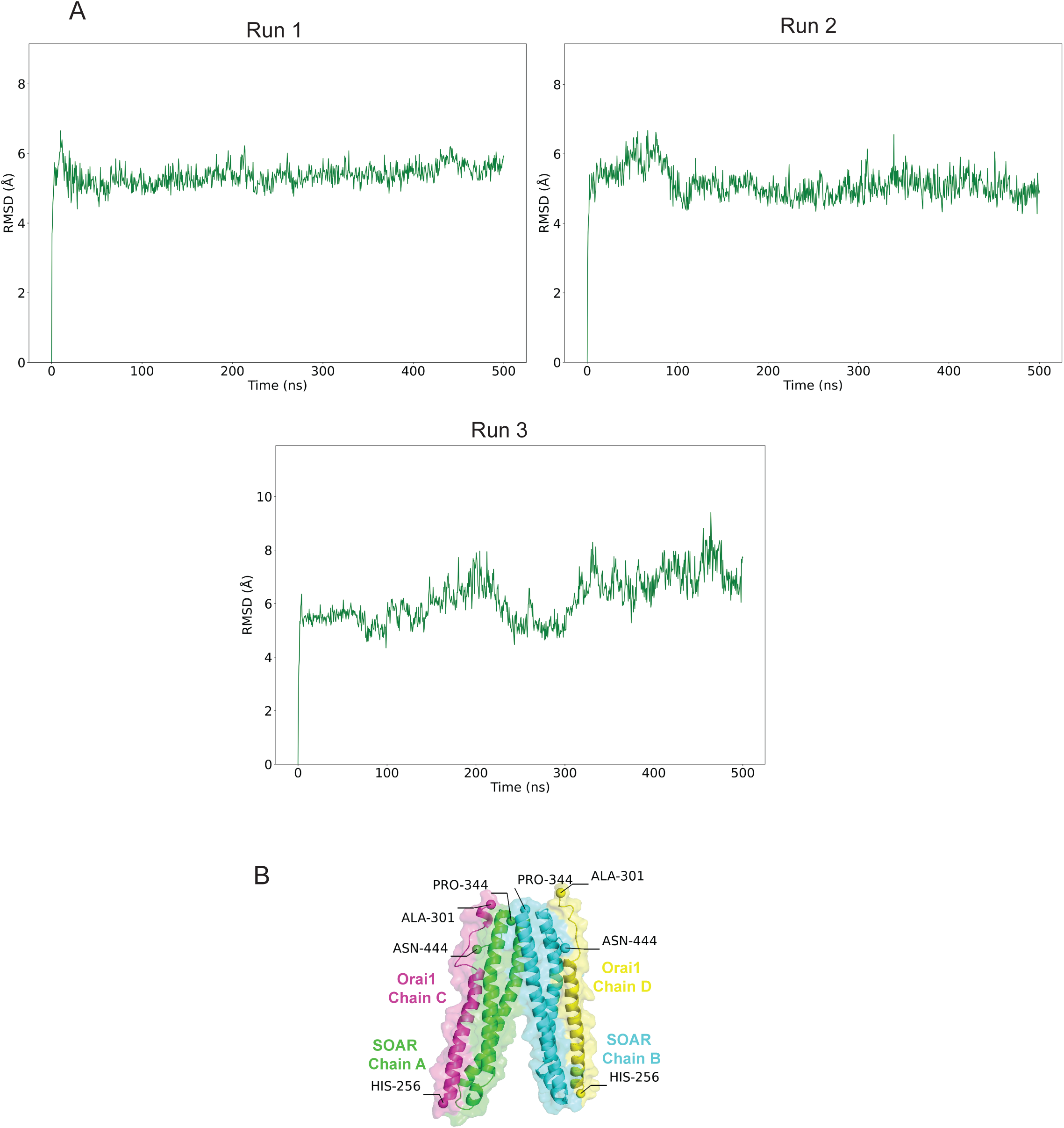
**(A)** RMSD of SOAR dimer and Orai1 C-termini complex. **(B)** Cartoon representation of the structure of Stim1-SOAR(344-444) dimer in complex with Orai1 C-termini.

**Figure 7-figure supplement 1.**
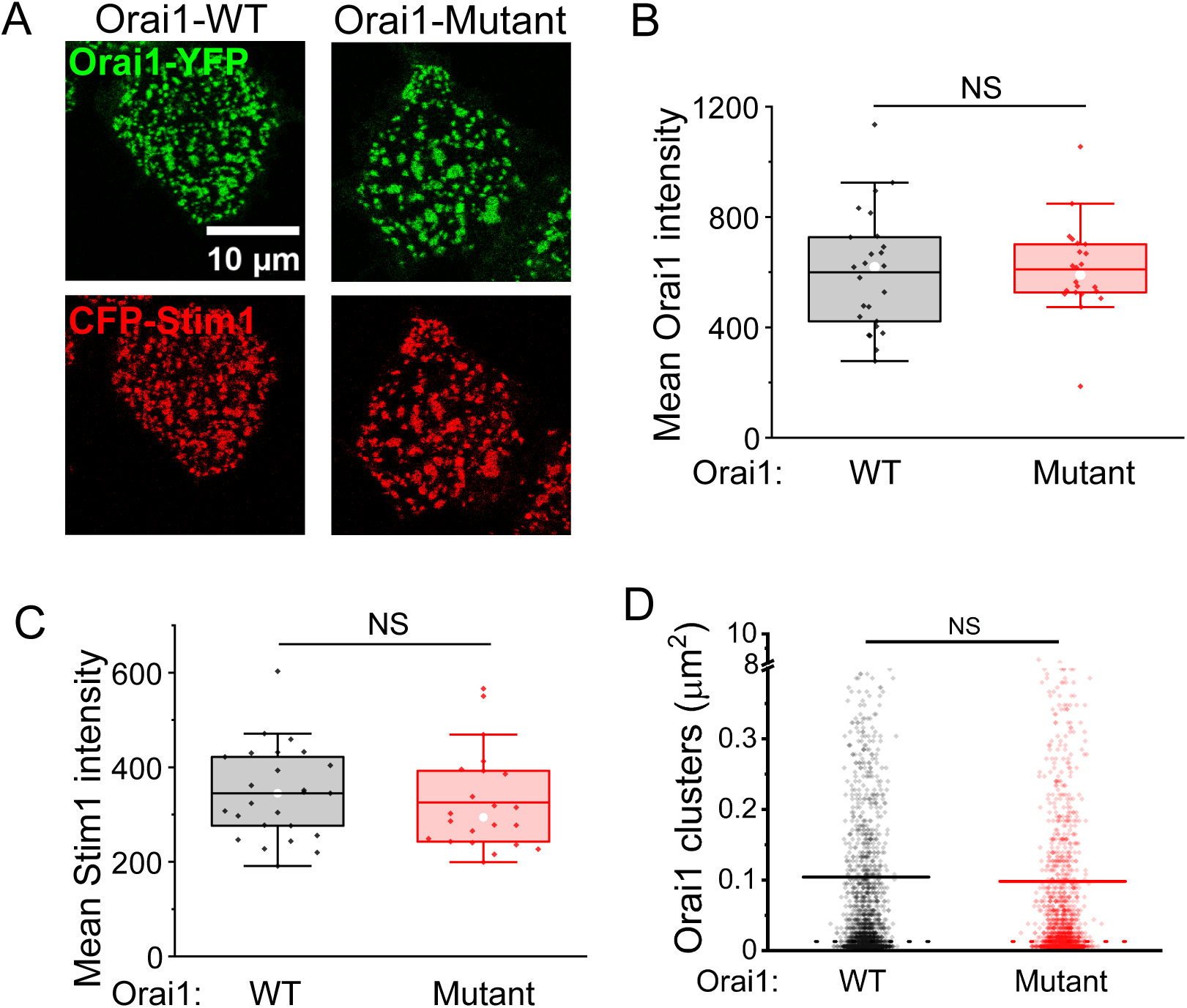
**(A-D)** Representative confocal images **(A)** and quantification **(B-D)** of YFP-tagged wild-type or R289A_E272A_E275A-E278A mutant Orai1 and CFP-Stim1 intensities and area of clusters in store-depleted HEK293 cells. n=20.

**Figure 8-figure supplement 1.**
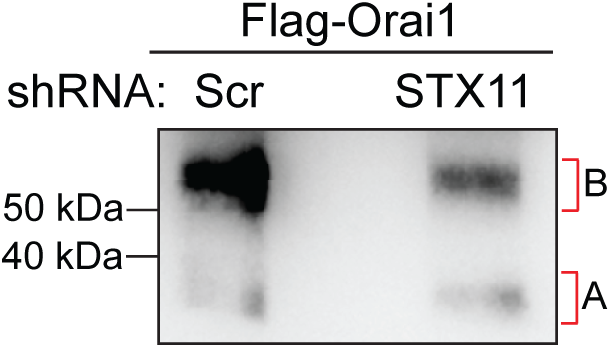
Lower half of the Western blot of BS3 cross-linked Flag-Orai1 acquired at a higher exposure showing the monomeric Flag-Orai1 band (labelled as A)

**Figure 8-figure supplement 2.**
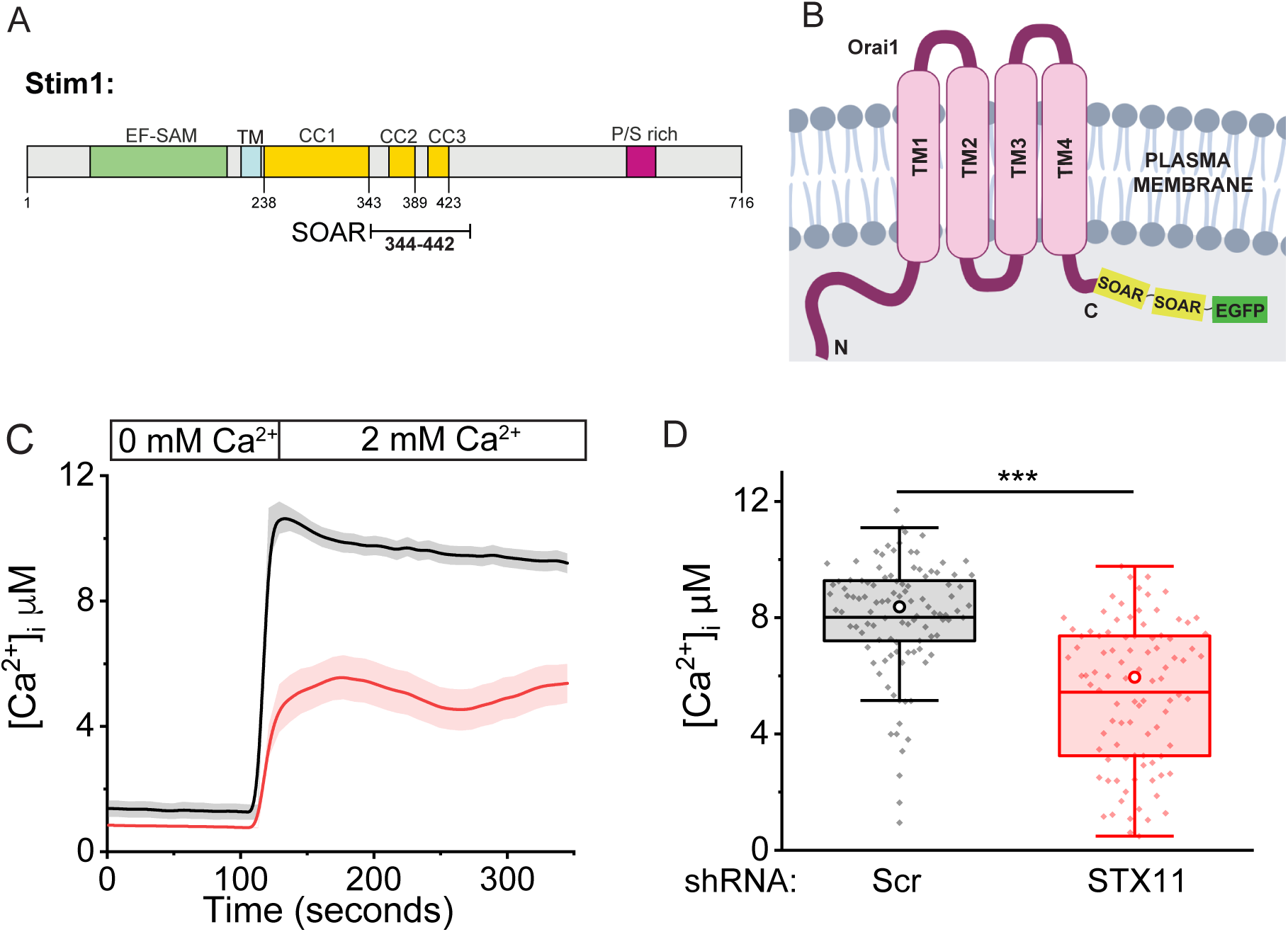
**(A)** Schematic representation of SOAR domain in full length Stim1. **(B)** Cartoon showing the design of the Orai1-SOAR-SOAR-EGFP (Orai1-S-S-GFP) construct. **(C)** Representative Fura-2 calcium imaging assay to measure constitutive calcium influx in Orai1-S-S-GFP expressing control (black) or STX11 depleted (red) HEK293 cells. Cells were imaged in Ringer’s buffer containing 0 mM followed by 2 mM extracellular Ca^2+^. **(D)** Quantification of constitutive calcium influx across experiments as shown in **(C)**. n= 80-90, N=3.

**Figure 8-figure supplementary 3.**
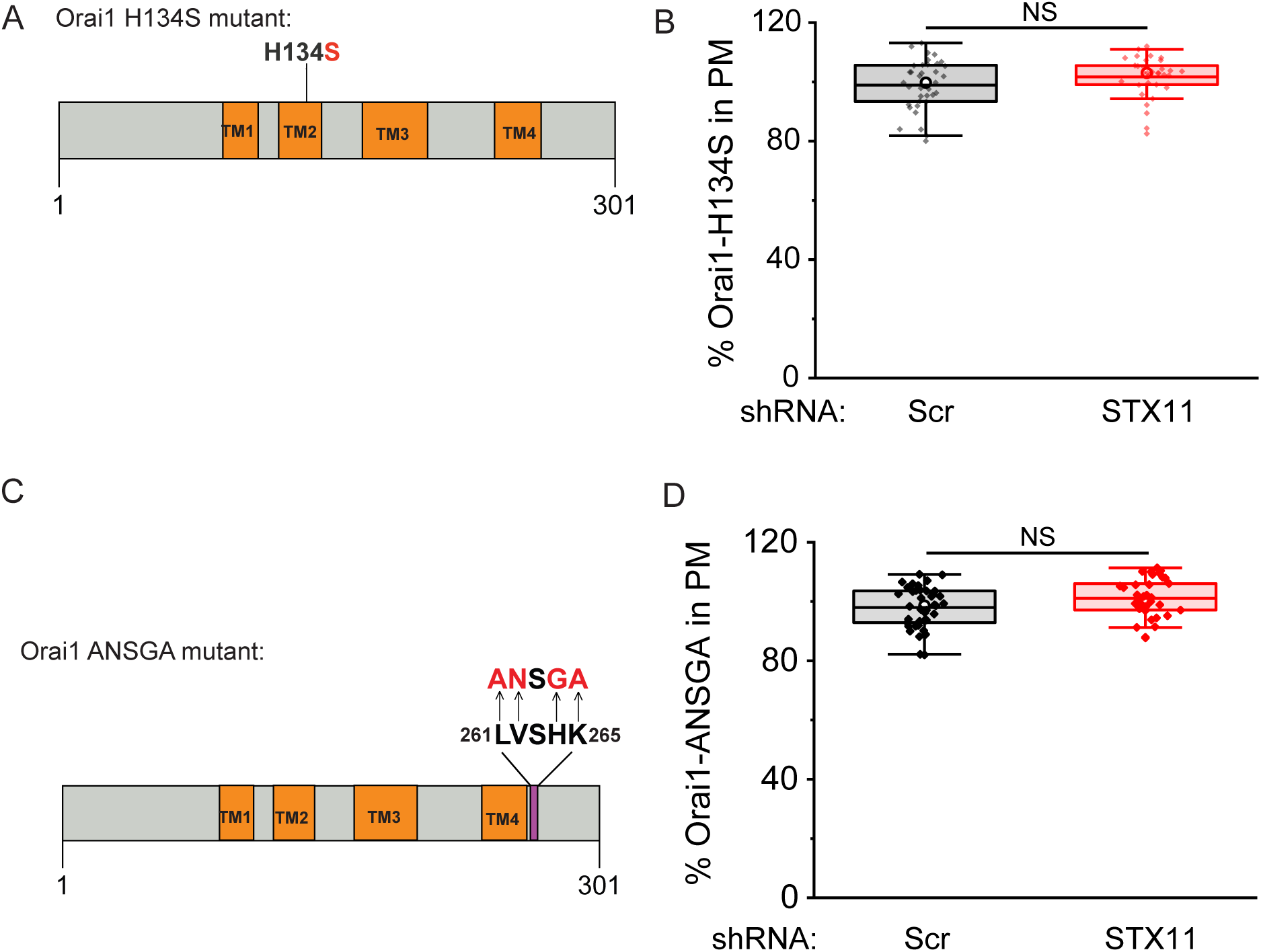
**(A)** Schematic representation of H134S mutation in full length Orai1. **(B)** Quantification showing percent Orai1-H134S-YFP intensity normalized to Orai1-WT-YFP in the PM of Scr and STX11 shRNA treated HEK293 cells. **(C)** Schematic representation of ANSGA mutation in full length Orai1. **(D)** Quantification showing percent Orai1-ANSGA-YFP intensity normalized to Orai1-WT-YFP in the PM of Scr and STX11 shRNA treated HEK293 cells.

